# The Pentose Phosphate Pathway Regulates Myelo-Lymphoid Lineage Specification

**DOI:** 10.64898/2026.09.07.747966

**Authors:** Jason Cosgrove, Alessandro Donada, Anne-Marie Lyne, Noemie Karabacz, Ildefonso Rodriguez-Ramiro, Adil Midoun, Vincent Cabeli, Cecile Conrad, Gregoire Jouault, Adeline Durand, Sabrina Tenreira-Bento, Emilie Tubeuf, Helen Demollin, Erica Russo, Fanny Tabarin, Yannis Belloucif, Shayda Maleki-Toyserkani, Sophie Reed, Federica Monaco, Lucas Ruffinatto, Berengère De Laval, Ann Ager, Camille Lobry, Philippe Bousso, Pablo Jose Fernández-Marcos, Céline Vallot, Herve Isambert, Nina Cabezas-Wallscheid, Rafael J. Argüello, Leïla Perié

## Abstract

Following infection, hematopoietic stem and progenitor cells (HSPCs) support immunity by increasing the rate of innate immune cell production but the metabolic cues that guide this process are unknown. To address this question, we combined *in situ* RNA barcoding and metabolomics approaches to perform metabolic state-fate mapping *in vivo*. This approach revealed a subset of myeloid-biased HSPCs that express a distinct set of metabolic enzymes and transporters as well as the surface marker CD62L. Metabolically, CD62L^high^ HSPCs have differential activity of the pentose phosphate pathway (PPP), OXPHOS and translation, as well as differential levels of S-adenosylmethionine (SAM) cycle metabolites associated with epigenetic modifications. Inhibition of the PPP skews HSPC lineage fate decisions by disrupting myeloid associated enhancers, while simultaneously increasing enhancer activity for the master B-lymphoid regulator Ikaros. *In vivo*, overexpression of glucose-6-phosphate dehydrogenase – a rate limiting enzyme of the PPP - skewed HSPC output from B-lymphocytes. In summary, our data shows that HSPCs undergo significant metabolic changes to facilitate the bioenergetic and epigenetic demands of myeloid versus lymphoid lineage specification. We highlight a key role for the pentose phosphate pathway which modulates myeloid rather than lymphoid commitment by shaping the HSPC enhancer landscape, providing proof of principle that HSPC metabolism can be targeted to modulate immune system dynamics.

## Introduction

Throughout the body, stem cells need to constantly adapt the amount and type of cells that they produce to maintain tissue homeostasis, compensate for cell loss and promote tissue repair. A key challenge in the stem cell field is to understand the molecular signals that selectively differentiate stem cells into specialized cell types *in vivo*. Much emphasis has been placed on the role of transcription and growth factors in guiding lineage specification, but the role of metabolism in this context is less established.

Under homeostasis, hematopoiesis is the dominant biosynthetic process in the human body. It generates 10^11^ cells each day^1,2^ and accounts for ∼86% of daily cell turnover^1^. Following infection, hematopoietic stem and progenitor cells (HSPCs) rapidly change the number and the types of immune cells produced^1,2,3^. HSPCs (Lin^-^ Sca1^+^ cKit^+^) are a rare and functionally diverse cell type, comprising both quiescent hematopoietic stem cells (HSCs) and downstream multipotent progenitors (MPPs). Fate-mapping and cellular barcoding studies have shown that the MPP compartment acts as the major source of new cells^4–6^, and is where lineage branchpoints occur ^7,8^. Thus, MPPs play an active role in immune responses, constantly adjusting the type and number of cells that they produce in response to context-specific signals from the BM microenvironment or peripheral tissues^9,10,11^. To date, much emphasis is placed on identifying and dissecting the networks of cytokines, transcription factors, and niche-forming stromal cells that promote lineage specification in MPPs, while metabolic processes were historically considered as passive house-keeping functions necessary for cell division and maintenance.

While cellular metabolism has been shown as a key regulator of HSC dormancy, self-renewal and erythroid specification^12–22^, relatively little is known about the metabolic regulation of myeloid versus lymphoid specification^23,24^. Currently, it is unclear which metabolic cues shape the type and amount of immune cells that MPPs produce *in vivo,* in steady state and in inflammation. Metabolites could act as signalling molecules^25^ and epigenetic modulators^26,27^ in addition to their roles in bioenergetics and biosynthesis. Metabolic pathways provide the substrates required for the epigenetic modifications preceding RNA expression changes. They also provide the cofactors regulating epigenetic enzymes. Little is known about how metabolic and epigenetic crosstalk regulates hematopoietic lineage specification.

Technical challenges are associated with measuring metabolic processes in rare cell types such as HSPCs. Metabolites, typically short-lived - in the order of minutes^28^ - display a large structural diversity. State-of-the-art mass-spectrometry-based assays need at least 10^4^ cells as input^20,29^. Such limitation in sensitivity make it difficult to link metabolite measurements of bulk populations to the functional heterogeneity of individual HSPCs^7,8,30,31^. In addition, most metabolomics approaches require cell permeabilization or lysis, preventing inferences on the cellular outcome *in vivo.* The lack of functional information from existing metabolomics approaches is critical, critical, with recent studies showing that -omics profiling should be paired with functional measurements to resolve HSPC heterogeneity^32,33^.

Lineage specification causes significant changes in cell size, macromolecular composition, and gene expression. It is thus an energy consuming process, challenging the notion of a passive housekeeping function. Here we hypothesise that metabolism regulates HSPC fate by providing the epigenetic and biosynthetic substrates required for myeloid versus lymphoid lineage specification. To test this hypothesis, we perform metabolic state-fate mapping of HSPCs *in vivo* through the integration of *in situ* RNA barcoding and metabolomics approaches. Using this fate-resolved multi-omics approach, we identify a subset of CD62L-expressing MPPs that are the major source of myeloid cells in steady state and emergency myelopoiesis and express a distinct set of metabolic enzymes and transporters. Metabolomics and metabolic profiling assays corroborated this data, revealing that CD62L^high^ myeloid-biased MPPs have a higher rate of pentose phosphate pathway (PPP), OXPHOS dependence and translation levels relative to other HSPCs. In addition, CD62L^high^ MPPs exhibit increased levels of associated methylation-based epigenetic modifications metabolites from the S-adenosylmethionine (SAM) cycle, suggesting that metabolic and epigenetic regulation are coupled to orchestrate myeloid lineage commitment. Specifically in CD62L^high^ MPPs, PPP inhibition resulted in an increased of H3K4Me1 marks at the enhancer of the Ikaros gene, a master regulator of B-lymphocyte lineage development^34^. *In vivo,* overexpression of the glucose-6-phosphate dehydrogenase enzyme - rate limiting enzyme of the pentose phosphate pathway - limited the production of B-cells from transplanted MPPs, skewing output towards the myeloid lineages.

Overall, our data shows that the pentose phosphate pathway shapes the HSPC commitment to myelopoiesis over lymphopoiesis. Our data also provide proof of principle that HSPC metabolism can be targeted to modulate the magnitude and specificity of immune cell regeneration *in vivo*.

## Results

### Metabolic State-Fate Mapping Identifies Enzyme and Transporter Markers of Myelopoiesis

In this study, we hypothesized that metabolism regulates the type and number of immune cells that HSPCs produce *in vivo* by fueling the epigenetic and biosynthetic demands of myeloid lineage specification. To address this question, we used *in-situ* barcoding combined with single-cell transcriptomics to identify metabolic regulators of myeloid specification *in vivo*. Although transcriptomics cannot be used to directly infer metabolite levels, pairing expression patterns of enzymes and transporters (metabolic state) with cellular barcoding (cell fate) allows functional and phenotypic information for a single cell, which cannot be achieved using existing metabolomics methods.

To this end, we used our DRAG (**D**iversity through **RAG**) *in situ* barcoding technology, allowing inducible labelling of cells in their native environment^35^ (**Figure 1A, Figure S1A)**. We extended the DRAG system to recover barcode transcripts from 10X genomics 3’ scRNAseq libraries using a targeted amplification of the invariant region of DRAG barcodes (**Figure S1B-E**). We injected RosaCreERT2^+/-^ DRAG^+/-^ mice with tamoxifen, inducing barcode recombination at 8-11 weeks of age (**Figure 1A, Figure S1A)** and waited 47-67 weeks post-induction to track only barcodes originating from HSPCs^35^. Then, we isolated barcoded HSPCs (Sca1^+^ cKit^+^ GFP^+^), myeloid (Cd11b^+^ GFP^+^) and erythroid (Ter119^+^ CD44^+^ GFP^+^) cells from the bone marrow of 5 mice, using fluorescence-activated cell sorting (FACS) (**Figure 1A; Figure S2**). We next processed HSPCs for single-cell RNA sequencing and targeted barcode amplification (materials and methods). In nucleated erythroid and mature myeloid cells, we detected barcodes from cell populations at the DNA level^35^. We developed a custom bioinformatics pipeline to analyse the sequencing results, and to deal with the analytical challenges of metabolic state-fate mapping (*supplementary information*). Following data QC and preprocessing (**Figure S3-S4**, *detailed in materials and methods),* we recovered RNA barcode information for 668 hematopoietic stem and progenitor cells (14.9% recovery rate at the RNA level), corresponding to 158 unique lineage barcodes, 97 of which were also detected in mature cell subsets (**Table S4**). Analysing the barcodes distribution across the myeloid and erythroid lineages, we observed that HSPCs are highly heterogeneous in the number (**Figure 1B**) and type (**Figure 1C**) of cells that they produced.

**Figure 1.**
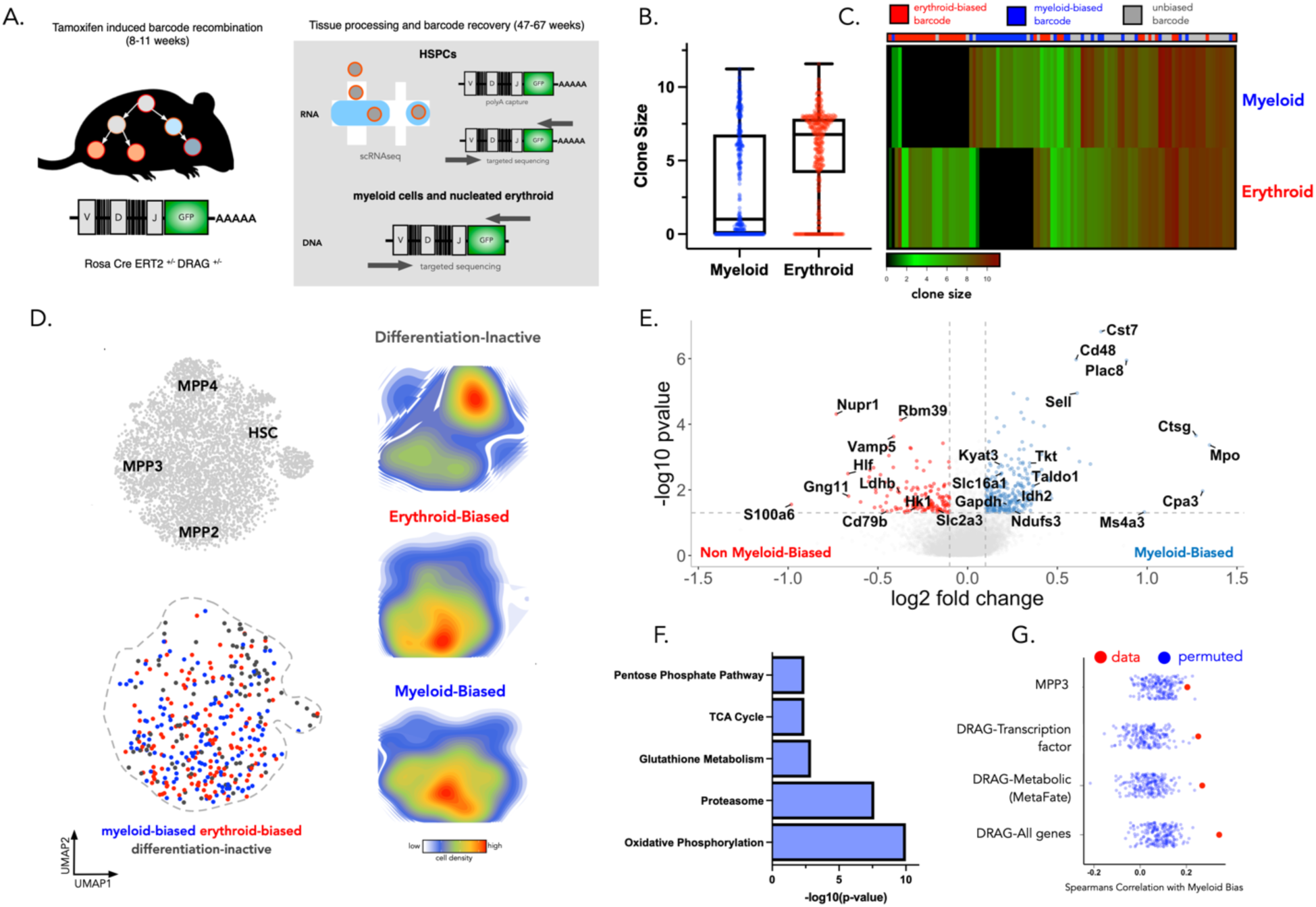
Myeloid-biased HSPCs have a distinct expression program of enzymes and transporters that are predictive of cell fate. (A) Overview of the fate-resolved transcriptomics approach: Our pipeline begins with the induction of a lineage barcode in single cells in situ, then the propagation of barcodes in vivo when cells divide and differentiate and finally then the recovery of the transcriptomes of HSPC and their barcode from RNA as well as the recovery of the barcode of their mature progeny by DNA. (B) Clone sizes (number of cells per barcode) in the erythroid and myeloid lineage, the y-axis is transformed using the hyperbolic arcsin function. Each point represents a single barcode. (C) Heatmap representation of DNA barcode expression in myeloid and erythroid cells. Normalized and hyperbolic arcsin transformed cell counts (clone size) data were clustered by hierarchical clustering using Euclidian distance. Color indicates hyperbolic arcsin transformed cell counts (clone size) The top column indicates differentiation active barcodes that are erythroid biased (red), myeloid-biased (blue) or unbiased (grey). (D) UMAP representation of the MetaFate dataset of LSK cells overlaying the positioning of known HSC and MPP subsets (top left) as well as the localisation of myeloid-biased (blue), erythroid-biased (red), and differentiation inactive (grey) progenitors based on lineage barcode (bottom left). This figure represents 4,485 Sca1^+^ cKit^+^ GFP^+^ cells (668 RNA-barcoded cells ; 158 unique barcodes). (right hand side) Density map highlighting the localization of lineage biased barcoded cells on our UMAP representation of the data. (E) Volcano plot showing differentially expressed genes between myeloid-biased barcoded cells and other (erythroid-biased and differentiation inactive) barcoded subsets. (F) Metabolic pathways from the KEGG database that are enriched amongst genes upregulated in myeloid biased barcoded progenitors compared to erythroid and differentiation inactive barcoded cell subsets. (G) Spearmans Correlation between different transcriptomic signatures and the myeloid bias of lineage barcodes. Red points represent the correlation between signature scores and myeloid bias score, while blue points represent the correlations observed for randomised gene-sets of an equivalent size. The MPP3 signature is taken from Sommerkamp et al (2021). To create the DRAG gene signatures we identified 271 genes upregulated in myeloid biased clones (DRAG - all genes). The subset of genes from the DRAG signature relating to cellular metabolism form the DRAG metabolism signature, while the subset of genes encoding transcription factors form the DRAG transcription factor signature. The signature score corresponds to the average expression values of these gene sets for each cell and is projected onto the UMAP visualisation of the data. In this figure all data was taken from 5 mice from 3 independent experiments.

To characterize the functional heterogeneity of HSPCs, barcode labelled HSPCs were classified as differentiation inactive (98 cells ; 61 unique barcodes) if we could not detect their barcode in any mature cell compartments^31,32,36,37^, or as erythroid-biased (143 cells ; 35 barcodes), myeloid-biased (143 cells ; 31 barcodes), or unbiased (284 cells ; 31 unique barcodes) depending on the relative abundance of the barcode across the respective lineages (**Figure 1C-D).** Sampling was not affecting these results, as bulk replicates displayed high correlation (**Figure S3a**) and barcode clone sizes were comparable across all classes (**Figure S3b**). In addition, barcodes that had more than 75% of their barcode reads in the myeloid or erythroid lineage were classified as lineage-biased (**Figure 1C-D**), or otherwise classified as unbiased. Similar results were obtained with thresholds close to 75% (**Figure S4E-G**), while more extreme thresholds (90% or below 55%) impacted the number of barcodes and the magnitude of difference in gene expression, precluding robust analysis (**Figure S4E-G**). Many of barcodes had a low barcode generation probability indicating that they labelled a single cell rather than multiple cells, confirming the single-cell resolution of our approach (**Figure S1F-G**).

Within our UMAP representation of the data (**Figure 1D, Figure S4A**), we observed a significant overlap in the distribution of erythroid and myeloid-biased clones within MPP-associated regions of UMAP space. This was not resolved by unsupervised clustering on gene expression alone (**Figure S4D**) or mapped to an existing known MPP subset (**Figure 1D**). Differential expression analysis between myeloid-biased, erythroid-biased and differentiation inactive clones identified a total of 271 genes which were upregulated in myeloid-biased clones, defining the DRAG-Fate myeloid gene signature (**Figure 1E, Table S5-7**). Among these genes were existing markers of myeloid potential including *Mpo, Ctsg, Ms4a3* and *Cpa3*^32,38,39^. Interestingly, 57/271 genes within the DRAG-Fate myeloid signature encoded enzymes and transporters from metabolic pathways of the KEGG, REACTOME and GO databases (**Figure 1F**). This subset of metabolically-associated genes, hereafter called the MetaFate signature, comprises genes relating to OXPHOS (*Idh3a, Cox7b, Ndufa4, Uqcr10*), proteostasis and ribosome biogenesis (*Hdc, Kyat3, Sec61b,Slc35b1,Psmc4*), the pentose phosphate pathway (*Tkt, Taldo1, Gpi1, Pgls*) as well as genes relating to the regulation of redox state (*Idh2, Gsto1, Mgst2, Txn2,Txndc11,Gpx1*) and epigenetic modifications (*Wdr5, Pcgf6, Satb1, Baz1a, Aebp2*) **(Figure 1E).**

Within HSPCs, expression of the DRAGFate and MetaFate gene signatures for only metabolic genes and for the whole transcriptome are highly correlated (**Figure S5A**), suggesting that enzyme/transporter expression state alone could be used to predict myeloid fate in HSPCs. To compare the predictive power of metabolic-associated genes from the MetaFate myeloid signature against other signatures, we computed the Spearman’s correlation coefficient between gene signature expression scores and the myeloid bias score of HSPCs derived from lineage barcoding information (**Figure 1G**). In this analysis the DRAG fate signature consistently outperformed the MetaFate signature, suggesting that metabolism is not the only program contributing to myeloid bias (**Figure 1G**). However the MetaFate signature predicted myeloid bias to a similar extent than gene sets relating to transcription factor activity (**Figure 1G**), which are established regulators of fate choice, highlighting the importance of metabolic regulation in cell fate decisions. Furthermore, in comparison to known signatures of myeloid bias in HSPCs, the MetaFate signature had a higher correlation (rho = 0.24, p-value = 2.5 x 10^-10^) with myeloid bias compared to the existing MPP3 signature^39^ (289 genes) (rho = 0.2, p-value = 1 x 10^-7^) (**Figure 1G**). DRAG-barcode derived signatures also outperformed the MPP3 signature of myeloid bias when we performed 4-fold cross-validation analysis, to assess the sensitivity of our result to overfitting (**Figure S4H**), revealing the power of combined barcoding and transcriptome in the same cells to identify lineage bias subsets. To assess the broader predictive power of the MetaFate signature, we quantified its expression across 3 scRNAseq datasets of hematopoietic progenitors^40–42^. In these datasets, the MetaFate signature was upregulated in myeloid progenitors relative to other progenitor subsets (**Figure S5B**). Taken together these analyses showed that the MetaFate signature, comprising only genes associated with metabolism, was a robust predictor of the myeloid output of MPPs in native hematopoiesis.

In summary, by combining RNA expression and fate analysis, we showed that enzyme and transporter expression patterns are associated with HSPC commitment to myeloid lineage *in vivo*. It suggested that metabolic rewiring accompanies the earliest stages of myeloid development.

### Fate-Resolved Transcriptomics Identifies CD62L as a Marker of Metabolically Distinct Myeloid-Biased HSPCs

As fate-resolved transcriptomics identified a novel myeloid-biased MPP subset with a distinct expression program of enzymes and transporters, we developed a purification strategy to isolate this subset and further assess their functional and metabolic properties. Differential expression analysis between barcoded HSPCs (**Figure 1G**) highlighted *Sell*, the gene encoding the adhesion molecule CD62L, as a putative marker of cells expressing the MetaFate expression program (**Figure 1E**). We also observed significant differences in *Sell* between MetaFate^low^ and MetaFate^high^ cells (cells below and above the 75^th^ percentile of MetaFate signature expression respectively, p < 0.001) (**Figure 2A**). Flow cytometry profiling showed that CD62L is heterogeneously expressed in HSPCs (**Figure 2B**), with high expression in a subset of MPP3 and MPP4 cells, and little to no expression in LT-HSCs, ST-HSCs and MPP2s (**Figure S6A**), consistent with our MetaFate analyses. Based on these results, we selected CD62L as a marker of MetaFate^high^ myeloid-biased MPPs.

**Figure 2:**
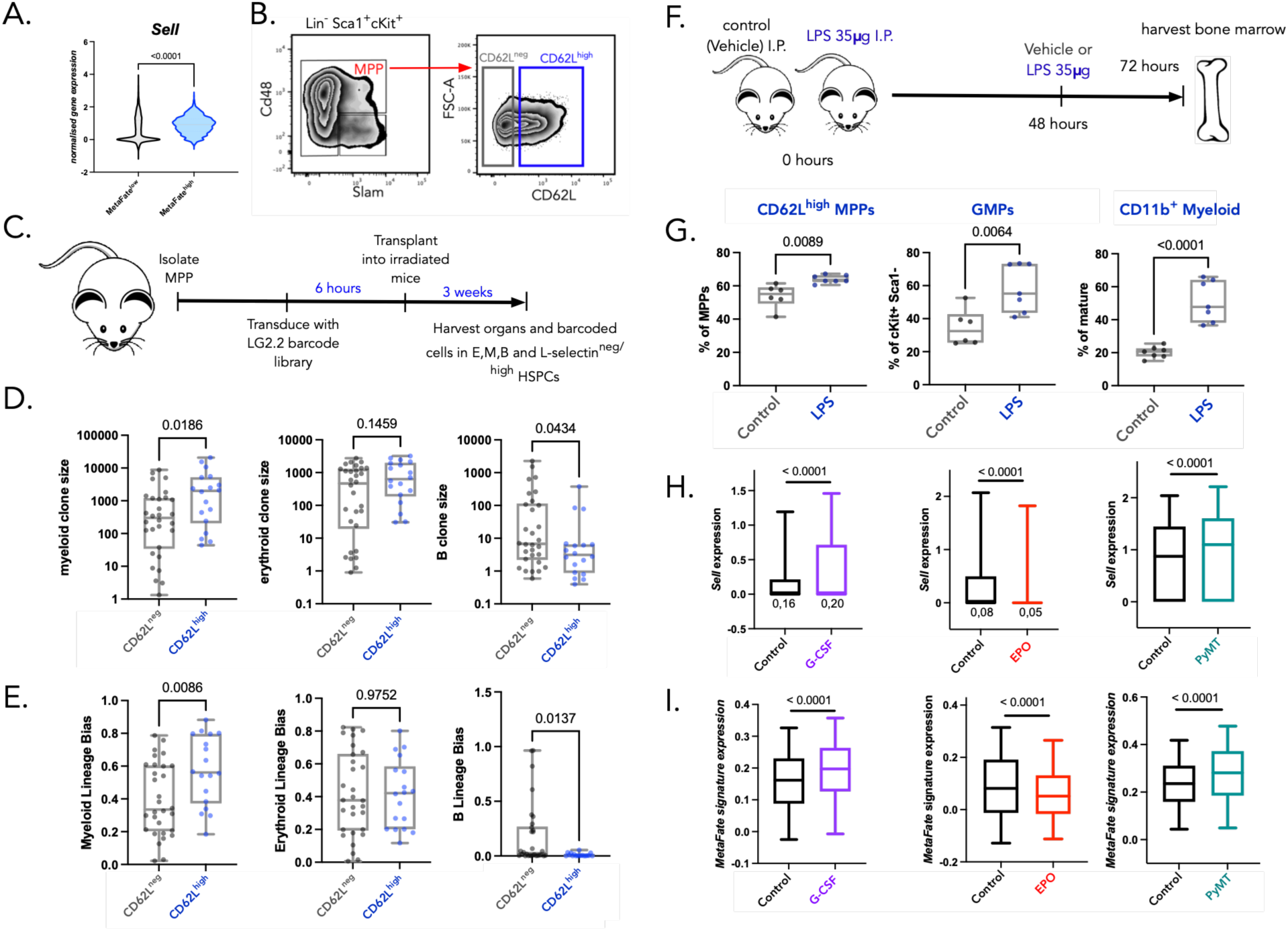
CD62L is a marker of myeloid bias and differential expression of metabolic enzymes and transporters in steady state, and emergency myelopoiesis. (A) Comparison of *Sell* expression between MetaFate^low/high^ expressing populations. MetaFate-low cells are defined as HSPCs in the bottom 25^th^ percentile of MetaFate signature expression. MetaFate-high cells are defined as HSPCs in the top 75^th^ percentile of MetaFate signature expression (B) Flow cytometry gating strategy to purify L-selectin expressing MPPs (C) Overview of lentiviral barcoding analysis. MPPs were purified from male donor B6 WT mice by FACs and infected with the LG2.2 lentiviral barcoding library for 6 hours to achieve a multiplicity of infection of 10%. Transduced cells were transplanted into 4 sublethally irradiated (6Gy) control recipient mice (male littermate controls) and 3 weeks later CD62L^neg/hi^ MPPs, CD19^+^ B cells, CD44^+^ Ter119^+^ erythrocytes, and CD11b^+^ myeloid cells, as well as CD62L^high^ or ^neg^ MPPs from the bone marrow were sorted and processed for targeted sequencing of lentiviral lineage barcodes. At the 3 week timepoint we observe 10% chimerism of GFP+ barcoded cells (Figure S8a-b). (D) Clone sizes (number of cells of the myeloid, erythroid and B-cell lineages) produced per barcode for the CD62L^neg^ (grey) and CD62L^high^ (blue) MPP subsets. Each point represents a distinct barcode. Statistical comparisons were made using a Mann-Whitney test. Each point represents a unique barcode. (E) lineage bias value for the myeloid, erythroid and B-cell lineages for barcodes in CD62L^neg^ (grey) and CD62L^high^ (blue) MPP subsets. Lineage bias represents the relative frequency of each barcode across the 3 mature cell lineage. Each point represents a distinct barcode. Statistical comparisons were made using a Mann-Whitney test. All Boxplots represent the median and interquartile range with whiskers extending to the minimum and maximum values. N= 4 mice. (F) Overview of the LPS challenge model. 18 week old B6j Mice were injected with LPS (35ug/mouse) I.P. at 0 hours and 48 hours. At 72 hours bone marrow cells were harvested and analysed by flow cytometry. (G) Quantification of the percentage of the different cell subsets in control (grey) and LPS treated (blue) mice. Normality of the data was assessed using a Shapiro-Wilk test and significance was assessed using a T-test. N = 13 mice and data was pooled from 2 independent experiments each. (H) Differences in Sell RNA and (I) MetaFate gene signature expression from single cell RNAseq in HSPCs in different cytokine stimulation (G-CSF stimulation of HSPCs ex vivo (Fast et al 2022); HSPCs taken from EPO stimulated mice (Tusi et al 2021)) and inflammation contexts (PyMT Breast cancer model - Gerber et al 2023). Boxplots represent the median and interquartile range and whiskers extend to the 5^th^ and 95^th^ percentiles each point represents a different mouse for panel D,E,G. As the difference in Sell expression for the G-CSF and EPO group compared to control in the panel H are difficult to see, we have added the value of the mean for each group in the figure.

To independently validate CD62L as a marker of myeloid bias HSPCs, we purified CD62L^high^ and CD62L^neg^ MPPs by FACS and transplanted them into irradiated recipient mice. However the use of the MEL-14 anti-CD62L antibody clone to sort cells and transplantation led to poor engraftment of CD62L^high^ MPPs (**Figure S6B**). This is in line with the inhibition of CD62L function by MEL-14 on leukocytes^43^. To overcome this limitation, we lentivirally barcoded total MPPs and transplanted them into irradiated recipient mice (**Figure 2C, Figure S7A**). Barcodes present in the CD62L^neg^ and CD62L^high^ MPPs, as well as the nucleated erythroid, B cells and myeloid cells (172 barcodes in total) were analysed 3 week post-transplantation – the timepoint when myeloid production from MPPs peaks post-transplantation^44^ (**Figure 2C**). Following data QC and preprocessing (**Figure S7B-F,S8A-C, Table S8**), we focused on the fate outcome of the barcodes that had more than 95% of their reads in either the CD62L^neg^ (30 barcodes) or the CD62L^high^ (18 barcodes) MPPs (**Figure S8D**). We found that CD62L^high^ barcodes produced more myeloid cells and less B cells, compared to CD62L^neg^ barcodes, while erythroid production was similar (**Figure 2D-E**), showing that CD62L ^high^ MPPs were myeloid biased. Importantly, we detected the same total number of unique barcodes for both the CD62L^high^ and the CD62L^neg^ subsets (**Figure S8C**). This suggests that lineage biases are not affected by sampling, sensitivity issues or differences due to the engraftment rates. Furthermore, we observed similar patterns in myeloid bias when we transplanted lentivirally barcoded CD150^+^ HSCs, and purified CD62L MPP subsets at 12 months post transplantation (**Figure S9-S10, Table S9**). Collectively, both *in situ* and lentiviral barcoding confirmed that CD62L marks HSPCs biased toward myeloid rather than lymphoid output *in vivo*.

### MetaFate and *Sell* Expression Are Modulated by Growth Factor and Inflammatory Signalling

Following infection, HSPCs dramatically increase the amount of myeloid cells they produce^9,10,11^. We therefore hypothesised that the CD62L^high^ Metafate^high^ MPPs are responding to growth factor and inflammatory signals associated with emergency myelopoiesis. To assess the role of CD62L^high^ MPPs in infection, we first used an LPS challenge model, in which TLR4 signalling in endothelial cells leads to G-CSF mediated stimulation of myelopoiesis^10^. Mice were given 35μg of LPS i.p. at 0 and 48 hours, and bone marrow samples were processed for flow cytometry analysis at 72 hours^45^ (**Figure 2F**). In this model of emergency myelopoiesis^45^, we observed an increased proportion of CD62L^high^ MPPs (**Figure 2G**) which correlated with increases in both the Granulocyte-Monocyte Progenitor (GMP) (cKit^+^ Sca1^-^ CD16/32^+^ CD34^-^) and myeloid (Cd11b^+^) compartments of the bone marrow (**Figure 2G**). The increased proportion of CD62L^high^ MPPs was not associated with an increased CD62L expression induced by LPS, as shown both *ex vivo* and *in vivo* (**Figure S11**). Similarly, with G-CSF^46^, another inflammatory cytokine stimulating myelopoiesis, *ex vivo* stimulated HSPCS had increased *Sell* and MetaFate gene expression. Conversely, with the EPO, a cytokine stimulating red blood cell production, *in vivo* stimulation^40^ led to decreased *Sell* and MetaFate expression (**Figure 2H-I**). These results were corroborated in the context of a live infection by analysing a scRNAseq dataset of cKit^+^ progenitors from WT mice, or mice infected with *Plasmodium Berghei* 7 days post infection^47^ (**Figure S12**) – an established model of emergency myelopoiesis driven by interferon mediated signals. In this setting, MetaFate and Sell expression program gene signature were significantly increased in MPPs from infected mice (**Figure S12**).

A number of chronic disorders including cancer are associated with changes in HSPC function leading to the increased production of myeloid cells. To assess whether *Sell* and MetaFate expression are altered by tumor-associated inflammation, we analysed the transcriptomes of HSPCs taken from the bone marrow of C57BL/6 MMTV-PyMT (PyMT) and associated control mice. In this established non-metastatic model of spontaneous breast cancer, increased levels of myelopoiesis are driven by an expansion of myeloid biased HSPCs in response to tumor-derived inflammatory signals^48^. In this context, *Sell* and the MetaFate expression program were significantly increased relative to MPPs taken from tumor bearing mice compared to control mice (**Figure 2H-I**), corroborating the data from both LPS, cytokine, and infection challenge experiments. While in all these emergency myelopoiesis models the classical MPP subpopulations varied in proportion, this variation was specific to each model (**Figure S13**). In contrast, the increased expression of MetaFate^pos^ CD62L^high^ MPPs was consistent across models (**Figure S13**).

In summary, infection and inflammation leading to increased demand for new myeloid cells induce expansion of the CD62L^high^ HSPC compartment in the bone marrow and increased expression of the MetaFate program of enzymes and transporters. This demonstrates that the CD62L^high^ myeloid-primed HSPCs with specific associated enzyme and transporters program are active players in a variety of infection and inflammation settings, most probably to respond the increased demand of myeloid cells.

### Metabolic Adaptations Enable HSPCs to Meet the Biosynthetic Demands of Myeloid Commitment

Functional and transcriptomics analyses showed that CD62L-expressing HSPCs express a distinct program of metabolic enzymes and transporters that are predictive of myeloid lineage fate. This program included activation of genes related to the PPP, OXPHOS and redox metabolism. To assess the metabolic activity of these pathways in CD62L^neg/high^ MPP subsets in comparison to other MPPs, we used two complementary fluorescence based assays (**Figure 3A**): SCENITH (Single Cell Metabolism by Profiling Translation inhibition) and SPICE-Met. SCENITH is a flow cytometry-based method^49^ based on profiling protein synthesis rates in response to metabolic inhibitors using flow cytometry. SPICE-Met provides a measure of cellular ATP:ADP ratio using a genetically encoded PercevalHR biosensor, that can be measured using fluorescence microscopy or flow cytometry^50^. PercevalHR is composed of a mutated version of the ATP-binding bacterial protein GlnK1 and the circular permuted monomeric Venus fluorescent protein. ATP but not ADP binding to the PercevalHR causes a ratiometric shift in the probe fluorescence excitation spectrum, providing a read out of ATP:ADP intracellular ratio. In these two metabolic profiling assays, cells are purified from the bone marrow of wild-type CD57BL/6j (SCENITH) or Vav-iCre Perceval^fl/fl^ (SPICE-Met) mice, then treated with either DMSO (control; Co), or small molecule inhibitors of glycolysis (2-deoxy-D-glucose; 2-DG), OXPHOS (Oligomycin; O) and protein synthesis (harringtonine; H) for < 1 hour *ex vivo*. We then compared the fluorescent intensities of ATP, ADP and puromycin incorporation rates across different experimental conditions and used this information to calculate the dependencies of ATP:ADP and translation on the activity of different metabolic pathways.

**Figure 3.**
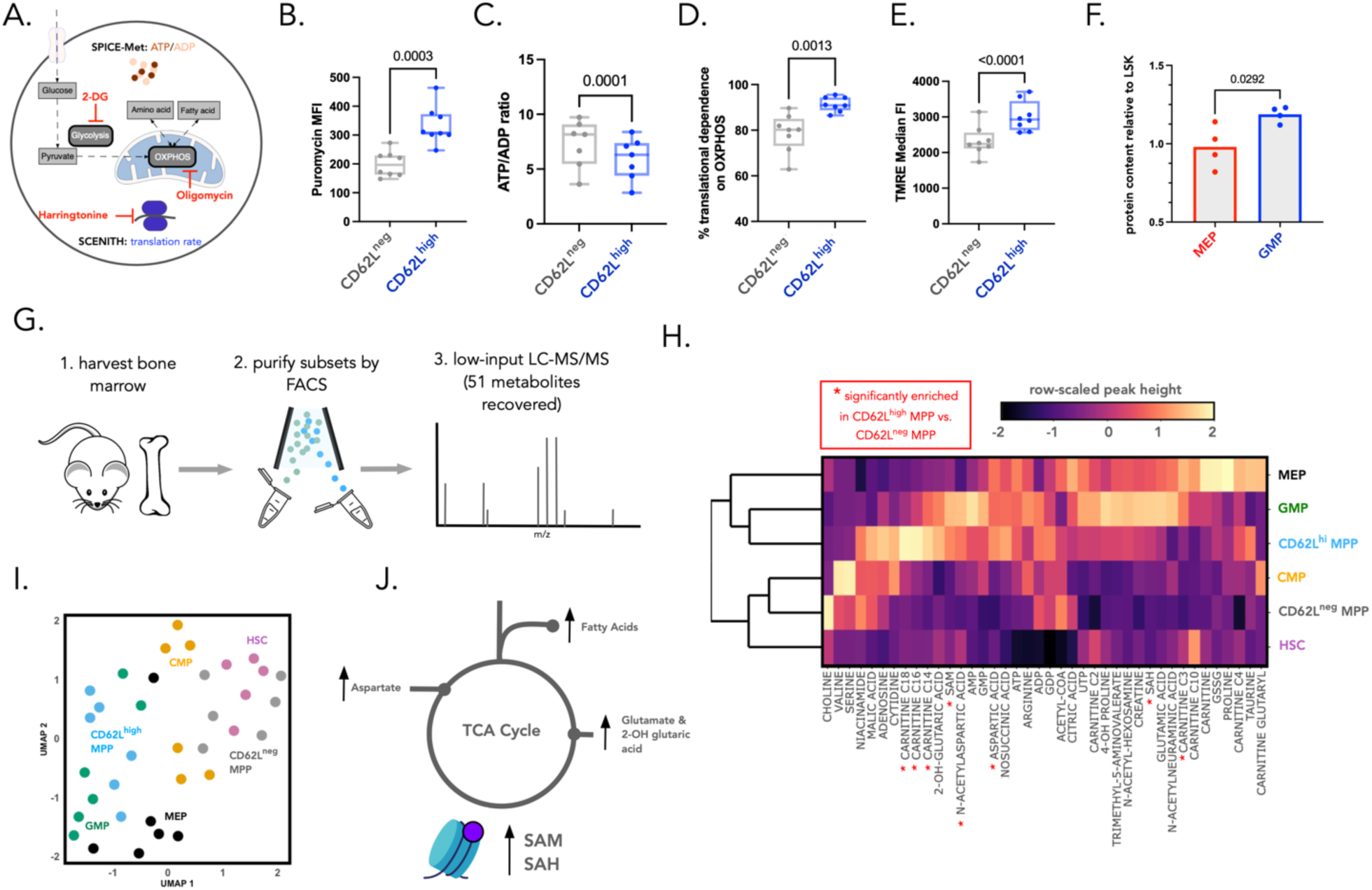
CD62L^high^ Multipotent Progenitors Are Metabolically Distinct from other Progenitor Subsets: (A) Overview of our strategy to profile the metabolic state of different HSPC subsets. Cells are incubated in the presence of DMSO (control) or inhibitors of glycolysis (2-DG), OXPHOS (Oligomycin) or translation (Harringtonine). Following incubation of cells with these inhibitors, we measure either translation rate (puromycin labelling), and or the percevalHR biosensor as a measure of ATP:ADP ratio. (B) Median fluorescence intensity values for puromycin across CD62L MPP subsets. Each point represents 1 mouse, data pooled from 2 independent experiments (N = 8 mice). (C) ATP:ADP measurements obtained by dividing the median fluorescent intensity of the ATP channel by the median fluorescent intensity of ADP channel. Each point represents one mouse (L) Comparison of ATP:ADP ratio in CD62L^neg^ and CD62L^hi^ MPP control samples (D) Comparison of mitochondrial dependence measures calculated based on changes in puromycin labelling across control, oligomycin and harringtonine treated conditions. Formula for data transformation is provided in the materials and methods (E) Median fluorescent intensity measures for tetramethylrhodamine, ethyl ester (TMRE) labelling. N = 8 mice, data pooled from 2 experiments. Each point represents 1 mouse. Statistical comparisons were made using a paired T-test (F) Relative protein content of MEP and GMP compared to LSK as determined using the microBCA assay. Data were reanalysed from Hidalgo et al. (2020).(G) Overview of the experimental design. Cell suspensions were prepared from the femur, tibia and iliac crest bones taken from 6-8 week old male B6j mice (n = 6 mice pooled from 2 independent experiments). Cell subsets (HSC, CD62L^neg/high^ MPP, GMP, MEP, CMP) were then purified by FACS and processed for low input LC-MS/MS profiling. (H) Heatmap showing the median peak height value across all mice, normalised per metabolite for 40 metabolites that were detected across both independent experiments. Clustering was performed using complete linkage of the Euclidean distances between cell types (I) UMAP projection of the metabolomics data where colors represent cell type and each point represents a biological replicate. Each point represents one mouse (n = 6). (J) Figure summarising differences in metabolites that are enriched in CD62L^high^ MPPs compared to CD62L^neg^ MPPs. Normality of the metabolomics data was assessed using a Shapiro-Wilk test and statistical comparisons comparing the CD62L^neg/high^ subsets were made using a paired T-test or a Wilcoxon signed rank test, depending on whether the data were normally distributed.

To assess whether SCENITH and SPICE-Met could be applied to study HSPCs, we first benchmarked them by comparing the metabolic profiles of HSCs (Lin^-^ cKit^+^ Cd48^-^ Slam^+^) and MPPs, for which a number of metabolic differences have already been reported^15,51,52^. Consistent with previous reports using targeted genetics, mitochondrial labelling dyes, and *in vivo* puromycin labelling ^15,51,52^, SCENITH and SPICE-Met profiling showed that HSCs had a higher glycolytic capacity and lower protein synthesis rate than MPPs (**Figure S14**), confirming that our approach can be successfully applied to study other hematopoietic progenitor subsets.

After validating the use of SCENITH and SPICE-Met on HSPCs, we then compared the metabolic <u>activity of</u> <u>glycolysis, OXPHOS and protein synthesis in</u> CD62L^neg^ and CD62L^high^ MPPs (Lin^-^ cKit^+^ Sca1^+^) (**Figure 3B-D**). SCENITH and SPICE-Met profiling showed that CD62L^high^ MPPs have a significantly higher rate of protein synthesis than CD62L^neg^ MPPs (p < 0.001) (**Figure 3B**) as well as a lower ATP/ADP ratio (p < 0.001) (**Figure 3C**) than CD62L^neg^ MPPs. The translation rates of CD62L^high^ MPPs were highly sensitive to oligomycin treatment (**Figure 3D**), suggesting an increased OXPHOS-dependence of translation relative to CD62L^neg^ MPPs (p = 0.001) (**Figure 3D**). Similar results were obtained when we measured the uptake of mitochondrial membrane potential TMRE dye in CD62L^high^ and CD62L^neg^ MPP subsets (p < 0.001) **(Figure 3E)**. While MPP2-4 displayed differences of translation rates and OXPHOS-dependent translation (**Figure S15A,C,E**), splitting these MPP populations on the basis of CD62L expression increased differences between MPP3 and 4 (**Figure S15B,D,F**). Therefore, we concluded that CD62L expression discriminates between metabolically distinct MPP populations. Collectively, this data showed that specification toward myeloid rather than lymphoid lineage incurs a biosynthetic (protein) and associated bioenergetic (ATP) cost which is fuelled by increased OXPHOS activity of CD62L^high^ MPPs. Reanalysing data from Hidalgo et al^53^ (**Figure 3F**), GMP, the first myeloid committed progenitors, had a significantly higher protein content than MEPs and HSPCs. This increase cannot be explained only by changes in cell volume, as GMP are smaller than MEP^53^. This supported a marked requirement for increased protein synthesis and energy production during the first myeloid specification steps.

To further enrich our metabolic profiling of CD62L-expressing HSPCs, we performed low-input liquid chromatography mass spectrometry (LC-MS/MS) optimised for the targeted detection of polar metabolites^54^ **(Figure 3G)**. This method permits relative quantification of TCA cycle intermediates as well as amino acids, fatty acids and associated derivatives^54^. Specifically, we purified 5000 HSCs, MPPs (CD62L^neg/high^) and erythromyeloid restricted potential progenitors (CMP, GMP, and MEP) for each biological replicate and identified a total of 56 metabolites. 40 of them were detected at a higher level than negative control of cell debris and sheath fluid in at least one cell type across 2 independent experiments (**Figure 3H, Figure S16**). UMAP and PCA visualisation (**Figure 3I, Figure S16D)** as well as unsupervised clustering (**Figure 3h**) of the data showed that CD62L^high^ MPPs are most similar to GMP in their metabolomes, consistent with our fate-resolved transcriptomics results. In total 8 metabolites were differentially enriched (adjusted p-value <= 0.05) when comparing CD62L^neg/high^ MPP subsets including the OXPHOS-associated metabolites carnitine-bound long-chain fatty acids, aspartic acid, and n-acetylaspartic acid (**Figure 3H,J**). These results are consistent with fate-resolved transcriptomics, SCENITH and SPICE-Met which also suggested increased OXPHOS activity in myeloid-biased progenitors. In addition, we also observed differences in the metabolites S-adenosylmethionine (SAM) and S-adenosylhomocysteine (SAH). Together with the overexpression of redox regulating gene (Idh2, *Gsto1, Mgst2, Txn2,Txndc11,Gpx1*), it suggests that CD62L^neg/high^ MPP subsets may exhibit differences in epigenetic modifications through modulation of NADP+/NADPH (**Figure 3J**).

In summary, metabolomics profiling validated our fate resolved transcriptomics data, confirming the presence of a metabolically and functionally distinct CD62L expressing HSPC subset. Integrating the transcriptomics and metabolomics results, we found that CD62L^neg^ MPPs represented a less committed MPP, resembling but not equivalent to HSC, with an increased ATP/ADP ratio, a reduced dependency on OXPHOS and giving higher chimerism in transplantation than CD62L^high^ MPPs. In contrast, CD62L^high^ MPPs had a specific metabolic program associated with early myeloid versus lymphoid lineage. This CD62L^high^ MPPs specific program consisted on one end in a higher rate of OXPHOS, lower ATP/ADP ratio and increased translation, fuelling the increased protein requirements of transitioning to a myeloid progenitors. On the other hand, CD62L^high^ MPPs differed from CD62L^neg^ MPPs by their PPP activity, redox regulation and epigenetic associated co-factors. Given these results, we next investigated the extent by which this metabolic program can control HSPC fate.

### The Myeloid-Lymphoid Enhancer Landscape is Shaped by the Pentose Phosphate Pathway

To identify which metabolic pathways were most likely to play a causal role in myeloid versus lymphoid development, we developed a systems biology approach that models metabolic pathway expression changes during myeloid lineage development. To this end, we integrated fate resolved transcriptomics, published single-cell and bulk level transcriptomics, and metabolomics datasets together (**Figure 4**). We first tested how enzyme and transporter expression changes over myeloid development using the trajectory inference PAGA algorithm^55^ to a published scRNAseq dataset of cKit^+^ progenitors^41^ (**Figure 4A**). Fatty acid, retinol, ascorbate and aldrate metabolism were decreased during myelopoiesis, while PPP, glutathione metabolism and OXPHOS were increased. The enzymes associated with the PPP were upregulated earlier along myelopoiesis and also heterogeneously across MPPs (**Figure 4A)**. Next, we applied the MIIC causal network reconstruction algorithm^56^ to enzyme and transporter gene expression data obtained from bulk RNA sequencing samples across the entire hematopoietic system^57^ (**Figure 4B**). MIIC network reconstruction identified a network of metabolic genes associated with myeloid biased lineage identity (**Figure 4C-D**). The PPP enzymes (*Taldo1, G6pdx*) were strongly associated with redox state (*Gsr, Mgst1, Mgst2*), glucose metabolism (*Hk2, Pkm*), NADPH-oxidase activity (*Ncf1*) and lipid metabolism (*Scarb1, Abcd1, Acer3*). *G6pdx and Taldo1* were causally linked with *Gsr* and *G6pdx* with *Mgst1*, suggesting that PPP impacted directly the redox activity. Recent reports suggested that the PPP directly influence epigenetic state^58^ and our own results supported this hypothesis.

**Figure 4.**
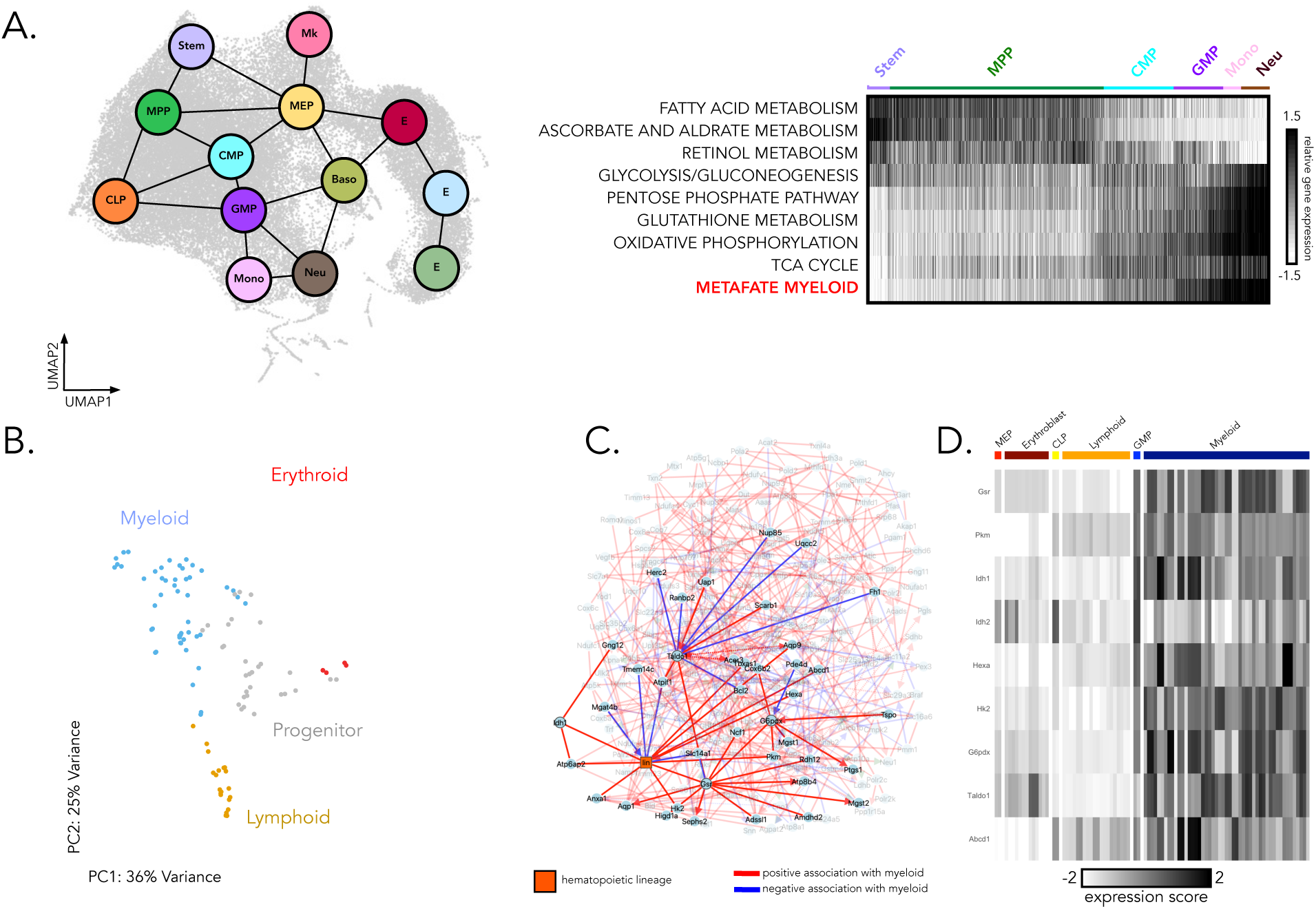
Systems Biology Modelling Identifies the Pentose Phosphate Pathway as a Core Regulatory Component of Myeloid Development and Cell State. (A left) A partition graph-based model of hematopoiesis generated on the scRNAseq dataset of cKit+ and cKit+ Sca1+ bone marrow cells (Dahlin et al (2018)) using the PAGA algorithm^55^. Preprocessing of the data was performed as described in Wolf et al.^55^. In this model each grey dot represents a single cell, each coloured circle represents a cell state (obtained by unsupervised clustering of gene-expression data), and edges between clusters represent putative transitions between cell states. (A right) Dynamics of metabolic pathway gene expression signatures along developmental trajectories inferred by the graph based differentiation model. Each vertical line represents a single cell and the color bar at the top represents the cell states (colored circles) depicted in panel A. Pathway gene-sets were taken from the KEGG database with the exception of MetaFate myeloid and signature scores were calculated as the mean expression value across all genes in the pathway minus the mean expression values taken from randomly sampled gene-sets of an equivalent size. (B) PCA plot of the Hemopedia Bulk RNAseq database^57^ using variably expressed metabolic enzymes and transporters. On this plot each point represents a bulk RNA-seq library of a FACs sorted hematopoietic cell types. (C) MIIC network generated on enzyme and transporters expression patterns in bulk RNAseq patterns from the Haemopedia database. To generate an input list of enzyme and transporter genes for miic analysis we next took genes from the myeloid metafate signature as well as genes encoding enzymes and transporters that were (i) upregulated in the myeloid lineage, erythroid or lymphoid lineages and (ii) variably expressed in LSK hematopoietic progenitors using published datasets^41,57^. A full list of these genes is provided in Table S10.Each circular node (blue) represents a gene. Hematopoietic cell lineage (erythroid, myeloid, and lymphoid) is represented by the square orange node and is based on which cell type the bulk RNAseq sample represents. Edges represent a predicted molecular association between genes or cell lineage with red edges highlighting genes that are positively associated with the myeloid lineage and blue lines representing genes that have a negative association with the myeloid lineage. (D) Expression patterns of enzymes/transporters that have a strong myeloid-association within the miic-generated network. Each column represents a bulk RNAseq sample taken from a FACS-sorted cell type defined in the haemopedia database.

Indeed the PPP was upregulated in the MetaFate signature of myeloid specification (**Figure 1F**) and the metabolomics and transcriptomics results indicated differences in epigenetic associated factors in CD62L^high^ MPPs. We thus decided to test if the PPP regulate myeloid lineage specification in MPPs by modulating their epigenetic state.

To confirm that PPP is was more active in the CD62L^high^ myeloid biased MPPs, we measured the protein levels of the PPP rate limiting enzyme Glucose-6-phosphate dehydrogenase (G6PD) by intracellular flow cytometry. We observed higher G6PD expression in CD62L^high^ MPPs compared to CD62L^neg^ MPPs **(Figure 5A).** The differential sensitivity of CD62L^neg/high^ MPPs to the PPP inhibitor 6-AN across a range of concentrations **(Figure S17**) corroborated the result that CD62L^high^ myeloid biased MPPs have differential PPP requirements.

**Figure 5.**
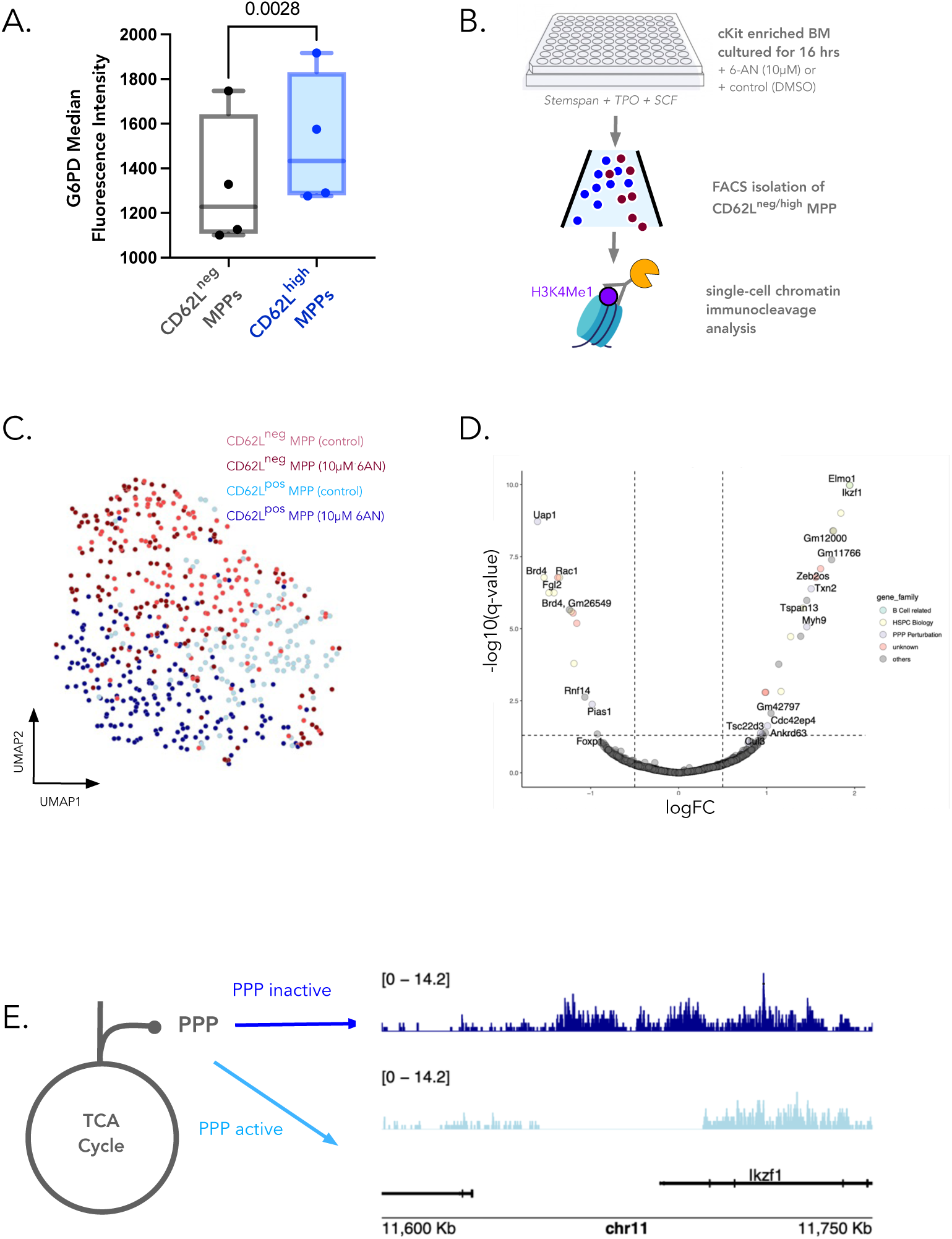
The Pentose Phosphate Pathway Shapes the Enhancer Landscape of CD62L^high^ MPPs. (A). G6PD protein expression measured in MPP subsets using intracellular flow cytometry. Each point represents a single mouse and statistical comparisons were made using a paired T-test. Normallity of the data was assessed using a Shapiro-Wilk test. (B) Overview of the scCHICseq experiment to assess H3K4me1 enhancer activity in MPPs following PPP inhibition with 10mm 6-AN. Data were pooled from 5 mice. (C) UMAP projection of H3K4me1 signal (blue represents CD62L^high^ and red represents CD62L^neg^, dark color shades represent 6-AN treatment and lighter shades represent DMSO control) (D) Differentially enriched enhancer regions in 6-AN treated versus control CD62L^high^ MPPs using the negative binomial method and defined by a q-value < 0.05 and an absolute log2 fold change > 0.5. Points highlighted in colours with different gene family. (E) Read distribution for H3K4me1 signal at the Ikzf1 locus for control and treated CD62L^high^ MPPs (H) Quantification of H3K4me1 signal at the Ikzf1 locus for control and treated CD62L^high^ MPPs. Read counts were normalised using the CPM method.

To test whether the PPP influenced the epigenetic state of MPPs, we performed single-cell chromatin immunocleavage sequencing (scCHICseq) analysis^59^ of H3K4me1 enhancer marks in CD62L^neg/high^ MPPs, treated with a non-toxic concentration (10mM) of the PPP inhibitor 6-AN (**Figure 5B-E; Figure S17B**). As H3K4me1 accumulates at enhancer to establish lineage commitment, we used this epigenetic modification to reveal potential commitment changes induced by the blockage of the PPP pathway. Following scCHICSeq library preparation, sequencing and QC, we analysed a total of 584 cells which had a coverage of 7,425 reads per cell (**Figure 5B-C, S18A-C**). First, several of the differentially represented enhancers in CD62L^high^ MPPs were associated with PPP perturbation *(Uap1, Pias1, Txn2, Tsc22d3, Myh9, Cdc42ep4)*. Many of the differentially represented enhancers were directly or indirectly associated with HSPC lineage specification and activation genes *(Brd4, Rac1, Elmo1, Ikzf1, Zeb2os, Fgl2)* (**Figure 5D**) in CD62L^high^ MPPs but not in CD62L^neg^ MPPs. In particular, *Ikzf1*, which encodes Ikaros, a master regulator of B-cell development was the most differentially enriched for H3K4me1 in CD62L^high^ MPPs versus CD62L^neg^ MPPs (**Figure 5D-E, S18D**). These results showed that CD62L^high^ MPPs responded to PPP inhibition by increasing H3K4me1 enhancer marks at the B-cell gene Ikaros, suggesting an epigenomic rewiring towards the B-lymphoid lineage.

In summary, PPP inhibition led to an increase of H3K4me1 marks at the enhancer of the master B-cell regulator Ikaros and other genes associated with HSPC lineage specification and activation specifically in CD62L^high^ cells. This suggested that the pentose phosphate pathway shapes the enhancer landscape linked to the commitment of myeloid versus lymphoid lineage.

### The Pentose Phosphate Pathway Regulates Immune-Cell Lineage Specification *In Vivo*

Next, to test whether the PPP-mediated epigenetic changes are functionally important *in vivo*, we use a murine model where Glucose-6-Phosphate-Dehydrogenase (G6PD), the rate limiting enzyme of the PPP, is overexpressed^60^ (**Figure 6A**). In this G6PD overexpression system a large genomic fragment (20.1Kb) of the entire human G6PD gene, including upstream and downstream regulatory sequences, was inserted into the genome of a transgenic mouse line (G6PD-Tg), leading to a 2-fold increase in enzyme activity^60^. By flow cytometry we confirmed that the transgene led to an increase in G6PD expression in MPPs as compared to wild-type littermate controls (**Figure 6B**).

**Figure 6.**
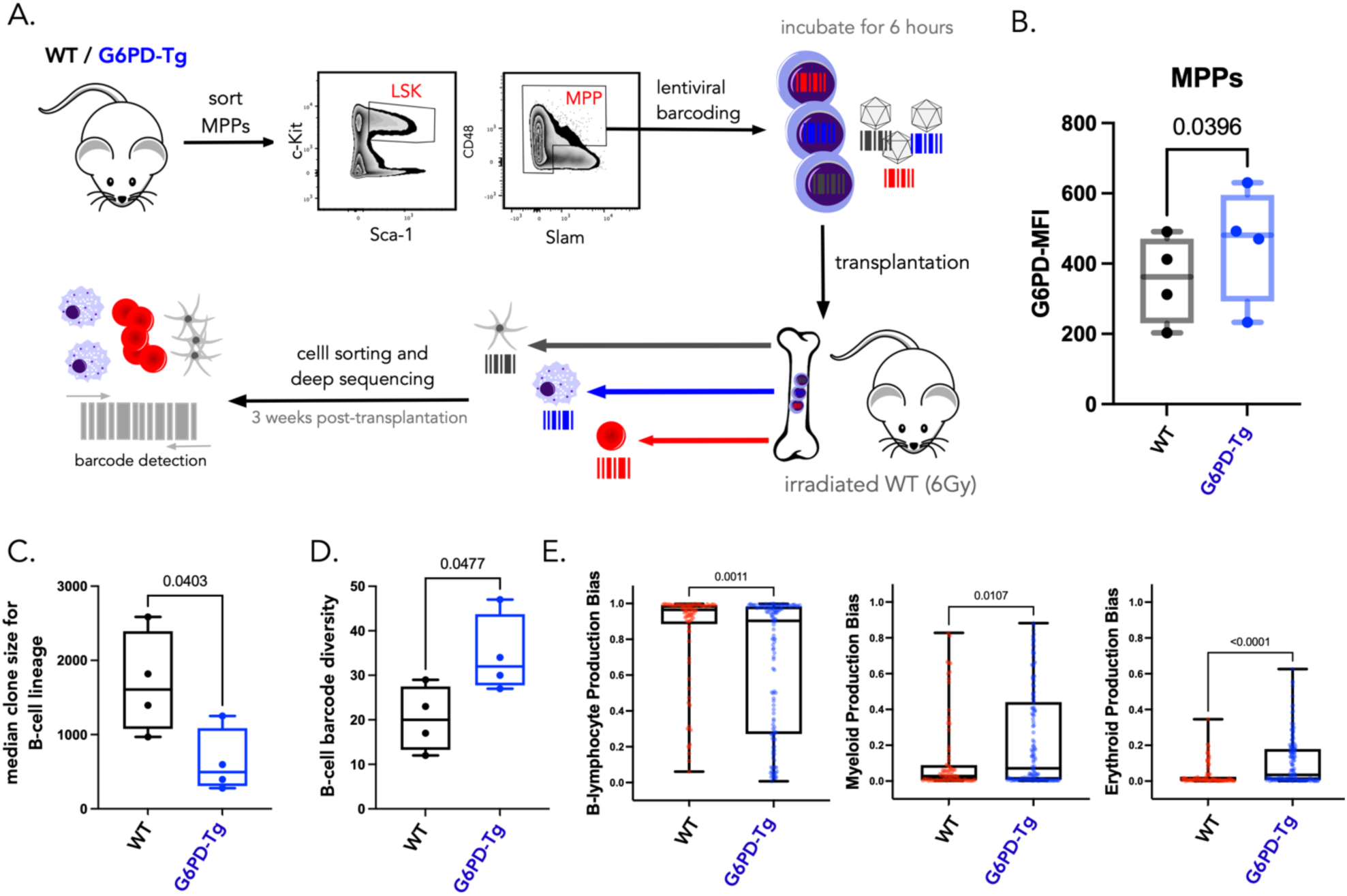
Upregulation of Glucose-6-Phosphate-Dehydrogenase in MPPs Inhibits B-Lymphopoiesis Post-Transplantation. (A) Overview of the lentiviral barcoding experiment. MPPs were purified from the bone marrow of WT or G6PD-Tg mice by FACS and were infected with the LG2.2 lentiviral barcoding library to achieve a multiplicity of infection of 10%. 6 hours later transduced 15,000-18,000 MPPs (10% of which have a barcode) were injected I.V. into 6Gy irradiated WT recipients. 3 weeks post-transplantation bone marrow was harvested and cells were sorted and their bulk DNA was processed for barcode detection through nested PCR and sequencing. (n = 4 mice per condition). At the 3 week timepoint we observe 10% chimerism of GFP+ barcoded cells (Figure S13b). (C) Median clone sizes for the top B-cell producing barcodes (the top *n* barcodes defined as contributing to 95% of all read counts for the B-cell lineage) from WT (black) and G6PD-Tg (blue) transplanted MPPs. Each point represents a single mouse. Pairwise comparisons are made using a Student’s T-test. (D) Barcode diversity as defined by the number of unique barcode for B-cell producing barcodes from WT (black) and G6PD-Tg (blue) MPPs, each point represents a single mouse. Pairwise comparisons are made using a Student’s T-test (E) The production bias of each B-cell producing barcode is shown per lineage and per experimental condition. The bias is calculated by scaling proportional read counts to cell numbers based on number of cells sorted per condition, then comparing the relative frequencies of each barcode across all lineages. Each point represents a single barcode (81 WT barcodes (red), 138 G6PDtg barcodes (blue)) with data pooled from 4 mice per condition. Pairwise comparisons were made using a Mann-Whitney test. Boxplots represent the median and interquartile range with whiskers extending to the minimum and maximum values.

To assess the functional consequences of G6PD overexpression on MPP fate, we transplanted G6PD-Tg and WT MPPs and quantified their cellular outputs *in vivo* using a single cell lineage lentiviral barcoding approach that allowed us to follow the fate of functionally heterogenous cell populations (MPPs) (**Figure 6A**). In this experiment, MPPs were purified from the bone marrow of G6PD-Tg mice or WT littermate controls and infected with the LG2.2 lentiviral barcode library as previously described (**Figure S19A**)^61^. Cells were then transplanted into sub-lethally irradiated (6Gy) recipients and left to engraft, divide and differentiate. At day 21 after transplantation, the timepoint where myeloid production from transplanted MPPs peaks^44^ , barcoded (GFP^+^) erythroblasts (E; Ter119^+^ CD44^+^), myeloid cells (M; Ter119^-^ CD19^-^ CD11b^+^), and B-cells (B;

Ter119^-^ CD11b^-^ CD19^+^) were sorted from the bone marrow and their barcode identity assessed from their bulk DNA (**Figure 6A and S19A**). We observed no significant differences in the diversity, clone sizes, and bias of the erythroid- and myeloid-producing MPPs between the G6PD-Tg and control group (**Figure S20C-H**). However when we analysed B-cell producing MPPs, we observed a 2.7-fold reduction in the number of B-cells derived from each barcoded MPP in the G6PD-Tg group compared to WT. WT B-cell barcodes displayed a median clone size of 1692 ± 689 cells, but only 630 ± 434 cells for G6PD-Tg B-cell barcodes **(Figure 6C).** The reduction of B-cells per MPP due to G6PD overexpression was partially compensated by an increase in the number of MPPs producing B-cells in the G6PD-Tg group compared to WT (**Figure 6D**). This result was corroborated by the analysis of chimerism post-transplantation, with a significant decrease in GFP^+^ cells in G6PD-Tg-derived leukocytes relative to WT controls. We observed a trending decrease in B-cell chimerism (p = 0.057) but no change in myeloid chimerism (p = 0.34) (**Figure S21**). Overall, per individual MPP, G6PD over-expression led to a net-skewing of cell production towards the erythro-myeloid lineages at the expense of the B-lymphoid lineage (**Figure 6E**).

Importantly, our phenotype could not be explained by differences in relative engraftment rates of MPPs, or by the transgene affecting cell viability or cell division (**Figure S22**). Specifically, there were no significant differences in the number of engrafted MPPs between G6PD-Tg versus WT conditions (p = 0.57, **Figure S22A**) and the clone sizes in MPPs were similar for both conditions (p = 0.18, **Figure S22B**), confirming no differences in MPP engraftment, expansion or viability in the BM. Furthermore, we saw no differences in B-cell or progenitor frequencies in the bone marrow of WT or G6PD-Tg mice, confirming that transgene expression did not perturb progenitor and B-cell viability and proliferation *in vivo* (**Figure S22C-E**).

Finally, we performed epigenomic profiling of G6PD-Tg or WT HSPCs, split into CD62L^high^ and CD62L^low^ MPPs using scCHICseq (**Figure S23A-C**). Several of the differentially represented enhancers in G6PD-Tg CD62L^high^ MPPs were associated with PPP perturbation (*Prkcd, Mgat5, Serpine, Mul1, Dctd, Hatatip2*) (**Figure S23E**) compared to WT CD62L^high^ MPPs. Some enhancers were directly or indirectly with HSPC lineage specification or survival, self-renewal, proliferation gene (*Etv6, IL-10*) (**Figure S23E**). Other enhancers pointed indirectly toward a disturbed myeloid production or function; for e.g. enhancers of genes that when mutated are associated to hematological disease (*Extl3, Ptprj, Pou4f1*) or link to inflammatory response and myeloid function (*trim30c, Adamts17, PxK, zak, Card14, nampt, hivep3, dapk1, ddp9*) (**Figure S23E**). This further supports our model that perturbations of the PPP, leading to epigenetic changes that influence myeloid associated lineage commitment.

In summary, by combining targeted genetics and cellular barcoding approaches, we showed with single cell resolution that G6PD overexpression limits skews HSPC outputs away from B cells and towards the myeloid lineages, consistent with our epigenetic profiling results (**Figure 5**). This result shows that metabolism can be targeted to control the magnitude and specificity of innate immune cell dynamics *in vivo*.

## Discussion

Recent work has shown that HSPCs play an active role in immune responses, constantly adjusting the type and amount of cells that they produce in response to changing demands. Importantly, changes in myelopoiesis due to inflammation play a pivotal role in many chronic diseases contexts including cancer, autoimmunity and metabolic disorders, perpetuating a cyclical pattern of inflammation. Previous work has highlighted the critical role of transcription and growth factors in controlling this feed-forward inflammatory loop, but the role of metabolism has been understudied. By combining *in situ* RNA barcoding, metabolic assays, and genetics approaches we identified the metabolic processes that regulate the magnitude and specificity of myelopoiesis. Specifically, we identified a developmental program of enzymes and transporters that facilitates myeloid lineage specification, and by perturbing enzyme expression it is possible to alter the epigenetics and immune cell production capacity of HSPCs. More broadly, our data illustrates that lineage commitment is an active process that requires energy and substrate inputs to meet associated epigenetic, biosynthetic and bioenergetic demands (**Figure 7**).

**Figure 7:**
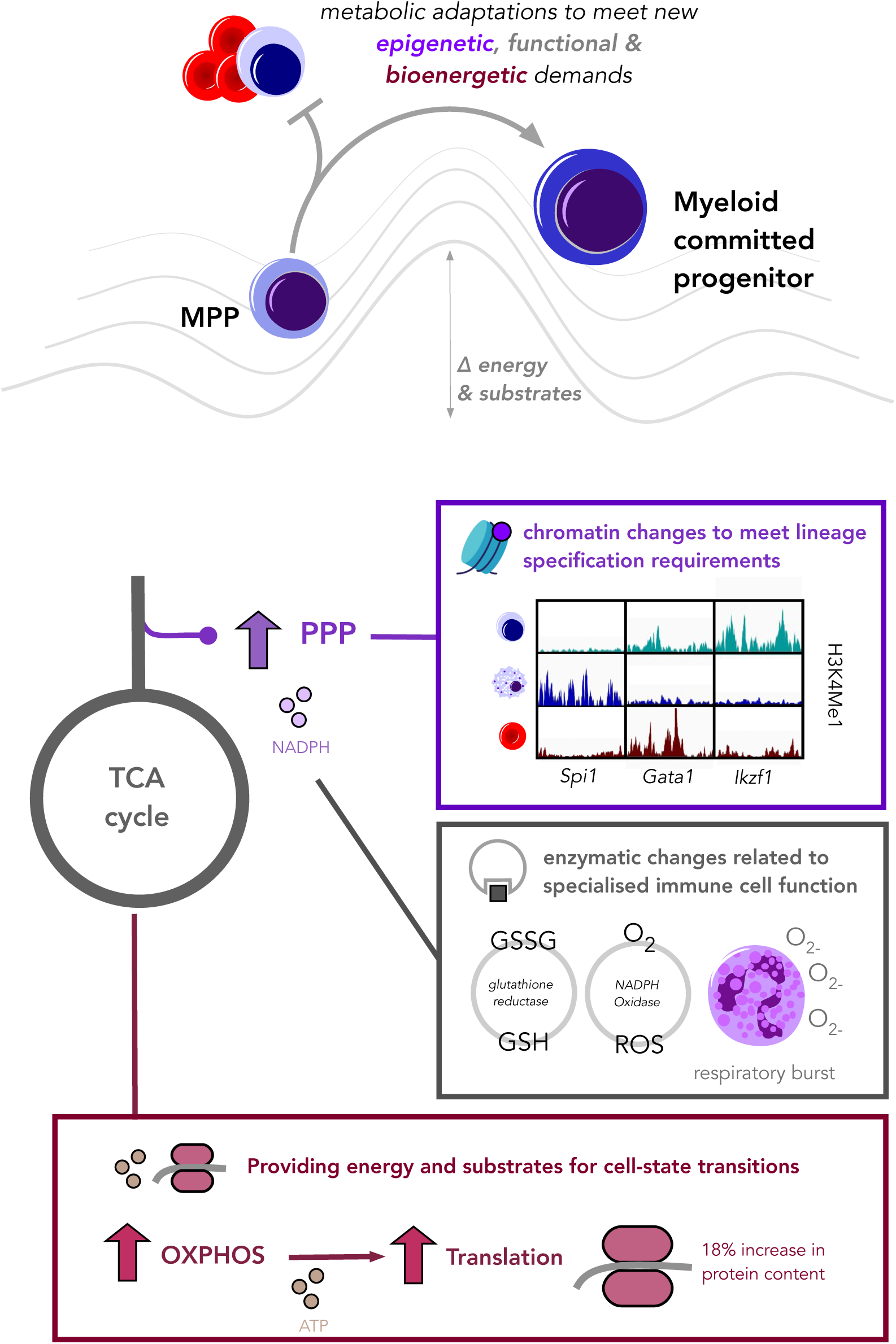
Pentose phosphate pathway regulates myeloid commitment by controlling the availability of key biosynthetic and epigenetic substrates. We posit that lineage commitment is an active energy consuming process In this setting metabolism fuels fate decisions by providing the energy, enzymes and substrates required to (i) alter gene expression patterns (ii) fulfil enzyme-dependent cell functions that are essential for myeloid function such as respiratory burst, and (iii) change cell size and molecular composition when transitioning from a stem cell state to an immune cell state. Supervised multi-omics analysis of transcriptomics, metabolomics and epigenomics datasets highlights a conserved link between the PPP, OXPHOS, redox, SAM/SAH pathways which collectively shape the availability of key substrates required for myeloid lineage commitment.

To link functional properties with metabolic properties of rare cell types *in vivo*, we devised a fate-resolved multi-omics strategy focused on enzyme and transporter state-fate mapping and follow-up metabolomics validation. This multi-modal approach enabled us to identify metabolic differences within the rare but functionally important MPP population, which collectively represent just 0.03% of total bone marrow cellularity^2^. Importantly, such differences were not resolved by other metabolomics studies^20^, highlighting the sensitivity of our approach. Given the diverse array of lineage tracing and metabolomics technologies that are emerging, our fate-resolved multi-omics approach can be readily adapted to other stem cell and developmental systems, including human tissues using human-compatible retrospective lineage tracing methods.

In this work, we identified an expression program of enzymes and transporters that confers differences in myeloid cells production within a subset of MPPs. Leveraging the ability of the DRAG barcoding system to detect barcodes at both RNA and DNA level, we were able to measure barcode abundances in both HSPCs and themature myeloid and nucleated erythroid progenitor compartments. This experimental design enabled us to trace lineage commitment over longer developmental trajectories, compared to studies that measure barcodes only in progenitors^32,42^. Using only genes encoding metabolic enzymes and transporters, our signature had a higher correlation with myeloid bias than the existing MPP3 signature^39,62^, prompting us to develop a novel purification strategy using the surface marker CD62L. Through *in situ* and lentiviral barcoding experiments, we show that CD62L enriches for myeloid bias in MPPs. In humans, the CD62L-coding gene *SELL* has been associated with abnormal myeloid cell counts^63,64^, suggesting that CD62L may have implications in the regulation of human hematopoiesis. Metabolically, CD62L^high^ MPPs have higher rates of ATP turnover and protein synthesis compared to CD62L^neg^ MPPs, with a higher energy dependence on OXPHOS. It is likely that these metabolic adaptations enable MPPs transition to the GMP cell state.

Our results also highlight a key role for the PPP in shaping <u>myeloid rather than lymphoid lineage identity</u>. Network analyses of gene expression data suggest that the PPP regulates several multiple aspects of myeloid function, including NADPH oxidase activity, glutathione recycling, and fatty acid synthesis (**Figure 7**). Importantly, we show that the PPP plays an active role in lineage specification. Overexpression of G6PD reduces the rate of B-cell production, resulting in a net skewing towards the myeloid lineages. This result is consistent with reports that pharmacological inhibition of the PPP blocks erythropoiesis *in vitro*^16^ and that the pathway regulates the function of dendritic cells^65^ and macrophages^66,67^. However, patients with inherited mutations in G6PD display hemolytic anemia, but no reported myeloid related disorders.

Integrating our single cell multi-omics results from transcriptomics, metabolomics and epigenomics with current literature supports a mechanistic model in which the PPP modulates lineage commitment. In this model, the PPP modulated the myeloid enhancer landscape in HSPCs by controlling the availability of substrates and co-factors through OXPHOS, glutathione redox metabolism and one-carbon metabolism (**Figure 7**). OXPHOS was a key signature of early myeloid commitment in CD62^high^ MPPs, as evidence by increased expression of OXPHOS related gene (*Idh2, Idh3a, Cox7b, Ndufa4, Uqcr10),* increased OXPHOS activity in both SCENITH and SPICE-Met and increased of OXPHOS-associated metabolites (carnitine-bound long-chain fatty acids, aspartic acid, and n-acetylaspartic acid). The increased PPP activity, evidenced by over-expression of key PPP genes (*Tkt, Taldo1, Gpi1, Pgls*) in CD62L^high^ HSPCs, can regulate glucose flux into the OXPHOS pathway, controlling the production of key substrates for biosynthesis and epigenetic state. Our metabolic causal network showed a direct causal association between *G6pdx* (PPP) and *Cox6B2* (OXPHOS). This is also supported by the increased ATP required for protein synthesis, as evidenced by our SCENITH and SPICE-Met experiments. It also in line with the literature showing that the PPP activity can affect epigenetic state via regulation of glucose catabolism^58^. PPP flux also heavily influences the activity of NADPH-dependent enzymes affecting epigenetics and redox state, for example isocitrate dehydrogenase^27^ and glutathione reductase^60^ which were identified in our transcriptomics analyses. Our network reconstruction model *G6pdx* showed a direct causal association with glutathione reductase *Gsr*. Moreover, the PPP genes (*G6pdx*, *Taldo*) associated directly with the glycolysis gene *Pkm,* which regulates the flux of carbon intermediates which can be diverted into synthesizing the serine required to fuel the 1C-folate cycle. The link between the PPP and one-carbon metabolism is further supported by our metabolomics experiments, in which we observed an altered SAM/SAH ratio in CD62^high^ MPPs. This result is supported by the literature, in which the PPP has been shown in other studies to regulate the folate cycle^68^, which provides the methyl group for the SAM/SAH production through the methionine cycle. Collectively, our multi-omics datasets support a mechanistic model in which the PPP regulates the availability of biosynthetic and epigenetic substrates and co-factors via OXPHOS, glutathione redox metabolism and one-carbon metabolism.

Understanding the nutrients and metabolites that regulate myelopoiesis can inform the development of novel bone marrow organoid technologies to maintain and differentiate hematopoietic precursors *ex vivo*. Our data can also help inform the development of nutrient/metabolite biomarker panels for stem cell function. Such tools can inform dietary interventions to promote HSPC function, particularly in prospective recipients of bone marrow transplants, that are susceptible to malnutrition and nutrient deficiencies^70,71^. Lastly, our results suggest that therapeutic interventions to alter the metabolism of HSPCs may not target all cells uniformly, given their underlying metabolic heterogeneity. Our approach may therefore be useful in studying the regulatory role of metabolism in other heterogeneous cell populations, such as cancer stem cells.

Overall, our data shows that the pentose phosphate pathway shapes the HSPC commitment to myelopoiesis. Our data also provide proof of principle that HSPC metabolism can be targeted to modulate the magnitude and specificity of immune cell regeneration *in vivo*.

### Limitations of the study

This study has several limitations inherent to the experimental systems and biological materials used. First, the work focused on rare hematopoietic stem and progenitor cells (HSPCs), which restricted the range of metabolic assay that could be performed. Second, while the sorting process may transiently alter cellular metabolism, we had no choice than performing metabolomic profiling on sorted cells. Third, for functional validation, we relied on transplantation assays to assess the specific effects on multipotent progenitors (MPPs). While transplantation assay is a gold-standard approach, it may not fully recapitulate steady-state biology^22^. Finally, for *in vivo* functional validation, we were constrained by the number of mice available, but we designed the study to ensure sufficient statistical power and robustness of all analyses, enabling us to draw reliable conclusions despite these constraints.

## Materials and Methods

A detailed description of all materials and method is provided in the supplementary information

## Acknowledgements

We thank the Institute Curie flow cytometry, next-generation sequencing, animal, and UMR168 BMBC facility. We thank thank Single-Cell Core of the Oncode Institute, Utrecht, the Netherlands for performing the single-cell CHICseq sample preparation. Finally we thank all the members of the Perié team, as well as Julio Sampaio Lopes, Mark Coles, and Antoine Marcais for helpful discussion.

## Author Contributions

**J.C.** conceptualization, performed experiments, data curation, data analysis, methodology, writing, funding acquisition. **A.D** Performed experiments, data curation, data analysis. **A.M.L.** data analysis, methodology, writing - review and editing. **N.K.** performed experiments. **H.D** performed experiments. **I.R-R.** performed experiments. **V.C.** data analysis, review and editing. **A.M** Performed experiments, data curation, data analysis. **C.C.** methodological development, performed experiments. **G.J**. performed data analysis. **S.T.B.** methodological development. **E.T.** methodological development. **E.R.** performed experiments. **F.T.** performed experiments. **Y.B.** performed experiments. **S.M.T.** performed experiments. **S.R.** performed experiments. **F.M.** performed experiments. **L.R.** performed experiments. **BDL.** provided expertise. **A.A.** provided expertise and reagents, review and editing **C.L.** provided expertise and reagents, performed experiments, review and editing. **P.B.** provided expertise and reagents, review and editing. **P.J.F.M.** provided expertise and reagents, review and editing. **C.V.** provided expertise and reagents, writing. **H.I.** Formal analysis, review and editing. **N.C.W.** provided expertise and reagents, review and editing. **R.J.A.** provided expertise and reagents, writing, review and editing. **L.P.** conceptualization, performed experiments, data analysis, funding acquisition, methodology, supervision, writing.

## Ethics

All the experimental procedures were approved by the local ethics committee CEEA-IC (Comité d’Ethique en expérimentation animale de l’Institut Curie) under approval numbers DAP 2016 006, DAP 2021-010 and DAP 2021-013

## Data Availability

All datasets generated or reanalysed during this study are available at: https://github.com/TeamPerie/Cosgrove-et-al-2022

## Code Availability

All source code generated during this study is available at: https://github.com/TeamPerie/Cosgrove-et-al-2022

