## supplementary information for "The Pentose Phosphate Pathway Regulates Myelo-Lymphoid Lineage Specification"

### Materials and Methods

**Mice:** All the experimental procedures were approved by the local ethics committee CEEA-IC (Comité d'Ethique en expérimentation animale de l'Institut Curie) under approval numbers DAP 2016 006, DAP 2021-010 and DAP 2021-013 and by the Institut Pasteur Safety Committee in accordance with French and European guidelines (CETEA 190148). Within each experiment mice were sex and aged matched and littermates were used to generate control cohorts. SCENITH and LPS challenge experiments were performed in C57BL6/J mice. Transplantation studies were performed in either CD45.1 C57BL6N / CD45.2 C57B6N mice or in HG6PD(Tg+) or HG6PD(Tg-) littermate controls. G6PD-Tg mice and littermate controls were generated the Spanish National Cancer Research Center (CNIO) at the Transgenic Mice core facility and were provided by Pablo Jose Fernández-Marcos. PercevalHR Vav-iCre mice were generated by the Centre d'Ingénierie Génétique Murine at Institut Pasteur and performed in accordance with the European Community guidelines (2010/63/UE) and were provided by P. Bousso. For DRAG barcoding experiments DRAG1 mice were crossed with B6.Cg-Tg(CAG-cre/Esr1*)5Amc/J (CAGGCre-ER$^{TM}$) to obtain heterozygous mice.

**Cell isolation:** BM cells were obtained from wild-type C57BL/6 of 8-16 weeks of age by bone flushing of femur tibia and iliac crest. Bone marrow cells were MACS enriched for cKit+ cells using CD117 MicroBeads Ultrapure (Miltenyi Biotec cat #130-091-224) according to manufacturer's protocol. Cells were kept in RPMI + 10% fetal calf serum + 1% pen-strep or in PBS + 10% fetal-calf serum + 1% pen-strep.

**Flow Cytometry:** Cell suspensions were incubated with antibody master mixes for 30 minutes on ice protected from ambient light. Cells were then washed in RPMI + 10% FCS + 1% pen-step for 2 x 5 minutes for 1700rpm at 4C. For TMRE staining of mitochondrial membrane potential cell suspensions were processed for flow cytometry surface staining and were then incubated in 200ul of 100nM TMRE solution. A full list of the antibodies used in this study is provided in Table S1. Samples were processed on a Ze5 (bio-rad), or Cytoflex LX (Beckman Coulter) plate-reader cytometers. Data analysis was performed using FlowJo v10.2 software (TreeStar), R v4.2.0, and Prism v9.

**Fluorescence activated cell sorting:** FACS was performed at the flow cytometry facility of Institute Curie on a FACSAria$^{TM}$ (BD Biosciences) or sh800 (Sony). Cells were sorted using a 70 mm nozzle at precision 0/16/0 and high efficiency in epperdendorf tubes.

**SCENITH:** cKit enriched murine bone marrow cells were seeded in 96 well plates for studying blood cells metabolism at 2 × $10^6$ cells/mL. Wells were treated during 30-60 minutes with Control, Oligomycin (Oligo, final concentration 1 µM), or the translation initiation inhibitor Harringtonine. 2-Deoxy-D-Glucose (DG, final concentration 100mM), was not used in our final analysis as treatment with this molecule for long time-periods (> 30 minutes) along with fixing and permeabilization of the cells led to shedding of CD62L from the membrane of MPPs. This effect was not observed in SPICE-Met processed cells, which are exposed to the same concentration of 2-DG, but are measured immediately after staining in live cells, without fixation and permeabilization steps. Puromycin (final concentration 10 µg/mL) is added at the same time as the metabolic inhibitor treatment. After puromycin treatment, cells were washed in cold PBS and stained with a combination of Fc receptors blockade and fluorescent cell viability marker (fixable live/dead nearIR, ThermoFisher Scientific, cat # L10119), then primary conjugated antibodies against surface markers (**Table S1**) during 25 min at 4°C in PBS 1X 5% FCS, 2mM EDTA (FACS wash buffer). After washing, cells were fixed and permeabilized using FOXP3 fixation and permeabilization buffer (Thermofisher eBioscience) following manufacturer instructions. Intracellular staining of puro using fluorescently labeled anti-Puro monoclonal antibody with Alexa Fluor 488 was performed by incubating cells during 1 h at 4°C diluted in permeabilization buffer. Mitochondrial dependencies were calculated as $((\text{Puro-MFI}_{DMSO+H2O} - \text{Puro-MFI}_{oligo}) / (\text{Puro-MFI}_{DMSO+H2O} - \text{Puro-MFI}_{H}))$ x100.

**SPICE-Met:** cKit enriched cells were purified from the femur, tibia and iliac bones of vav-iCre Perceval$^{fl/fl}$ mice. Following surface staining with antibodies (**Table S1**), cells expressing PercevalHR were treated with either 1mM oligomycin, 100 mM 2-deoxy-D-glucose (2-DG), the combination of the two or DMSO as control and kept at 37°C for 30 minutes and then analyzed by flow cytometry. Samples were then recorded for one minute on a Cytoflex LX (Beckman Coulter). ATP:ADP ratios in individual cells were calculated from two fluorescence signals. In brief, excitation with a violet laser (405nm) and signal detection with a 525 band-pass filter was used to measure ADP contribution while ATP levels were estimated by excitation with a blue laser (488nm) and signal detection with a 525 band-pass filter. Data analysis was performed using FlowJo. OXPHOS and glucose contribution were calculated as $((R_{DMSO+H2O} - R_{oligo}) / (R_{DMSO+H2O} - R_{oligo+2\text{-}DG}))$ x100 and $((R_{DMSO+H2O} - R_{2\text{-}DG}) / (R_{DMSO+H2O} - R_{oligo+2\text{-}DG}))$ x100, respectively where R represents the ratio of fluorescent intensity measurements for ATP and ADP .

### Analysis of the Pentose Phosphate Pathway:

**G6PD protein expression by intracellular flow cytometry:** cKit enriched bone marrow cells were labelled with primary conjugated antibodies against surface markers before being fixed and permeabilized using FOXP3 fixation and permeabilization buffer (Thermofisher eBioscience) following manufacturer instructions. Intracellular staining of G6PD using fluorescently labelled antibody with Alexa Fluor 488 was performed by incubating cells during 1 h at 4°C diluted in permeabilization buffer.

**Sensitivity to PPP inhibitor 6-AN:** cKit enriched murine bone marrow cells were seeded in 96 well plates in StemSpan + TPO (50ng/ml) + SCF (50ng/ml) as well as 6-AN (10uM, resuspended in DMSO) or a DMSO vehicle control at a density of $1 \times 10^6$ cells/well. Cells were then incubated at 37C for 16 hours before labelling antibodies targeting key surface markers (Lin, Sca1, cKit, CD150, CD48, CD62L) and cell viability markers.

### Targeted Mass Spectrometry:

**Targeted metabolomics of polar metabolites:** For low-input targeted metabolomics on polar metabolites 5,000 HSCs (LSK CD150+CD48-) and MPPs (LSK minus HSC) were sorted using flow cytometry into 25ul ice-cold Acetonitrile. Targeted metabolite quantification by LC-MS was carried out using[1]. An Agilent 1290 Infinity II UHPLC was coupled to an Agilent 6495 QQQ-MS operating in MRM mode. MRM settings were optimized separately for all compounds using pure standards. For every target metabolite, one quantifier and 1 to 2 qualifiers were recorded. Chromatographic separation was performed using a HILICON iHILIC(P) classic column (100 mm × 2 mm, 5 µm particles) or a Waters Atlantis Premier BEH ZHILIC column (100 mm × 2.1 mm, 1.7 µm particles). Buffer A was 20 mM ammonium carbonate and 5 µM medronic acid in Milli-Q H2O and Buffer B was 90:10 acetonitrile/buffer A. The gradient profile was: 0 min, 95% B, 120 µL/min; 18 min, 55% B, 120 µL/min; 19 min, 20% B, 120 µL/min; 21.5 min, 20% B, 120 µL/min; 22 min, 95% B, 120 µL/min; 23.5 min, 95% B, 120 µL/min, 25.5 min, 95% B, 300 µL/min; stop time 30 min. Irrespective of the BEH ZHILIC column used, the flow rate was increased to 150 µL/min from 0 min to 23.5 min. The injection volume was 20 µL, the column temperature was 40 °C, and the autosampler temperature was 5 °C. MS source parameters were as follows: gas temp: 240 °C, gas flow: 15 L/min, nebulizer: 50 psi, sheath gas temp: 400 °C, sheath gas flow: 11 L/min, capillary voltage: 2000 V, and nozzle voltage: 300 V. iFunnel parameters were as follows: high-pressure RF positive: 110 V, high-pressure RF negative: 90 V, low-pressure RF positive: 80 V, and low-pressure RF negative: 60 V. At least three negative controls were measured for each experiment to evaluate metabolite background levels. Metabolites were only included if they were detected above background (sheath fluid and cell debris sorted from the same sample tube) levels and retention time was identical to the 13C yeast standard qualifier peak. To evaluate metabolic differences, the peak height was determined and subtracted for the associated peak height for negative control samples (sorted debris).

### scCHICseq profiling of H3K4Me1 marks

**Sample preparation:** cKit enriched murine bone marrow cells were seeded in 48 well plates in StemSpan + TPO (50ng/ml) + SCF (50ng/ml) as well as 6-AN (10uM) or a DMSO vehicle control. Cells were then incubated at 37C for 16 hours before processing for FACS purification. Here 50,000 cells were sorted for each cell type ($CD62L^{neg/high}$ MPPs) and experimental condition (6-AN treated and DMSO control) giving 4 samples in total. Cell suspensions were washed, filtered and passed through a Protein LoBind tube prior to ethanol fixation and snap freezing at -80C. scCHICseq profiling of H3K4Me1 was then performed by the Single-Cell Core of the Oncode Institute, Utrecht, the Netherlands. Briefly each celltype-treatment condition combination corresponds to a distinct plate and were processed for nuclei extraction, antibody labelling, pA-Mn targeting and sequencing library preparation as described in the original protocol defined by Zeller et al[2]. **Data analysis:** The data preprocessing has been performed using the SingleCellMultiOmics package and the sequences were mapped to the mus musculus genome (GRCm39 ensembl, version 97) using Burrows-Wheeler Aligner (BWA). Duplicated fragments were removed based on genomic coordinates. The data were then aggregated into 50 kb bins and imported into the ChromSCape[2] analysis framework. Cells with low read coverage ($<10^3$ reads per cell) were excluded, as were cells in the upper 95th percentile of read coverage, which were considered likely doublets. To define a peak set for dimensionality reduction and differential analysis, we selected the union of the top 5,000 most covered 5 kb bins from each sample. This resulted in a final set of 9,148 peaks. These quality-control steps were guided by recommendations from a recent benchmarking study proposing best practices for the analysis of H3K4me1-based epigenetic datasets[3]. After quality control, a total of 584 cells were retained for downstream analyses, with a median coverage of 7,425 reads per cell. For data visualization, TF-IDF normalization was applied, followed by latent semantic indexing (LSI) and UMAP. The first LSI component was excluded because it was correlated with sequencing depth, and the subsequent 20 components were used for downstream embedding. For differential analysis, CPM normalization was applied, and differential signal was assessed using the negative binomial method implemented in ChromSCape[4]. For genome track visualization, bigWig files from each experiment were plotted using the plotgardener package[5].

### Lentiviral Barcoding

**Lentiviral Barcoding Transduction**: The barcode library LG2.2 was used as in Eisele *et al* (2022)[6]. In brief, lentiviruses were produced by transfecting the barcode plasmids and p8.9-QV and pVSVG into HEK293T cells in DMEM-Glutamax supplemented with 10% FCS (Gibco), 1% MEM NEAA, and 1% sodium pyruvate using Polyethyleneimine. Supernatant was 0,45 um filtered, concentrated by 1h30 ultracentrifugation at 31 000g and frozen at -80°C. For isolation and labelling of cells with the LG2.2 lentiviral barcoding library bone marrow cells were isolated from femur, tibia and iliac bones by flushing using a 21G needle (Terumo), and C-Kit$^+$ cells enriched using anti-CD117 magnetic beads (Miltenyi) on the MACS column system (Miltenyi). Cells were stained with surface antibodies (**Table S1**) and sorted at the flow cytometry facility of Institute Curie on a FACSAria$^{TM}$ (BD Biosciences) or sh800 (Sony). Cells were sorted using a 70 mm nozzle at precision 0/16/0 and high efficiency in epperdendorf tubes. MPPs were transduced with the lentiviral barcode library in StemSpanMedium SFEM (STEMCELL Technologies) supplemented with 50 ng/ml mSCF (STEMCELL Technologies) through 1,5 h of centrifugation at 300 g followed by 4,5 h incubation at 37°C in order to obtain 10% barcoded cells. After the incubation 15-18,000 cells were injected in the tail vein of 6 Gy sub-lethally irradiated recipient mice.

**Lentiviral Barcode amplification and sequencing:** Bones (femurs, tibias and ilia) were isolated for barcode analysis from recipient mice. Bone marrow cells were extracted by flushing of the bones using a 21G needle (Terumo) and enriched using anti-CD117 magnetic beads (Miltenyi) on the MACS column system (Miltenyi). The ckit$^-$ fraction was further separated into a Ter119$^+$ and Ter119$^-$ fractions using anti-Ter119 coated magnetic beads (Miltenyi). Cells were stained with fluorescently conjugated antibodies (**Table S1**) and after sorting mature cells were lysed in 40 µl Viagen Direct PCR Lysis Reagent (cell) (Euromedex) supplemented with 0,5 mg/ml Proteinase K Solution RNA grade (Invitrogen) in a thermic cycler: (55°C for 120 min, 85°C for 30 min, 95°C for 5 min, indefinite at 4°C). Samples were then split into two replicates, and a three-step nested PCR was performed to, in a first step amplify barcodes (primers top-LIB (5'TGCTGCCGTCAACTAGAACA-3') and bot-LIB (5'GATCTCGAATCAGGCGCTTA-3')), in a second step add unique 4 bp plate indices (forward 5'ACACTCTTTCCCTACACGACGCTCTTCCGATCTNNNNCTAGAACACTCGAGATCAG3' and reverse 5'GTGACTGGAGTTCAGACGTGTGCTCTTCCGATCGATCTCGAATCAGGCGCTTA3'), and in a third step add P5 and P7 flow cell attachment sequences and one of 96 sample indices of 7 bp P5 5'AATGATACGGCGACCACCGAGATCTACACTCTTTCCCTACACGACGCTCTTCCGATCT3' and P7 5'CAAGCAGAAGACGGCATACGAGANNNNNNNGTGACTGGAGTTCAGACGTGCTCTTCCGATC3') (PCR program: hot start 5 min 95°C, 15 s at 95°C; 30 s at 57.2°C; 30 s at 72°C, 5 min 72°C, 30 (PCR1-2) or 15 cycles (PCR 3)). Both index sequences (sample and plate) were designed based on[7] such that sequences differed by at least 2 bases, and homopolymers or more than 2 bp, hairpins and complementary regions with the rest of the primer sequence were absent. To avoid lack of diversity at the beginning of the reads during sequencing, at least 4 different plate indices were used for each sequencing run. Primers were ordered desalted, and high-performance liquid chromatography (HPLC) purified. During lysis and each PCR, a mock control was added. The DNA amplification by the three PCRs was monitored by the run on a large 2% Agarose gel. Samples were pooled in order to guarantee a sequencing depth of 50 reads/cell. Five µl of the products of PCR3 for each sample and replicate were pooled, purified using the Agencourt AMPure XP system (Beckman Coulter), analyzed on a Bioanalyzer, and diluted to a concentration of 5 nM. These pools were sequenced on a HiSeq system (Illumina) (SR-65bp) at the sequencing facility of Institute Curie (10% of Phix Illumina phage genome library was added to generate a more diverse set of clusters).

**Lentiviral Barcode Analysis:**

Sequencing results were analyzed using R-4.2.0, Microsoft Excel (v16.16, MAC edition), and GraphPad Prism version 9.0.

**Data demultiplexing:** Reads were first filtered for perfect match to the input index- and common-sequences using XCALIBR (https://github.com/NKI-GCF/xcalibr) and filtered against a barcode reference list.

**Data QC:** The consistency of technical replicates for each sample were then assessed using a Pearsons Correlation but no filtering was applied based on this metric. Barcodes that were not present in both technical replicates were filtered from the data with 3606/3969 WT barcodes and 3461/3942 G6PD-Tg barcodes retained at this step.

We then assessed the prevalence of repeat-use barcodes in our dataset. In this context barcodes that occur in more than one mouse per transduction batch are likely to be the result of more than 1 cell being labelled with the same barcode. To estimate repeat-use frequency in our datasets we first had to ascertain which repeat use barcodes were the result of repeat labelling versus those that arose due to sequencing errors. To do this, we ranked repeat use barcodes based on how many sequencing reads mapped to each. Barcodes below the upper 95$^{th}$ percentile of read abundance are considered as sequencing noise and set to zero in each respective sample. Barcodes which are in the upper 95$^{th}$ percentile of read abundance and only found in one mice after noise correction are retained for further analysis. Following this filtering step 819 unique WT barcodes and 805 unique G6PD-Tg barcodes were retained for analysis.

**Data normalisation:** After filtering, we normalised barcode counts within each sample. In a barcode count matrix where rows are barcodes and columns are samples, we normalize the data so that each column sums to 1. More precisely, Let $R_{bc}$ represent the number of reads for barcode *B* in cell type *C,* and let $P_{bc}$ represent the proportional read abundance per barcode per cell type:

$$P_{bc} = \frac{R_{bc}}{\sum_b R_{bc}}$$

When calculating clone sizes for each barcode it is relevant to know the number of cells produced and so we scale normalized sequencing reads by the number of cells in each sample, giving the cell-scaled barcode frequency $C_{bc}$. During data acquisition, we do not always measure the entire tissue sample. In this setting the number of cells sorted by FACS is corrected by accounting for the fraction of the total sample that was measured. As a concrete example if a bone marrow cell suspension was placed in 5ml of medium but only 4ml of this medium was acquired during cell sorting we scale the number of cells sorted by the inverse of the proportion of the sample that was measured. (1/0.8 = 1.25 in this example)

$$C_{bc} = P_{bc}\ x\ (number\ of\ cells\ sorted * \frac{1}{fraction\ of\ total\ sample\ measured})$$

**Calculation of barcode diversity and clone sizes:**

Barcode diversity was calculated as the total number of unique barcodes found in a given sample, and where relevant the number of unique barcodes found in 2 developmentally related samples, for example CD62L$^{high}$ MPPs and myeloid cells. Clone sizes were calculated from the cell-scaled barcode frequencies ($C_{bc}$) of a given barcode in a given sample.

**Calculation of lineage bias score:** To classify barcodes by their lineage bias, an additional normalization step per barcode is applied in each individual, thereby enabling categorization of each barcode into classes of biased output towards the analyzed cell types. Let lineage-bias$_{bc}$ represent the relative representation of barcode *B* in cell type *C,* and let $P_{bc}$ represent the proportional read abundance per barcode per cell type

$$lineage\ bias_{bc} = \frac{P_{bc}}{\sum_c P_{bc}}$$

**Calculation of production bias:** In instances where the number of cells produced per progenitor is relevant to the assessment of bias, we perform the same calculation as for lineage bias but use cell-scaled barcode frequencies $C_{bc}$ as opposed to the proportional read abundance per barcode per cell type.

$$Production\ bias_{bc} = \frac{C_{bc}}{\sum_c C_{bc}}$$

### MetaFate In Situ RNA Barcoding

#### Induction of Barcode Recombination

8-11 weeks old male mice received 7 mg Tamoxifen/ 40 gr bodyweight each day for 3 consecutive days by intraperitoneal injection. Tamoxifen (T5648-1G, Sigma) was dissolved 10% EtOH and 90% sunflower oil.

#### Cell isolation and sorting

At sacrifice, BM was harvested from femurs, tibias and ilia and enriched using anti-CD117 magnetic beads (Miltenyi). The c-kit$^+$ fraction was stained with antibodies against CD117 (c-kit APC, clone 2B8, Biolegend), Sca-1 (Pacific Blue, clone D7, eBioscience). The c-kit$^-$ fraction was stained with antibodies against CD11b (PercPCy5.5 or Pacific Blue, clone M1/70, eBioscience), CD19 (APC-Cy7, clone 1D3, BD Pharmingen). FACS was performed at the flow cytometry facility of Institute Curie on a FACSAria$^{TM}$ (BD Biosciences) or sh800 (Sony). Data analysis was performed using FlowJo$^{TM}$ v.10 (TreeStar). Cells were sorted using a 70 mm nozzle at precision 0/16/0 and high efficiency.

**10X Genomics 3' Library preparation for single-cell transcriptomics analysis:** DRAG barcoded LSK (ckit$^+$ sca1$^+$ gfp$^+$) cells were sorted from ckit-enriched bone marrow fraction. Samples were then processed using the 10X genomics Chromium Single Cell 3' v3 kit. Specifically, 1,000-16,000 cells were loaded for each experiment for a targeted recovery of 500-10,000 cells. cDNA amplification was performed with 11-13 PCR cycles depending on the targeted cell recovery, as per the manufacturers recommendations. Sequencing was performed on a NovaSeq (illumina) on paired end PE28-8-91 mode. Raw sequencing reads were processed using Cellranger.

**Targeted RNA-Barcode Recovery from 3' scRNA-seq libraries:** 10ng of our cDNA libraries underwent a cDNA pre-amplification using the 10X 3' amplification mix, partial read1 F primer (1uM), PreAmp + read2 reverse primer (1uM) and $H_20$ and were incubated at 98C for 3 minutes followed by 15 cycles of 15 seconds at 98C, 20 seconds at 63C and 60 seconds at 72C, samples were then incubated for 60 seconds at 72C. Samples then underwent SPRI size selection at a ratio of 0.6X before undergoing a sample index PCR. 10ul of our preamplification product was placed in a master mix of 10X 3' amplification mix along with P5- read 1 forward primer (1uM) and P7 reverse primer (1uM). Each sample then underwent PCR amplification with incubation at 98C for 45 seconds, followed by 8-16 cycles (according to **Table S2**) of 98C for 20 seconds, 63C for 30 seconds, and 72C for 20 seconds. samples then underwent a 60 second incubation at 72C. Samples then underwent an additional SPRI cleanup at 0.8X. Sequencing was performed on a MiSeq (Illumina) on paired-end 300bp mode.

**Targeted DNA-Barcode using PCR and deep sequencing on bulk mature cells.**

**Lysis:** Sorted cells were lysed 40 ul DirectPCR Lysis Reagent (Cell) from Viagen Biotech with 0.4 mg Prot K, and incubated for 1 h at 55°C, followed by 30 min at 85°C heat inactivation and 5 min at 94°C. Samples were stored at -20°C. **Preamp PCR:** Samples were resuspended in 200 µl PCR mix (20 µl 5x phusion HF buffer (NEB); 1 µl Phusion DNA Polymerase (NEB); 2 µl 10 mM dNTP's; 0.5 µl 100 uM preamp forw. oligo; 0.5 µl 100 uM preamp rev. oligo; 76 µl PCR grade water) and split in two replicates. PCR program: 2 min. at 98ºC; N* cycles of 10 sec at 98ºC, 20 sec at 60ºC, 25 sec at 72ºC; 5 min. at 72ºC; hold at 4ºC (* N is adjusted to number of barcoded cells in the sample, so at the moment of tagging every sample has the same number of molecules (See table in[8]). **Tagging PCR:** 2 µl preamp PCR product was mixed with 48 µl tagging PCR mix (10 µl 5x phusion HF buffer (NEB); 1 µl Phusion Hot Start II DNA Polymerase (2U/ µl) (NEB); 1 µl 10 mM dNTP's; 0.25 µl 1 µM M1 tag forward oligo; 35.75 µl PCR grade water). PCR program: 1 min. at 98ºC; 2 cycles of 10 sec at 98ºC, 2 min. at 57ºC, 1 min. at 72ºC; 20ºC forever). To digest remaining M1 tag forward oligo: 3 µl 20U/ µl Exonuclease I (NEB) was added and incubated for 1 h at 37ºC, denaturing ExoI was done for 5 min at 98ºC, and the sample cooled down to 20ºC. 0.5 µl 100 µM Illumina forward seq oligo and M1rev oligo (both had 2 Phosphorothioate bonds at the 3'end to avoid breakdown from residual ExoI activity) were added, followed by PCR program: 1 min. at 98ºC; 30 cycles of 10 sec at 98ºC, 20 sec at 67ºC, 25 sec at 72ºC; 5 min. at 72ºC; 4ºC forever. **Sample index PCR:** 2 µl tagging PCR product was mixed with 18 µl PCR mix (4 µl 5x phusion HF buffer (NEB); 0.4 µl Phusion DNA Polymerase (NEB); 0.4 µl 10 mM dNTP's; 0.1 µl 100 uM P5 forw. oligo; 4 µl 2.5 uM P7 index rev. oligo; 9.1 µl PCR grade water). PCR program: 30 sec at 98ºC; 15 cycles of 10 sec at 98ºC, 20 sec at 67ºC, 25 sec at 72ºC; 5 min. at 72ºC; 4ºC forever). **NGS sequencing:** 5 µl from each index PCR was taken and pooled before being cleaned using a SPRI selection (ratio 1:1) The pooled samples were deep-sequenced on a MiSeq System (Illumina) in SR150bp run mode

**RNA DRAG Barcode Preprocessing and Filtering:** Cell Ranger was used to process the reads from the targeted sequencing, aligning them to the mouse genome and adding 10x cell and unique molecular identifier (UMI) barcodes. Unmapped reads were then extracted from the original 10x and targeted bams using 'samtools view -f 4' and concatenated. An exact grep match to 52 base pairs at the 5 prime (V) (corresponding to the targeted amplification primer) end of the DRAG lineage barcode plus 10 base pairs at the 3 prime (J) end, allowing 10-40 variables bases in between, was used to extract reads potentially containing a lineage barcode. This was designed such that the start and end of the variable bases matches the barcodes extracted from the DNA. These reads were further filtered to keep only those with a 10x cell barcode and UMI assigned, and then to keep only reads in cells (as defined by Cell Ranger).

To assign one VDJ barcode to each 10x cell barcode, we iterated through each 10x cell barcode, extracting all relevant reads. Then, UMIs were filtered to keep only those with 3 or more reads and one dominant VDJ barcode (defined as >=0.45 of total reads of that DRAG barcode for a given cell). The dominant barcode for each UMI was extracted, and finally we assign one VDJ barcode to a 10x cell if we have good agreement across UMIs, defined as >=0.75 of UMIs for that cell mapping to one VDJ sequence. If there is only one UMI retained, we further ensure that the VDJ barcode for this UMI is the dominant barcode across all the reads for that cell and has >=0.45 of reads.

As with VDJ barcodes extracted from DNA, we perform further checks and filters to ensure the barcodes are of good quality and unique. First we check that the barcodes conform to the expected VDJ structure, using an algorithm[8] to compare barcodes to the original VDJ template and identify which nucleotides were deleted due to exonuclease activity and which ones were inserted due to Tdt activity. This enabled us to recognize barcodes containing residual error, and also to quantify barcode creation patterns (i.e. the number of deleted and inserted nucleotides at the junctions between V, D and J segments per barcode). We used this approach to filter barcodes that did not have the expected barcode recombination pattern (this only removes ~1% of barcodes). We then use the iGoR algorithm to compute a generation probability for each barcode[8].

**Assessing the Consistency of RNA and DNA barcodes:** Following QC and filtering (materials and methods), we recovered RNA barcode information for 668 hematopoietic stem and progenitor cells (14.9% recovery rate at the RNA level), corresponding to 158 unique lineage barcodes (**Table S4**). From the mature erythroid and myeloid bone marrow compartment, we recovered 381 unique barcodes at the DNA level with high consistency between technical replicates (**Figure S3**), with 97 barcodes overlapping between RNA and DNA detection. Comparison of lineage barcodes detected from either DNA or RNA showed similar barcode lengths, as well as similar insertion and deletion patterns, confirming that full barcode sequences could be accurately recovered from transcripts (**Figure S1B-D).**

**DNA Barcode Preprocessing and Filtering.**

Each recombined sequence includes nucleotide additions and deletions (referred to as the 'DRAG barcode') and constant parts that flank both sides of this barcode. Moreover, each barcode was associated with a random unique molecular identifier (UMI) of 12bp during the tagging PCR step.

**DNA DRAG barcode Preprocessing.** We use the pipeline described below to demultiplex fastq files and identify the reads that match a potential recombination of the DRAG construct. First the bcl2fastq (Illumina) program is used to demultiplex the fastq files based on the i7 index sequence. Only records that match the i7 index perfectly are considered for the next step. In the constant part of the V and the J, the reads tend to be error-prone and a consensus sequence with Ns is manually created. The Xcalibr program (https://github.com/NKI-GCF/xcalibr) is then used to extract counts for all combinations of the 16bp UMI and the recombined barcodes only for the reads that contain the constant sequence of the V at the expected coordinates. After this, the J constant part (tagcaagctcgagagtagacctactggaatcagaccgccaccatggtgagc) is aligned to the barcode part using the NBCI blast2 program. When a suitable match is found, the barcode is trimmed at the start coordinate of the match, resulting in the final matrix.

**DNA DRAG Barcode Processing.** We used the steps described below to identify barcode sequences and remove PCR and deep-sequencing errors. First, we removed any barcode and associated UMI containing one or multiple 'N' values (within either the barcode, constant flanking parts or UMI). Second, barcodes that did not have an exact match to the expected constant parts of the V and J that precede or follow the barcode were removed. Third, when multiple sequences were found associated with a single UMI, only the most frequently occurring barcode associated with that UMI was kept. As a fourth step, we summed up the read counts for all UMI associated with the set of remaining barcodes. To verify that the barcodes obtained match the expected structure of a VDJ recombination product, we developed an algorithm to compare barcodes to the original VDJ template and identify which nucleotides were deleted due to exonuclease activity and which ones were inserted due to Tdt activity. This enabled us to recognize barcodes containing residual error, and also to quantify barcode creation patterns (i.e. the number of deleted and inserted nucleotides at the junctions between V, D and J segments per barcode)[8].

**Single-Cell RNA-seq Analysis:** Raw sequencing reads were processed using Cellranger. To obtain a reads/cell/gene count table, reads were mapped to the mouse GRCm38.84 reference genome. During filtering, Gm, Rik, and Rp genes were discarded as noninformative genes. Cells with less than 500 genes per cell and with a high percentage (> 10%) of mitochondrial genes were removed from downstream analyses. Following our filtering procedures, the average UMI count per cell was 11829. The median number of genes detected per cell was 3812, 3.4% mapped to mitochondrial genes. Cell cycle annotation using the cyclone method from the scran R package[9] showed that 3366 cells were in G1 phase, 869 cells were in G2M phase, and 250 cells were in S phase. Data normalization and integration were performed using the default Seurat v4 approach FindIntegrationAnchors() followed by IntegrateData(), and differentially expressed genes were determined using a logistic regression in Seurat on the non-integrated data using the FindConservedMarkers() function. Pathway based analyses were performed using the enrichR package[10]. Unsupervised clustering was performed on the significant variable genes using the 10 first PCA followed by the nonlinear dimensionality reduction technique UMAP[11]. Annotation of the data was obtained by mapping published signatures using the AddModuleScore() method of Seurat. The MolO LT-HSC signature was taken from Wilson et al., 2015[12], and the MPP2/3/4 signatures were taken from Sommerkamp et al., 2021[13]. An Excel file listing the genes in these signatures is available in Supplementary Table S7. For permutation testing we randomly sampled 200 sets of genes to create gene-sets of an equivalent size to our MetaFate, Fate, transcription factor and MPP3 signatures. Expression scores for each cell were computed using the AddModuleScore() function from Seurat and the correlation between signature scores and myeloid bias was computed using a Spearmans Correlation.

**Categorisation of lineage biased barcodes:**

Labelled cells were classified as differentiation inactive (98 cells ; 61 unique barcodes) if we could not detect their respective barcode in any mature cell compartments, or as erythroid-biased (143 cells ; 35 barcodes), myeloid-biased (143 cells ; 31 barcodes), or unbiased (284 cells ; 31 unique barcodes). Lineage biases were defined based on the relative abundance of the barcode across the respective compartments. More precisely, an additional normalization step per barcode is applied in each individual, thereby enabling categorization of each barcode into classes of biased output towards the analyzed cell types. Let lineage-$bias_{bc}$ represent the relative representation of barcode *B* in cell type *C,* and let $P_{bc}$ represent the proportional read abundance per barcode per cell type

$$lineage\ bias_{bc} = \frac{P_{bc}}{\sum_c P_{bc}}$$

Specifically, a barcode was classified as myeloid biased if it was found above the 75th percentile of all myeloid-bias$_{bc}$ scores. A barcode was classified as erythroid biased if it was found in below the 25th percentile of myeloid-bias$_{bc}$ scores. A barcode was classified as unbiased if it was found between the 25th and 75th percentile of myeloid-bias$_{bc}$ scores.

These specific thresholds were set using a sensitivity analysis approach that recorded the number of cells in each category, as well as the number of differentially expressed genes and log2 fold change in gene expression between barcoded subsets. In this setting we observe that stricter thresholds in lineage bias led to greater effect sizes in gene expression changes, but at the expense of having less cells in each category (**Figure S3**). Thresholds of 25% and 75% were chosen to keep a strict criteria for lineage bias, whilst also having > 100 barcoded cells within each lineage bias category (**Figure S3**)

**Defining Metabolically-Associated Genes:** Genes are classified as metabolically associated if they are found within a metabolic pathway defined in the KEGG database, or whether they map to reactome/GO pathways associated with the following terms: *'transport', 'import','Slc', 'Abc','Atp','abc', 'metabolic process','biosynthetic process','catabolic process'*. In total 3,095 genes met this definition and the list of genes can be found in Supplementary Table S7.

**PAGA Trajectory Analysis:** To perform trajectory analysis, a reference map was generated from a published single-cell sequencing dataset of 44,802 C-Kit+ cells[14]. Preprocessing was performed using a scanpy pipeline as performed in (Wolf et al., 2019) and trajectory inference was performed using the PAGA method implemented in scanpy[15]. Data was then imported into Seurat and visualised using the dimensionality reduction technique UMAP. Data imputation was performed using the Rmagic library[16] before calculating the expression patterns of KEGG metabolic signatures using the AddModuleScore method of Seurat.

**Causal Network Analysis of Bulk RNAseq Data** To construct a causal relationship network between our key predictive genes and a discrete lineage variable we use an information theoretic method known as miic which learns graphical models from purely observational data, including the effects of unobserved latent variables[17]. Briefly, the algorithm starts from a complete graph, the method iteratively removes dispensable edges, by uncovering significant information contributions from indirect paths, and assesses edge-specific confidences from randomization of available data. The remaining edges are then oriented based on the signature of causality in observational data.

To perform miic profiling of the haemopedia RNAseq data[18] we first processed the bulk RNAseq library. Genes with log cpm values which did not have a log cpm value of at least 1 were omitted for further analysis. This led to a dataset with 17761 genes and 110 FACs sorted haematopoietic cell population. Data was normalised using the *voom* framework in limma[19]. Differential expression analysis and pathway based analyses were also performed using the limma framework with a Benjamin Hochberg correction applied to p-values to adjust the false positive rates for multiple comparisons[19]. We next took genes from the myeloid metafate signature as well as genes encoding enzymes and transporters that were (i) upregulated in the myeloid lineage, erythroid or lymphoid lineages and (ii) variably expressed in LSK hematopoietic progenitors using published datasets[14,18]. A full list of these genes is provided in Table S10. The expression patterns of these 230 genes within 104 bulk RNA seq profiles from the haemopedia database, as well as a discrete lineage variable for each sample were used as inputs into the miic web server[20] run on default parameter settings: https://miic.curie.fr/. The results of the network analysis are available at: https://miic.curie.fr/job_results.php?id=1MgzD2kVPrtuYH9vlagI

**Analysis of Sell and MetaFate expression in response to growth factor and inflammatory signals.** Single cell transcriptomics datasets of HSPCs following in vitro G-CSF stimulation, in vivo EPO stimulation, and PyMT induction of breast cancer were downloaded from the data archives of the original articles and imported into the Seurat R package. For each dataset the MetaFate signature score was computed using the AddModuleScore method of Seurat[21]. Differential expression analysis was performed using Seurats logistic regression method.

**Data and script accessibility**

**Code and data availability:** All data and code are available at https://github.com/TeamPerie/Cosgrove-et-al-2022.

**Supplementary Tables**

Table S1: Fluorescently labelled antibodies

Table S2: PCR amplification cycles for targeted barcode recovery at the RNA level

Table S3: **:** Oligo Sequences Used in This Study.

Table S3: Metadata for MetaFate experiments

Table S4: Differentially expressed genes between barcoded HSPCs

Table S5: Differentially expressed pathways between barcoded HSPCs

Table S6: Gene Signatures used in scRNAseq analysis

Table S7: MetaFate expression matrices

Table S8: Lentiviral barcoding count matrix for MPP transplantation study 3 weeks post-transplantation

Table S9: Lentiviral barcoding count matrix for Slam HSPC transplantation study 12 months post-transplantation

Table S10: Input Matrix for MIIC causal network analysis

Table S11: Lentiviral barcoding count matrix for G6PD-Tg MPPs 3 weeks post transplantation

Table S12: Lentiviral barcoding count matrix for WT MPPs 3 weeks post transplantation

| Antibody target | Clone | Conjugate | Manufacturer | Relevant Figure Pan | Dilution |
|---|---|---|---|---|---|
| CD45.1 | A20 | PE/BUV737 | BD Biosciences | S11 | 1:50 |
| CD45.2 | 104 | BV605 | Biolegend | S11 | 1:100 |
| Cd34 | Ram34 | E450 | Invitrogen | 4 | 1:50 |
| Ter119 | TER119 | PE-Cy7 | BD Biosciences | 1,3,4,5,S2,S9 | 1:100 |
| CD19 | 1D3 | APC-Cy7 | BD Biosciences | 1,3,4,5,S2,S9 | 1:100 |
| CD3 | ebio500A2 | PE | eBiosciences | 1,3,4,5 | 1:100 |
| CD11b | M1/70 | E450,PerCP-Cy5.5 | eBioscience | 1,3,4,5,S2,S9,S11,S12 | 1:500 |
| CD117 (C-Kit | 2B8 | APC/APC-Cy7 | BioLegend | 1,2,3,4,S2,S7,S9, S11,S12,S14 | 1:100 |
| CD117 (C-Kit | ACK2 | BV650 | BioLegend | | 1:100 |
| CD135 (Flt3) | A2F10 | PE | eBioscience | 2,S7 | 1:100 |
| CD135 (Flt3) | A2F10 | PE-Cy5 | Life technologies | | 1:100 |
| Sca1 | D7 | Pacific Blue /APC-Cy7 | BioLegend | 2 | 1:200 |
| Sca1 | E13-161.7 | Pacific Blue | Biolegend | 1,2,3,4,5,S7,S9,S11, S12,S14 | 1:200 |
| CD150 | TC15-12F12.2 | PE-Cy7/ PerCPCy5.5 | BioLegend | 1,2,3,4,5,S7,S9,S11, S12,S14 | 1:100 |
| Ter119 | TER119 | biotin | BD Biosciences | Ter enrichment | 1:100 |
| CD44 | IM7 | PE | BD Biosciences | 1,3,5,S2,S9,S12 | 1:100 |
| CD41 | MVVREG30 | BV510 | BD Biosciences | | 1:100 |
| CD62L | MEL-14 | PE / BV605 | Biolegend | 2,3,4,5,S11,S12 | 1:100 |
| Cd48 | HM48-1 | APC-Cy7/PerCPCy5.5 | BD Biosciences | 2,3,4,5,S7,S9,S11, S12,S14 | 1:100 |
| CD16/32 | 2.4G2 | FITC | BD Biosciences | 4 | 1:100 |
| Lin | CD3ε, clone 145-2C11 Ly-6G/Ly-6C, clone RB6 8C5; CD11b, clone M1/70; CD45R/B220, clone RA3-6B2; TER-11 | PE | Biolegend | 4 | 1:200 |
| Lin | CD3 (17A2), Ter-119 (Ter119),B220 (RA3-6B2),Gr-1 (RB8-6C5) | PE-Cy7 | Biolegend / BD biosciences | 2,S7 | 1:200 |

**Table S1: Fluorescently labelled antibodies**

| cDNA Input | Total cycles (10x protocol) |
|---|---|
| 0,25-25ng | 16-18 |
| 25-150ng | 14-16 |
| 150-500ng | 12-14 |
| 500-1000ng | 10-12 |
| 1000-1500ng | 8-10 |
| >1500ng | 7 |

Table S2. PCR amplification cycles for targeted barcode recovery at the RNA level

| **Genotyping** | | |
|---|---|---|
| BCM transge | MBR26 Fwd | AAGCACTTGCTCTCCCAAAGTCG |
| | CaggRev2inn | GTAACGCGGAACTCCATATATGGG |
| Rosa26 wt | MBR26 Fwd | AAGCACTTGCTCTCCCAAAGTCG |
| | MBR 26 Rev | TCCCATTTTCCTTATTTGCCCT |
| Cre transgen | oIMR1084 | GCG GTC TGG CAG TAA AAA CTA TC |
| | oIMR1085 | GTG AAA CAG CAT TGC TGT CAC TT |
| Cre WT | oIMR7338 | CTA GGC CAC AGA ATT GAA AGA TCT |
| | oIMR7339 | GTA GGT GGA AAT TCT AGC ATC ATC C |

| **DRAG DNA Tagging PCR** | |
|---|---|
| Preamp forward | ACTCACTATAGGGAGACGCGTGTTACC |
| Preamp reverse | GACACGCTGAACTTGTGGCCGTTTA |
| M1 tag forward | ACACTCTTTCCCTACACGACGCTCTTCCGATCNNNNNNNNNNNNCCTCGAGGTCATCGAAGTATCAAG |
| Illumina forward seq (R1) | ACACTCTTTCCCTACACGACGCTCTTCCGA*T*C (*= Phosphorothioate bond) |

| | |
|---|---|
| M1 rev Read2 | AGTTCAGACGTGTGCTCTTCCGATC CAGCTCGACCAGGATG*G*G |
| P5 forward | AATGATACGGCGACCACCGAGATCTACACTCTTTCCCTACACGACGCTCTTCCGATC |
| P7 index rev | CAAGCAGAAGACGGCATACGAGAT**XXXXXXX**GTGACTGGAGTTCAGACGTGTGCTCTTCCGATC |
| **Primers for targeted amplification of lineage barcodes from 10X 3' libraries** | |
| Pre-amp Partial read1 F | CTACACGACGCTCTTCCGATCT |
| PreAmp + read2 R | GTGACTGGAGTTCAGACGTGTGCTCTTCCGATCTactcactatagggagacgcgtgttACC |
| Sample index PCR P5-Rd1 | AATGATACGGCGACCACCGAGATCTACACTCTTTCCCTACACGACGCTCTTCCGATCT |
| Sample index PCR P7 Sample index | caagcagaagacggcatacgagatNNNNNNNgtgactggagttcagacgtgtgctcttccgatc |
| **Primers used in Lentiviral Barcoding** | |
| top-LIB | 5'TGCTGCCGTCAACTAGAACA-3' |
| bot-LIB | 5'GATCTCGAATCAGGCGCTTA-3' |
| 4 bp plate index forward | 5'ACACTCTTTCCCTACACGACGCTCTTCCGATCTNNNNCTAGAACACTCGAGATCAG3' |
| 4 bp plate index reverse | 5'GTGACTGGAGTTCAGACGTGTGCTCTTCCGATCGATCTCGAATCAGGCGCTTA3' |
| P5 | 5'AATGATACGGCGACCACCGAGATCTACACTCTTTCCCTACACGACGCTCTTCCGATCT3' |
| P7 | 5'CAAGCAGAAGACGGCATACGAGANNNNNNNGTGACTGGAGTTCAGACGTGCT CTTCCGATC3' |

**Table S3:** Oligo Sequences Used in This Study. Complete list of the i7 indexes in[8]. All oligo's were ordered at IDT with HPLC purified grade.

### Supplementary Figures

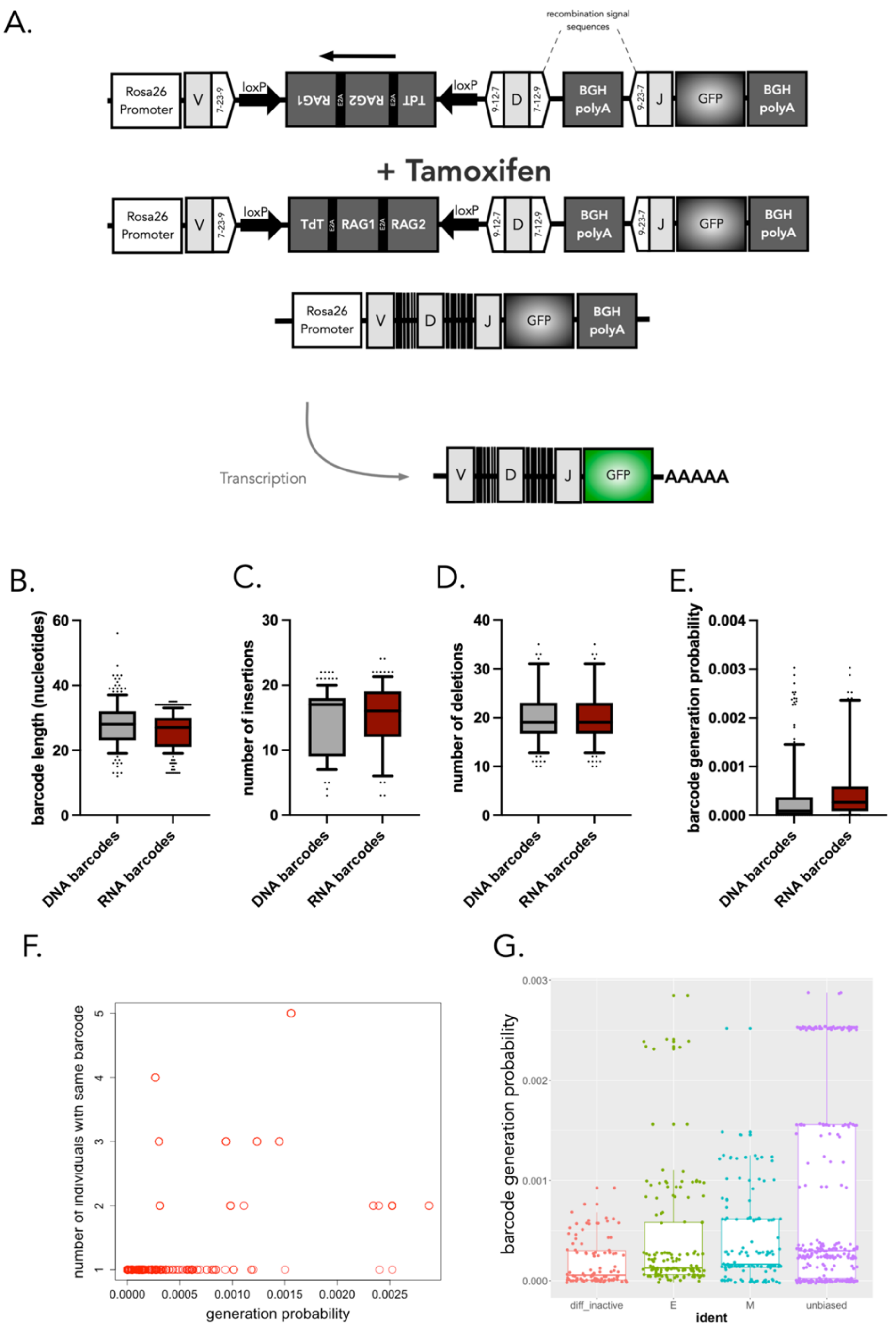

**Figure S1. Overview and QC of the DRAG barcoding technology.** (A)Description of the DRAG cassette, as inserted into the Rosa 26 locus before and after induction. DRAG recombination is induced by Cre activity and resulting barcode sequences are used for lineage tracing. The DRAG system has been designed such that upon CRE induction, a segment between two loxP sites is inverted, leading to the expression of both the RAG1 and 2 enzymes and Terminal deoxynucleotidyl transferase (TdT). Upon such expression, recognition of recombination signal sequences (RSSs) within the DRAG cassette by the RAG1/2 complex leads to recombination of the synthetic V-, D- and J-segments, with diversity being generated both by nucleotide deletion and TdT-mediated N-addition. Notably, as the RAG/TdT cassette and RSSs are spliced out during this recombination step, further recombination of the DRAG locus is prevented, and any generated VDJ sequence is thus stable over time. Finally, recombination of the DRAG locus results in the removal of a BGH polyA site that precludes GFP expression in the native DRAG configuration, allowing one to identify barcode+ cells by flow cytometry or imaging. To allow in situ barcode generation, DRAG mice were crossed with ubiquitously expressed tamoxifen-dependent Cre (RosaCre-ER™) mice, to obtain the heterozygous RosaCre$^{+/-}$ DRAG$^{+/-}$ mice used in all experiments. (B) Length in nucleotide of DRAG barcodes detected from RNA or DNA (C) Numbers of insertions in DNA and RNA retrieved DRAG barcodes (D) Numbers of deletions in DNA and RNA retrieved DRAG barcodes (E) Distributions of barcode generation probabilities for RNA and DNA barcodes (F) A dotplot showing the relationship between repeat use barcodes (barcodes that occur in more than 1 mouse), and their barcode generation probability. (G) The distribution of predicted barcode generation probabilities across different classes of lineage-biased barcode categories. Data from n = 5 mice, same experiments as Figure 1.

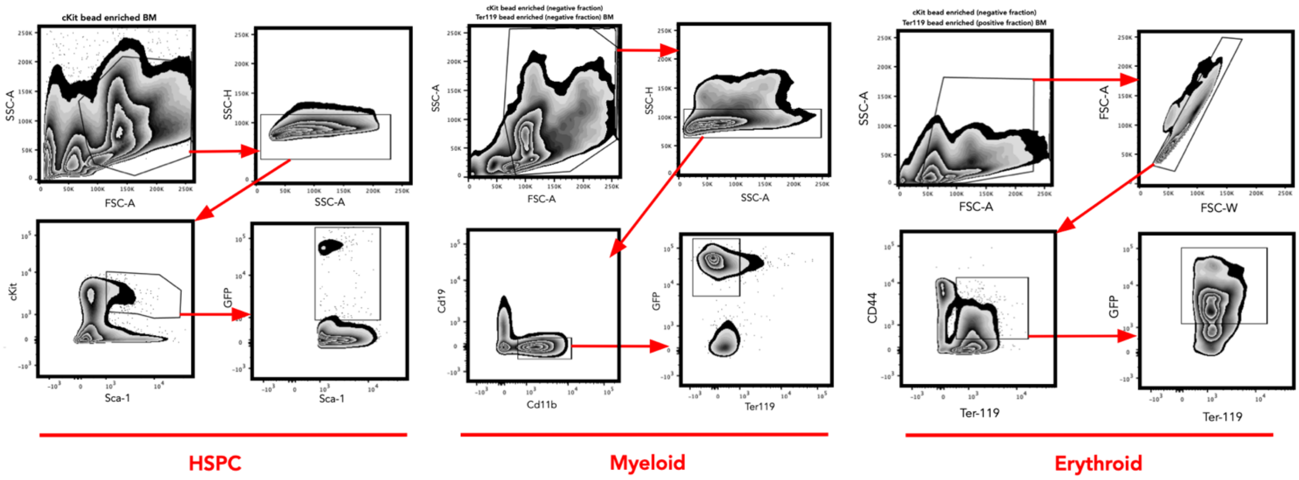

**Figure S2. Gating strategy for MetaFate experiment.** HSPCs are classified as cKit+ Sca1+ GFP+, myeloid cells are classified as CD19- Ter119- and Cd11b+ GFP+ , and nucleated erythroid cells are classified as CD19- Cd11b- and Ter119+ Cd44+ GFP+.

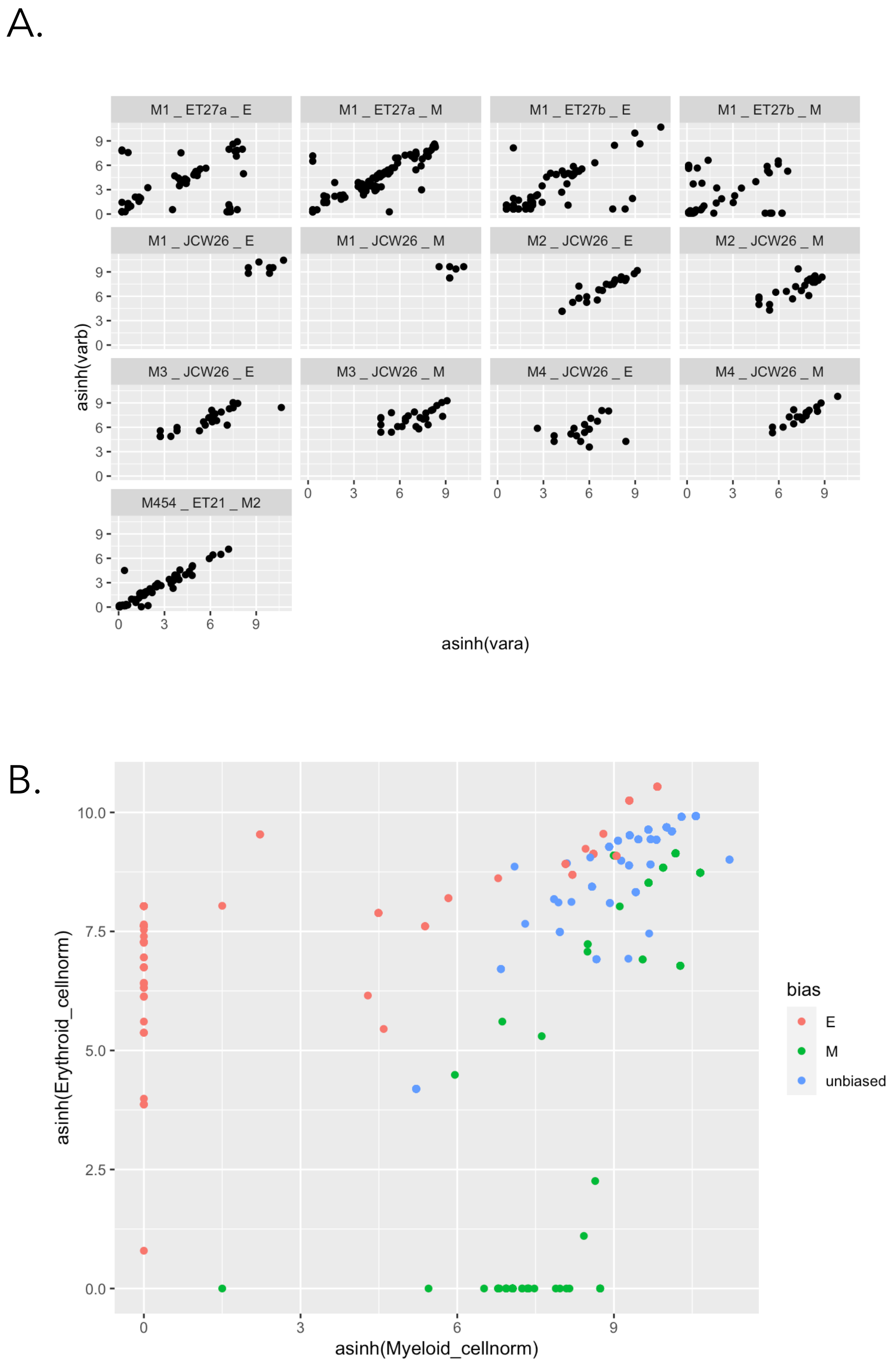

**Figure S3: Bulk DNA barcode QCs. (A)** the normalized and hyperbolic arcsin transformed read counts in one PCR duplicate (vara) compared to the second PCR duplicate (varb). Each plot is a distinct sample, and each point represents 1 barcode. Each panel represents an individual sample with the sample name shown above the plot labelled as mouse_experiment code_cell type with *E* corresponding to erythrocytes and *M* to myeloid cells. (B) The normalized and hyperbolic arcsin transformed read counts present myeloid cells versus erythroid cells for each biased categories (E in red for erythroid biased, M in green for myeloid biased and unbiased barcodes in blue). Each point represents 1 barcode, pool from 5 mice.

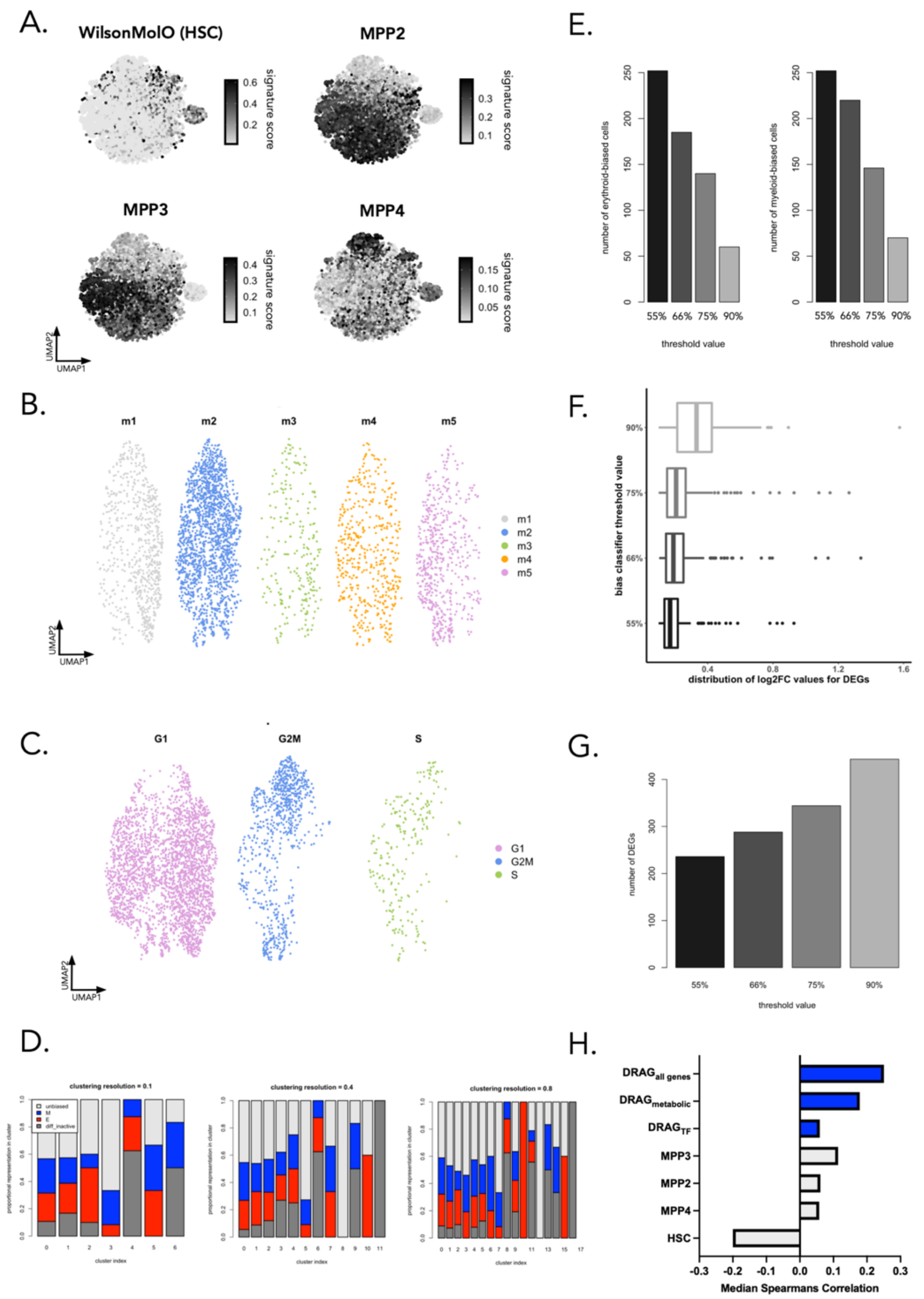

**Figure S4. MetaFate data QC. Show QC plots for 10X and for DNA barcodes.** (A) gene expression signatures for HSC (from Wilson et al[12]) and MPP subsets (from Sommerkamp et al[13] and Pietras et al.[22]) overlaid onto the UMAP representation of the data. These signatures were used to positioning cell subsets in Figure 1.B. (B) the distribution of cells from each biological replicate (individual mouse) onto the reference UMAP embedding of the data. (C ) distribution of cells in G1/G2M/S cell cycle phases in our data as determined using the cyclone method[23] implemented in the scran package. (D) The distribution of E/M/unbiased/differentiation inactive barcoded cells across unsupervised clusters for different clustering resolutions (0.1,0.4 and 0.8). This analysis was performed using Seurat's default implementation of the Louvain clustering method. (E) The number of cells belonging to each lineage bias class across a range of different lineage-bias threshold values (F) The distribution of log2fc changes in gene expression for differentially expressed genes using different lineage bias classifier threshold values. (G) The number of DEGs detected for the myeloid biased barcoded cells across a range of lineage-bias classifier values. (H) 4-fold validation of the metafate signature generation pipeline. In this analysis we partitioned barcoded cells into 4 folds that were iteratively used for training and testing. Signatures were obtained by performing differential expression between barcoded progenitor subsets and significantly genes upregulated in myeloid-biased cells were grouped into one of 3 signatures (DRAG-allgenes, DRAG-metabolic, and DRAG-TF). For each iteration (200 in total), the Spearman's correlation between gene signature and barcode derived myeloid bias was assessed and compared against published signatures from Wilson et al[12], and Pietras et al[22].

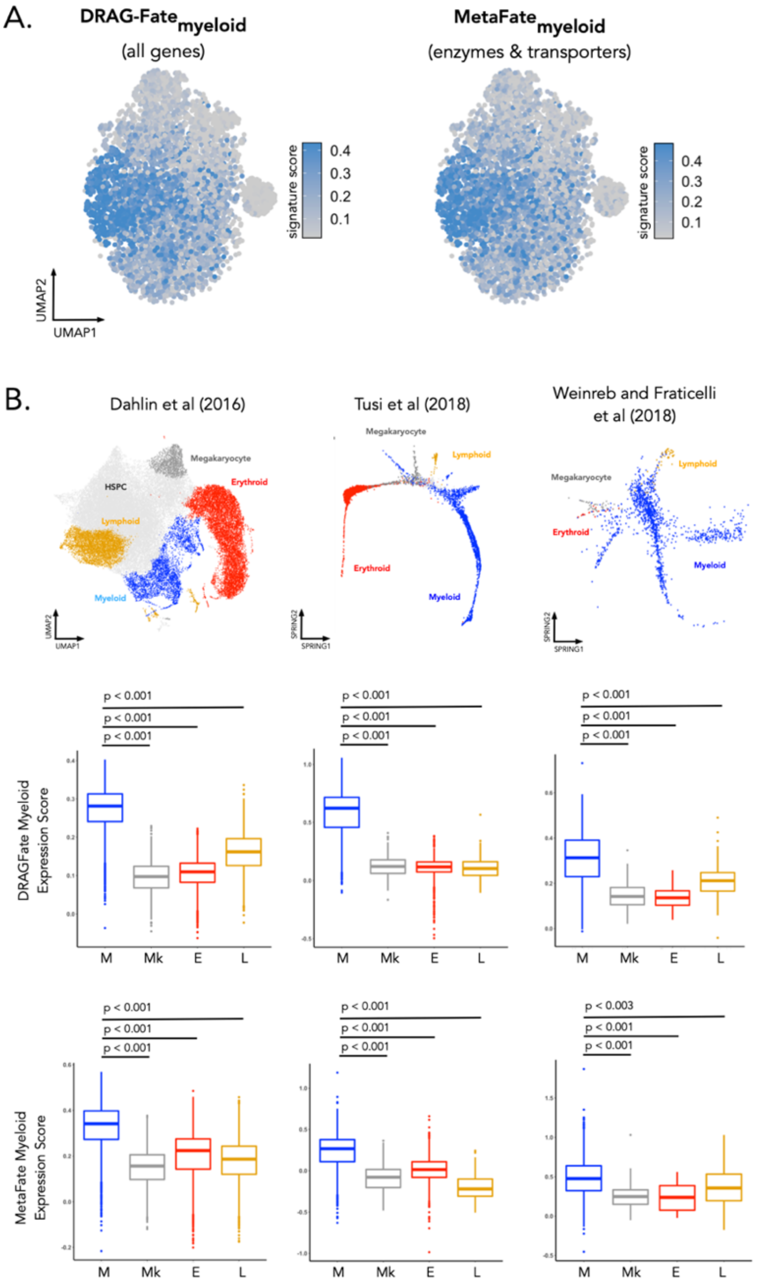

**Figure S5. Expression of the Fate-myeloid and metafate-myeloid signatures on previously published scRNAseq datasets.**(A) All genes upregulated in myeloid-biased barcoded cells compared to erythroid and differentiation inactive-barcoded cells form a gene-signature called DRAGFate-Myeloid. The subset of genes from the DRAGFate-myeloid signature relating to cellular metabolism form the MetaFate-myeloid signature. The signature score corresponds to the average expression values of these gene sets for each cell and is projected onto the UMAP visualisation of the data. (B) Benchmarking of signatures on published dataset. Here we show the expression of the DRAGFate myeloid and MetaFate myeloid signatures on published scRNAseq datasets. In these analyses we used cell-type definitions provided in the original article and signature scores were calculated using the AddModuleScore() function in Seurat. Dahlin et al (left) comprises 44,802 cKit+ and cKit+ Sca1+ hematopoietic progenitors[14]. Cell clustering and supervised assignment of cluster identity were taken from Wolf et al[15]. The dataset from Tusi et al (middle) comprises 4,763 cKit+ progenitors[24]. To annotate this dataset we performed unsupervised clustering of the data and supervised annotation using lineage-specific markers provided in supplementary table 1 of the original article[24]. Weinreb and Fraticelli et al.[25] (right) comprises 28,249 cKit+ and cKit+ Sca1+ progenitors that were lentivirally barcoded and cultured for 2 days in vitro. In this analysis cells were classified as M, Mk, E or L biased on whether the majority of cells within that clone are found within the lymphoid, myeloid, megakaryocyte and lymphoid clusters defined in the original publication. Cells that were undifferentiated or that could not be assigned to a lineage in this manner were excluded from further analysis, leaving a total of 1,518 barcoded cells for final comparisons. For all statistical tests we use a pairwise Mann-Whitney test corrected for multiple comparisons using the Benjamini-Hochberg method. M = myeloid (blue) ; Mk = Megakaryocyte (grey) ; E = erythroid (red) ; L = lymphoid (orange).

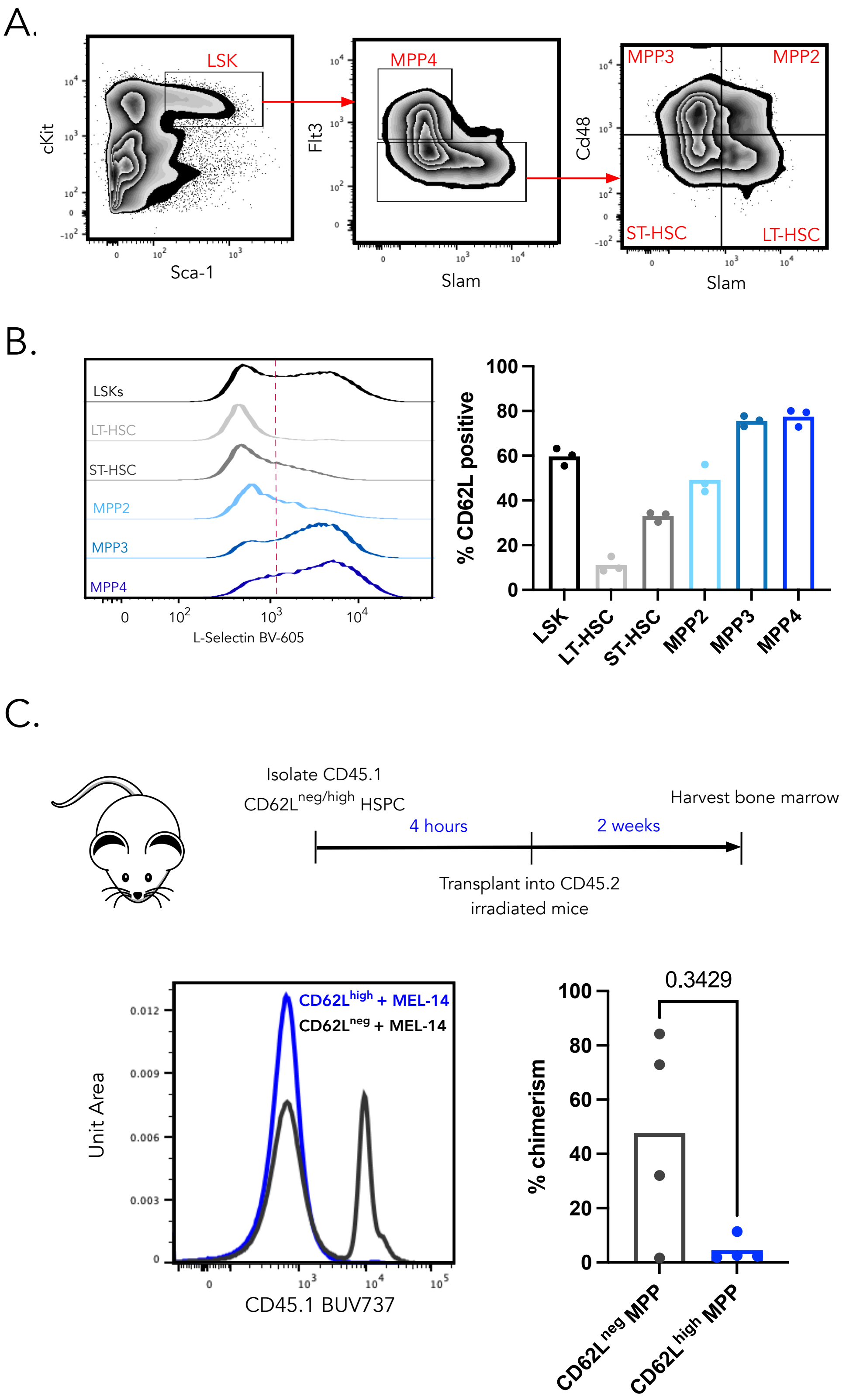

**Figure S6. CD62L expression amongst HSPC subsets and Anti-CD62L (Clone MEL-14) impairs the engraftment of CD62L-expressing HSPCs** (A) Gating strategy to purify different HSPC subsets (B) L-selectin expression on different HSPC subsets as quantified by flow cytometry (C top) Overview of the experimental timeline for the transplantation of CD62L$^{neg}$ and $^{high}$ MPPs (C bottom) % Chimerism in cKit- bone marrow cells for the CD62L$^{neg/high}$ transplanted HSPCs. N = 4 mice per condition.

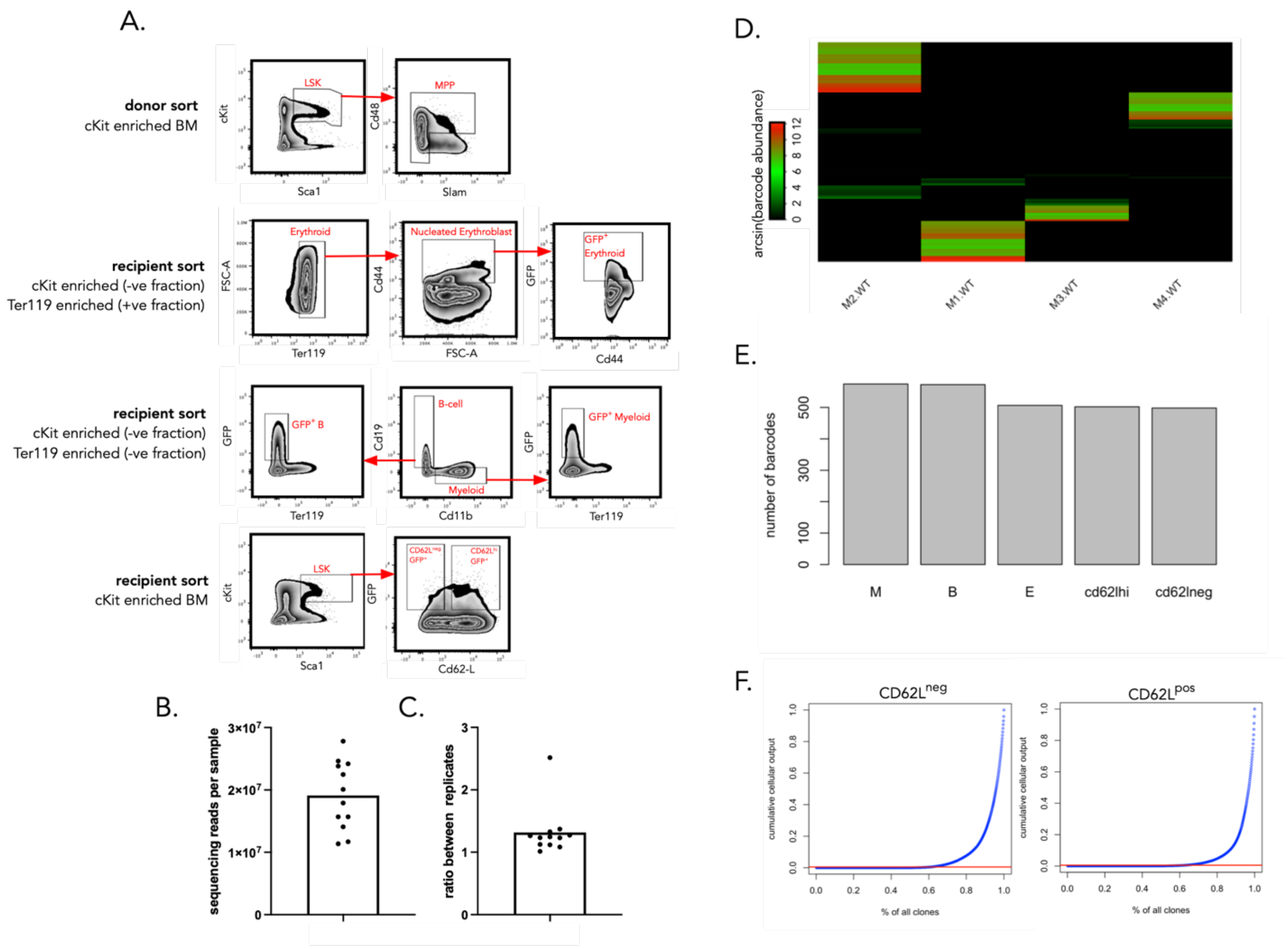

**Figure S7. lentiviral barcoding of CD62L$^{neg/high}$ MPPs at 3 weeks post-transplantation** (A) overview of the flow cytometry gating strategy used to purify cells for lentiviral barcoding analysis of Figure 5. (B) the number of sequencing reads for each sample in the barcoding analysis. Each point represents a bulk sorted cell population. N = 4 mice (C) The ratio between technical replicates where each point represents a bulk sorted cell population (D) Heatmaps showing the distribution of barcode abundance in reads across mice after data QC and filtering to ensure that we do not have significant numbers of barcodes shared between mice injected with the same batch, these shared barcodes are more likely to label more than 1 cell. Data are hyperbolic arcsin transformed reads. (E) Barcode diversity, as the number of unique barcodes, in QC processed samples pooled from 4 mice. (F) Thresholding strategy to define CD62L$^{neg/high}$ barcodes. Each point represents a single barcode and the red line represents a threshold value of 0.5%.

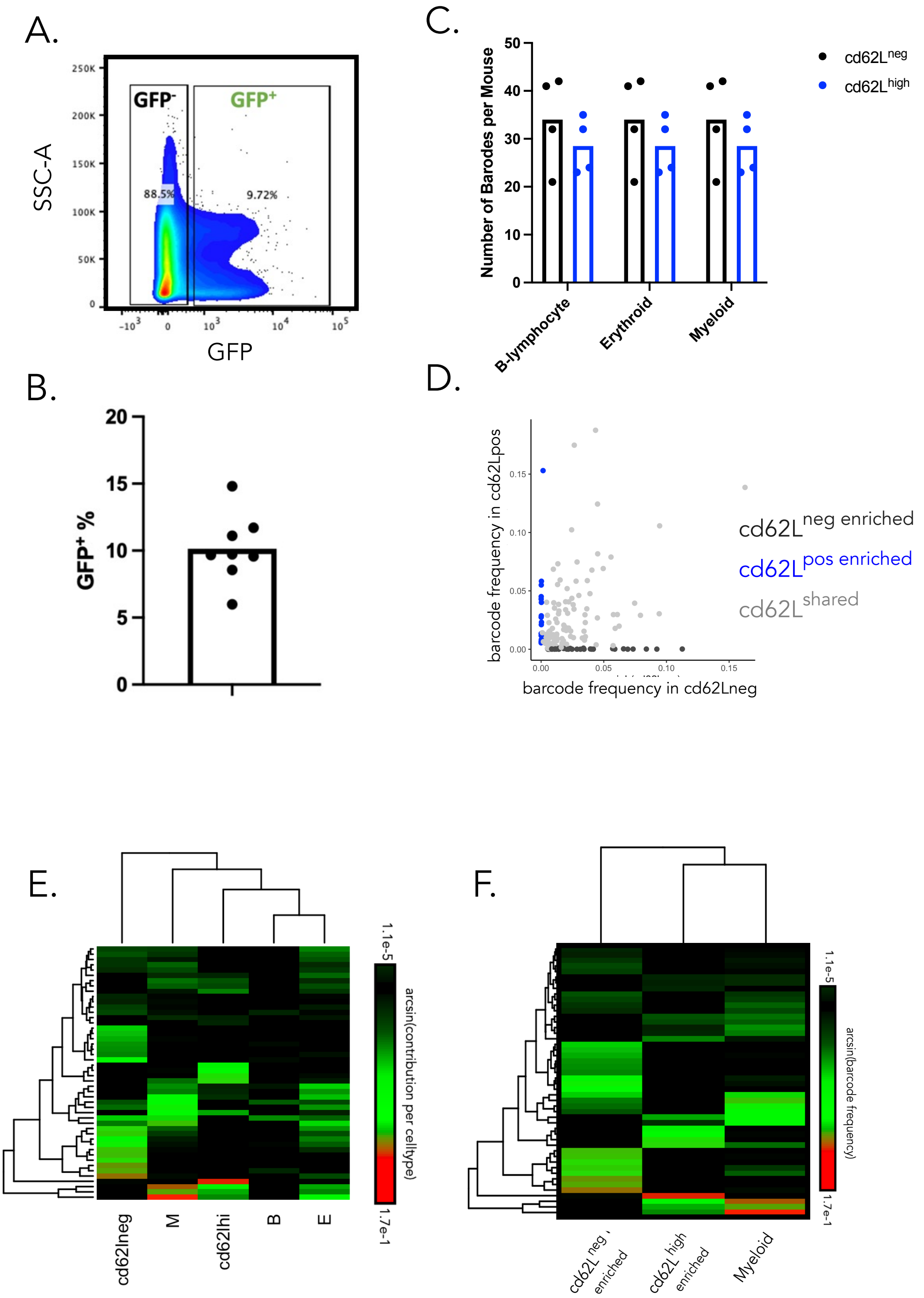

**Figure S8. QC and supporting analyses for lentiviral barcoding experiments.** (A-B) Percentage chimerism observed in lentiviral experiments. Data pooled across all mice (n = 8). (C) Barcode diversity scores in myeloid (*M*), erythroid (*E*) and B-lymphocyte (*B*) lineages for CD62L$^{neg/high}$ MPPs (black/blue points respectively). No statistically significant differences were observed between CD62L$^{neg/high}$ conditions for any lineage. Each point represents a single mouse and statistical comparisons were made using a paired Wilcoxon-Signed Rank test. N = 4 mice. (D) read abundance of barcode in the CD62L$^{neg\ enriched}$ and the CD62L$^{high\ enriched}$ HSPC (LSK ) fraction, each dot is a barcode and the axis are transformed using the hyperbolic arcsin function. Barcodes that had more than 95% of its reads in either the CD62L$^{neg}$ or the CD62L$^{high}$ MPPs were classified as CD62L$^{shared}$ (light grey ; 124 barcodes), CD62L$^{neg\ enriched}$ (dark grey ; 30 barcodes) and CD62L$^{high\ enriched}$ (blue ; 18 barcodes) MPPs (E) Unsupervised clustering and heatmap visualisation of CD62L$^{neg/high}$ barcodes across all samples. Data are pooled from 4 mice. (F) Unsupervised clustering (using the Euclidean distance) and heatmap visualisation showing hyperbolic arcsin transformed barcode abundances for CD62L$^{neg\ enriched}$ and CD62L$^{high\ enriched}$ barcodes. Every row is a barcodes and every column is a cell type. All Boxplots represent the median and interquartile range with whiskers extending to the minimum and maximum values. N= 4 mice.

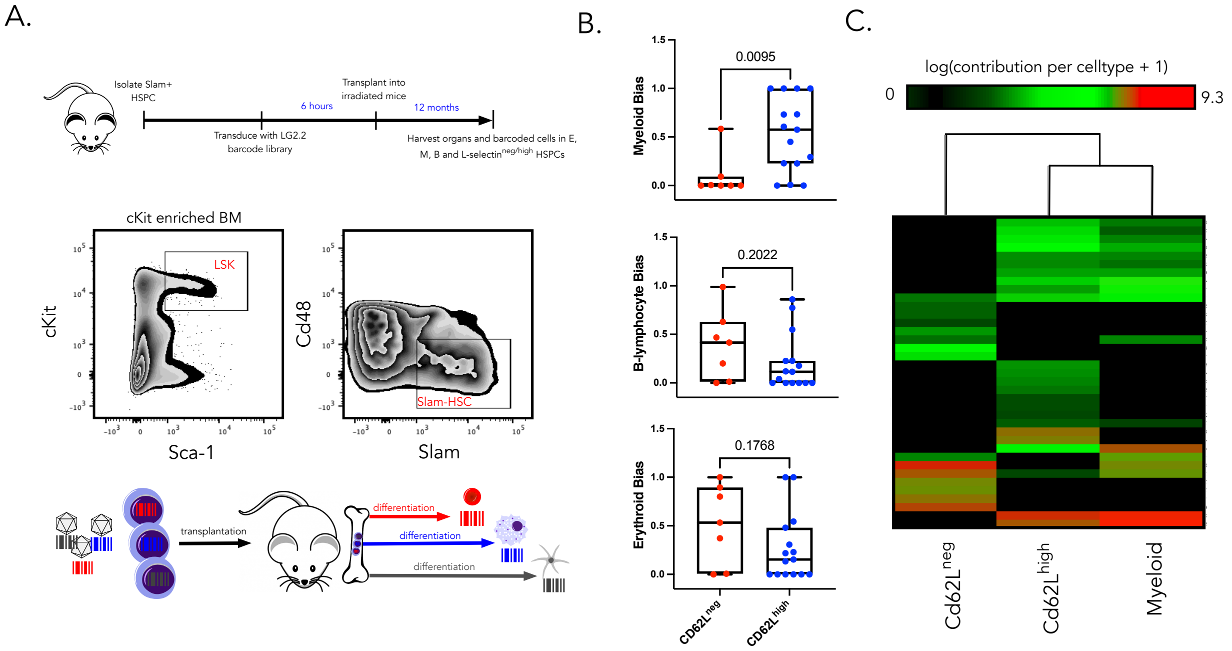

**Figure S9. The CD62L$^{high}$ MPP compartment reconstitutes the myeloid compartment following Slam HSC bone marrow transplantation** (A) cKit+ Sca1+ CD150+ Cd48- HSPCs were purified from donor mice by FACS and infected with the LG2.2 lentiviral barcoding library. Transduced cells were transplanted into 4 sublethally irradiated (6Gy) recipient mice and 12 months later CD62L$^{neg/high}$ MPPs, CD19$^{+}$ B cells, CD44$^{+}$ Ter119$^{+}$ erythrocytes, and CD11b myeloid cells from the bone marrow and were processed for targeted sequencing of lineage barcodes. (B) Lineage-biases of CD62L$^{neg\ only}$ or CD62L$^{high\ only}$ MPPs for the myeloid, erythroid and B-lymphocyte lineages. Data is pooled from 4 mice, and each point represents a unique barcode. Statistical significance was assessed using a Mann-Whitney test (C) Unsupervised clustering and heatmap visualisation of barcodes that occur in either CD62L$^{neg\ only}$ or CD62L$^{high\ only}$ MPPs.

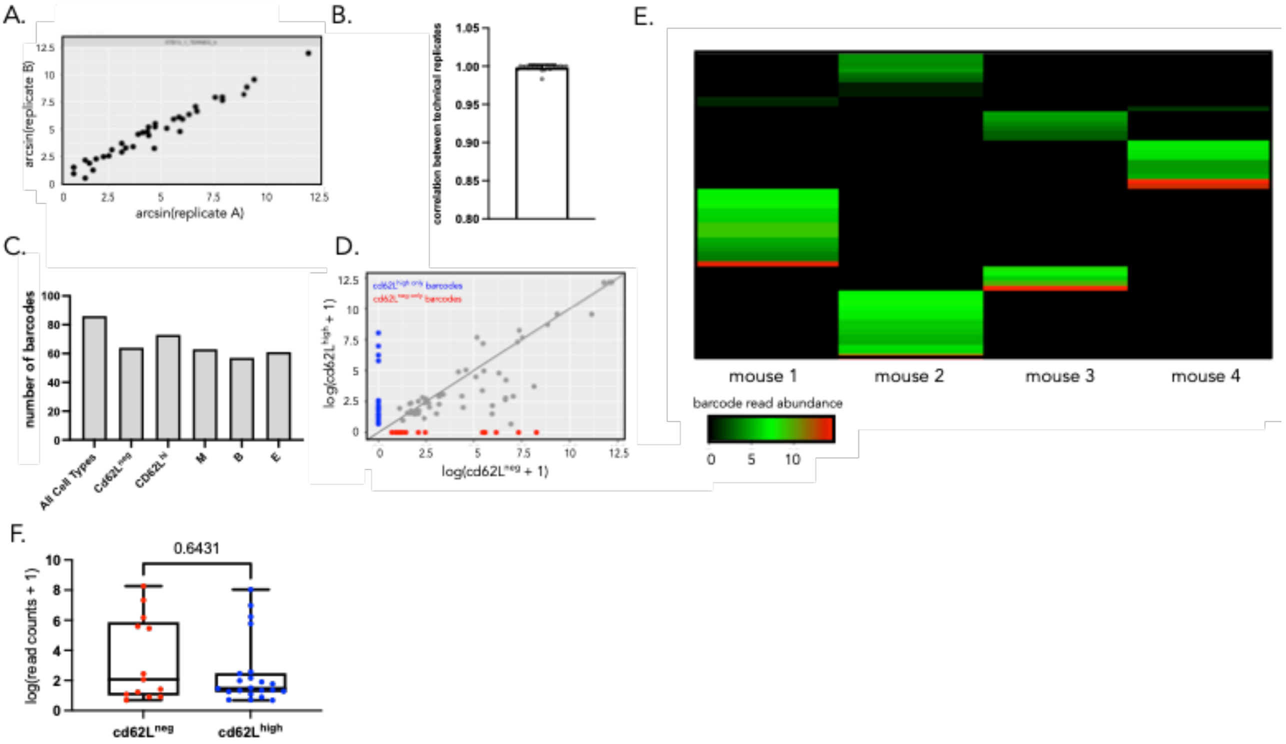

**Figure S10. QC for lentiviral barcoding of Slam HSCs**. (A) An example scatter plot showing the consistency of technical replicates – each point represents a distinct barcode (B) a barplot showing the Pearsons correlation of barcode abundances across technical replicates for all samples in our dataset. Each point represents a FACs sorted bulk population of cells. (C) The number of unique barcodes detected per cell type in the lentiviral barcoding dataset. Data pooled from 4 mice (D) log barcode abundance in CD62L$^{neg}$ and CD62L$^{high}$ MPPs. CD62L$^{shared}$ ,CD62L$^{neg\ only}$ and CD62L$^{high\ only}$ barcodes are highlighted in grey,red and blue respectively. Data pooled from 4 mice. (E) Heatmap to show the frequency of repeat-use barcodes across mice. N = 4 mice. (F) log distribution of read counts for CD62L$^{neg\ only}$ and CD62L$^{high\ only}$ barcodes. Statistical significance was assessed using a Mann-Whitney test.

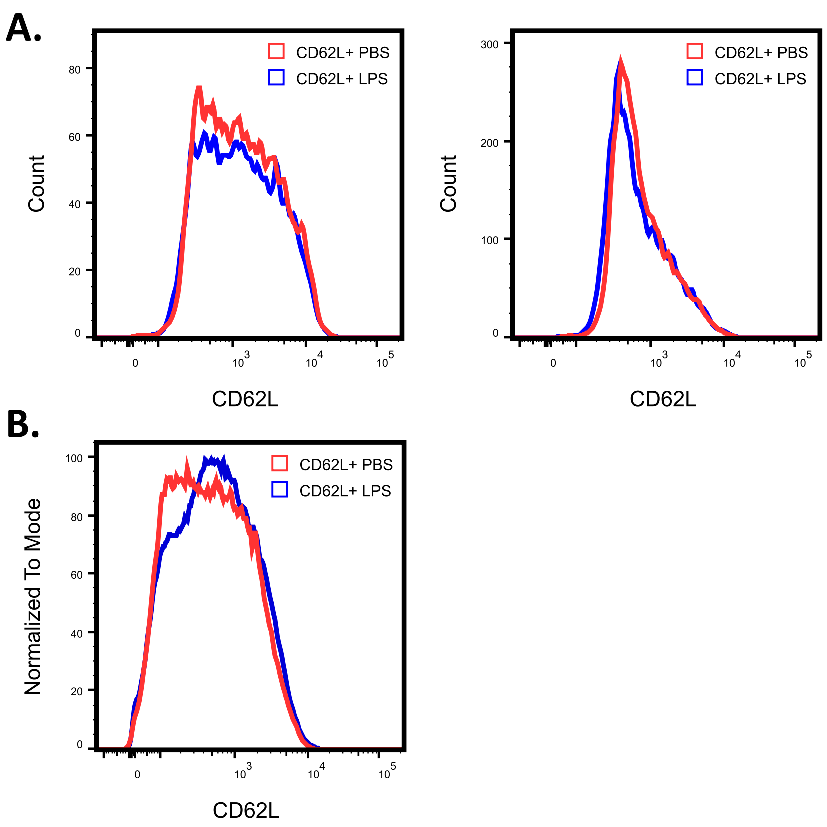

**Figure S11. CD62L expression after LPS-exposure. (**A) CD62L protein expression via flow cytometry, after treating FACS-sorted Cd62L$^{high}$ and Cd62L$^{neg}$ MPPs for 24 hours ex vivo with PBS (blue) or 200 ng/mL of LPS (red). The panels display one representative experiment, n = 2 in triplicate. Gating as in S7A to obtained MPPs. (B) CD62L protein expression via flow cytometry on MPPs of the bone marrow 72 hours after 2 I.P injections of LPS (35ug/mouse, in red) or PBS (blue) as in Figure 2F. The panel represent the 3 mice/group, representative of 2 experiments. Gating as in S7A to obtained MPPs.

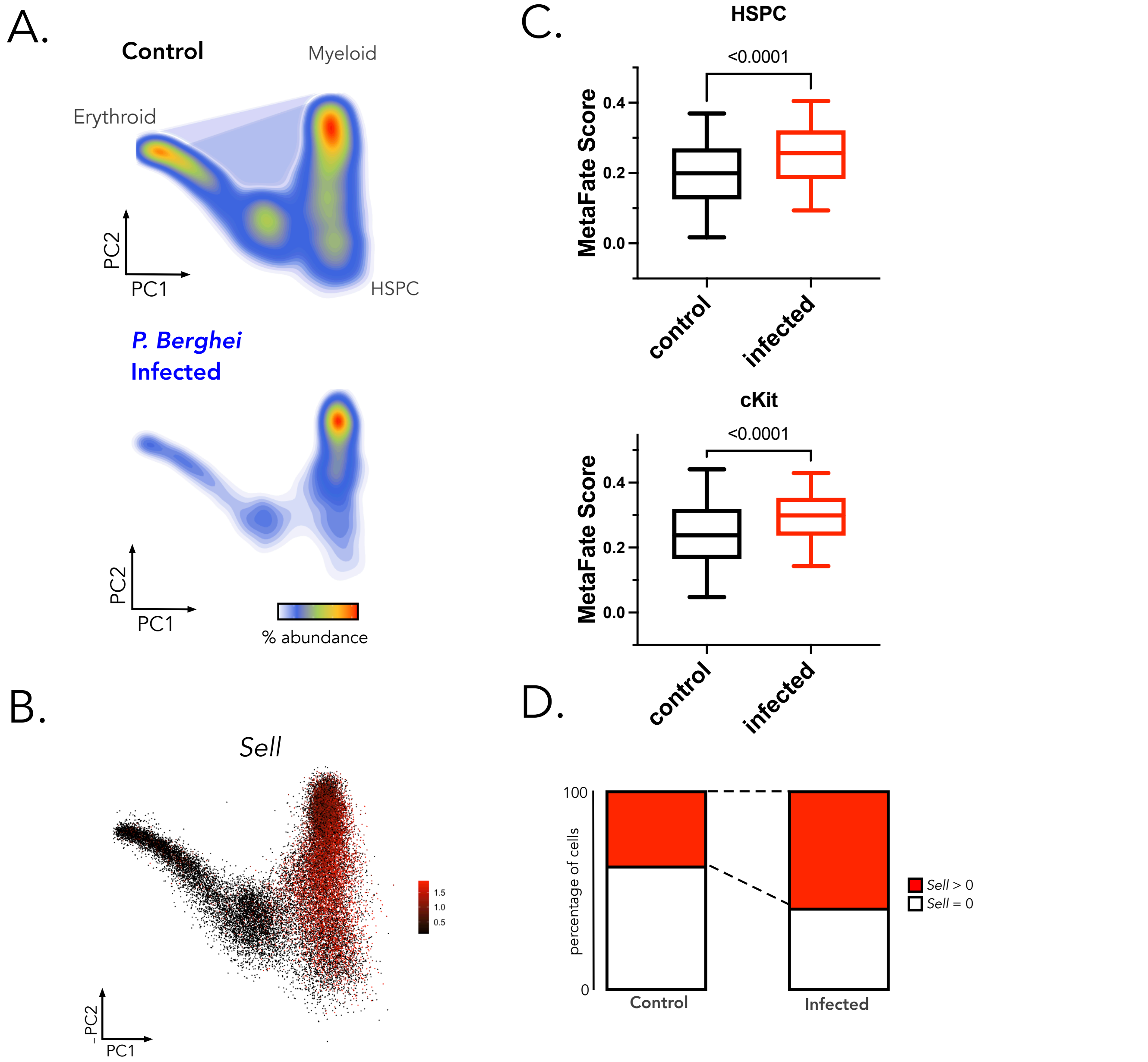

**Figure S12. Sell and MetaFate expression following plasmodium infection.** (A) scRNAseq data reanalysed from[26], where mice were treated with vehicle control or *P. berghei*. 7 days post infection cKit+ progenitors were purified from the bone marrow of each group and processed for scRNAseq profiling. Data are represented using a density projection of the cell abundances on to a PCA embedding of the data. Cell type annotations of the data were taken from the original publication. Control samples have 14193 cells and infected samples have 13905 cells. (B) *Sell* (gene encoding CD62L) normalised gene expression projected onto the PCA embedding of the data. (C top) Boxplot showing MetaFate signature expression score in HSPCs (defined as primitive HSPCs in the original article) from control (black) and infected (red) mice. Pairwise comparisons were made using a Students T-test. Boxplot showing mean and sd over cells? (C bottom) Same as F but for all cKit+ hematopoietic progenitors from control and infected mice. (D) The proportion of cells in control and infected mice that have non-zero expression of *Sell*. Boxplots represent the interquartile range and median values and whiskers represent the 5th and 95th percentile of the data.

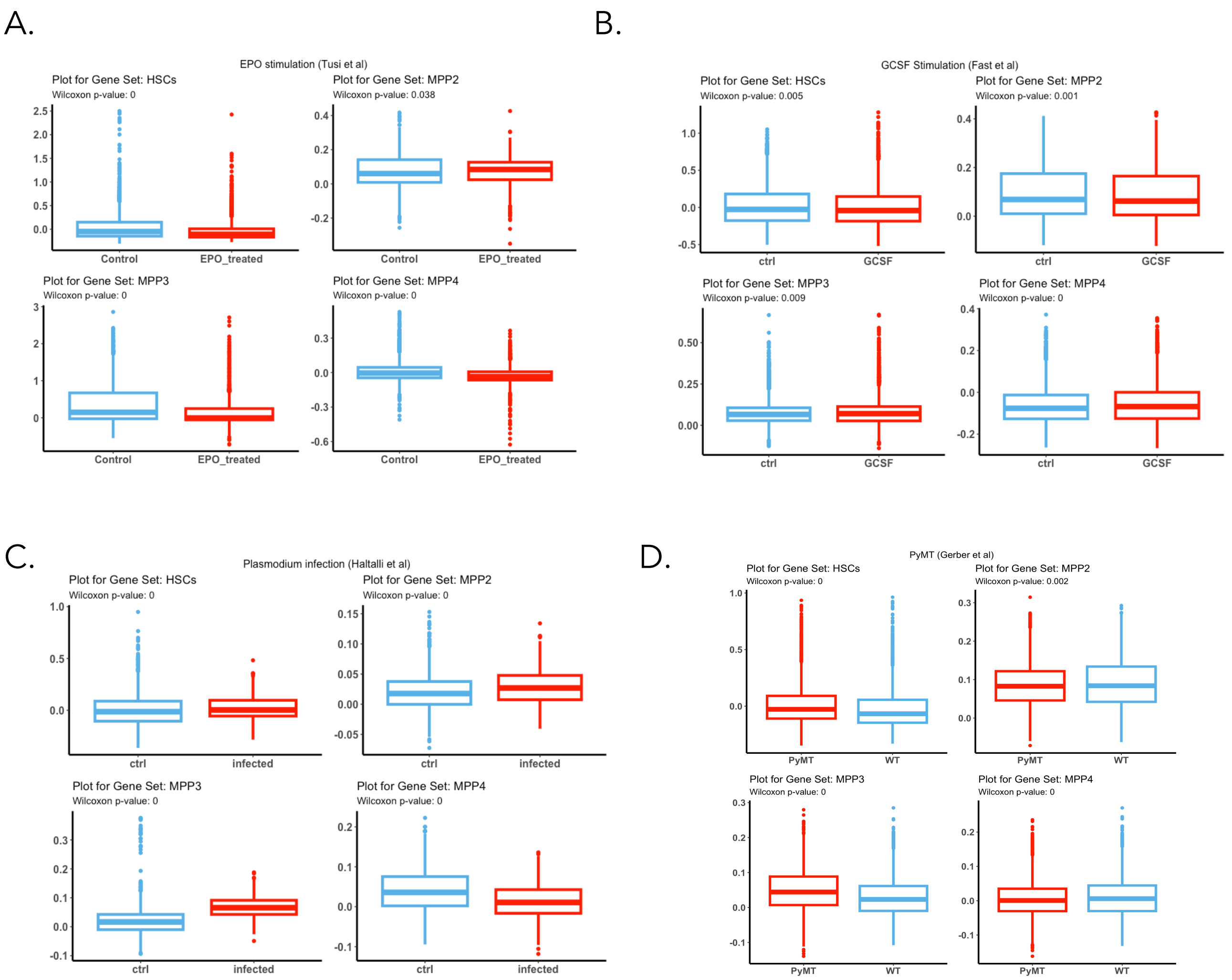

**Figure S13. HSC and MPP signature expression in inflammatory and cancer model.** (A) Boxplot showing HSC expression score from[12] and MPP2-4 signature expression score from[13] in individual scRNAseq data of HSPCs from EPO stimulated mice[27]. (B) Same from HSPCs stimulated ex vivo with G-CSF[28]. (C) Same from *P. berghei*. Infected mice[26] (D) Same from PyMT breast cancer model[29]. Control in blue and treated or infected in red. Pairwise comparisons were made using a Students T-test. Boxplots represent the interquartile range and median values and whiskers represent the 5th and 95th percentile of the data.

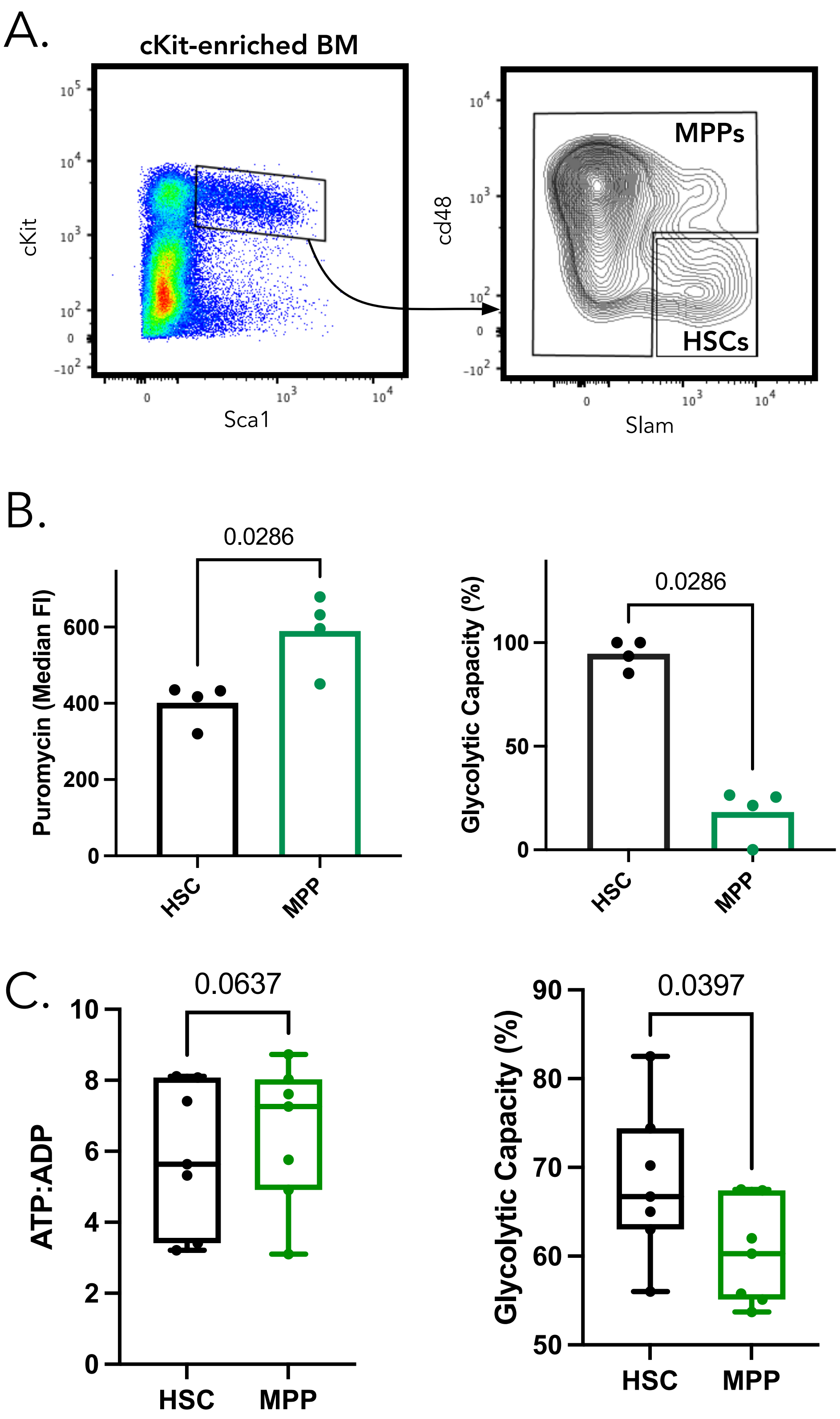

**Figure S14. Gating Strategy to define MPP versus HSC subpopulations and benchmarking and validation of metabolic profiling strategy**. (A) Gating strategy to compare the metabolic properties of HSCs and MPPs using SCENITH and SPICE-Met (B) SCENITH profiling of HSPCs gated as in Figure 1D, each point represents a unique mouse and N=4 mice. Statistical comparisons were made using a Mann-Whitney test. (C) SPICE-Met profiling of HSPCs, each point represents a unique mouse and N=7 mice. Statistical comparisons were made using a paired T-test. Data was pooled from 2 independent experiments.

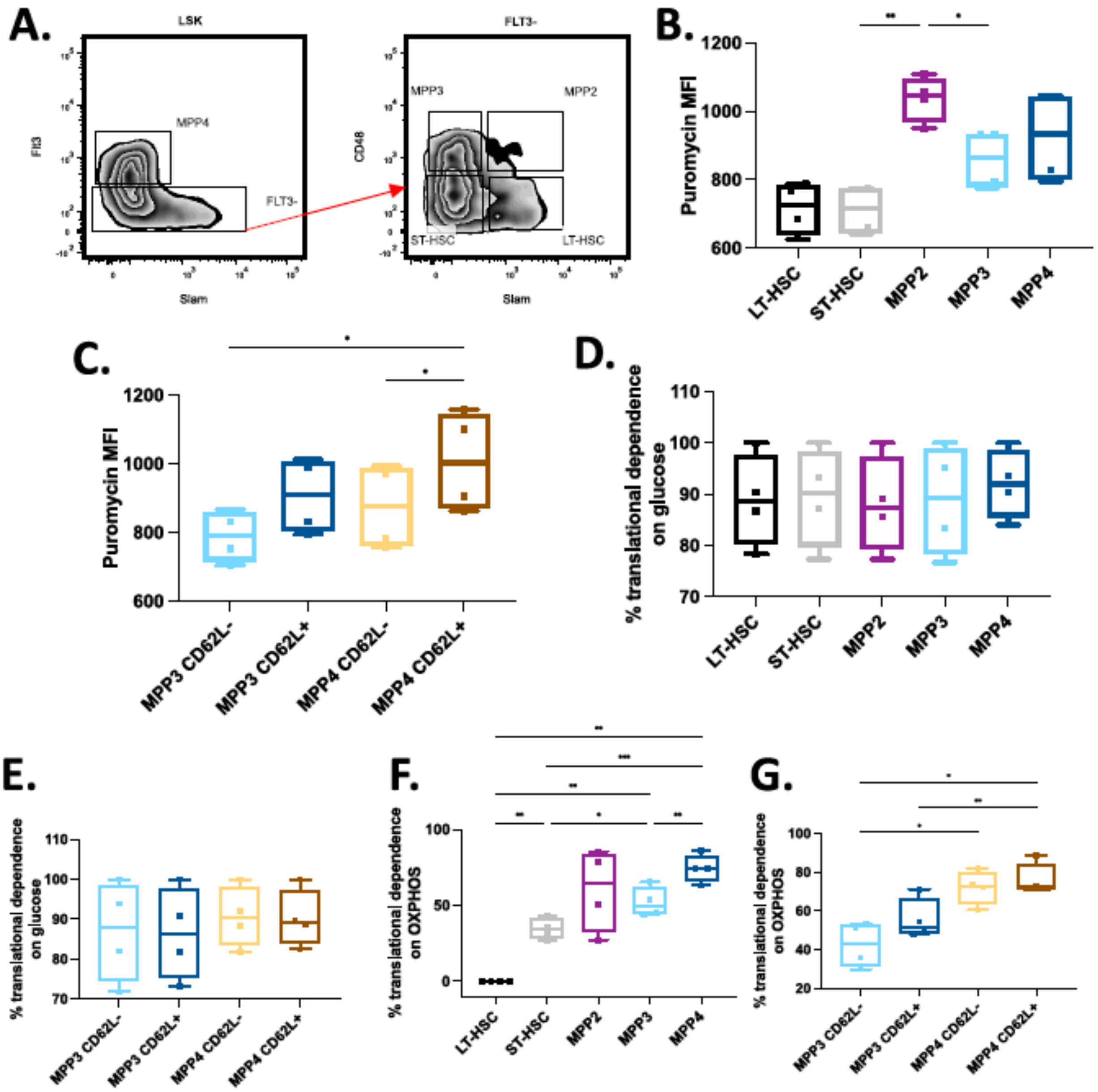

**Figure S15: SCENITH analysis on isolated MPPs.** (A) Gating strategy to isolate LT-HSC, ST-HSC and the different MPPs subsets. (B-C) SCENITH profiling of freshly isolated LSK, gated accordingly to (A). While (B) compares the 5 subsets, in (C) the MPP3 and MPP4 populations are separated accordingly to their CD62L expression. (D-E). Glucose dependence, measured via SCENITH. While (D) display the 5 subsets, (E) show only MPP3 and MPP4, separated based on their CD62L expression. (F-G) Oxidative phosphorylation dependence, measured via SCENITH. While (F) display the 5 subsets, (G) show only MPP3 and MPP4, separated based on their CD62L expression. Statistical comparisons were made using a Mann-Whitney test, p value *<0.05, **<0.01, ***<0.001, 4 independent mice.

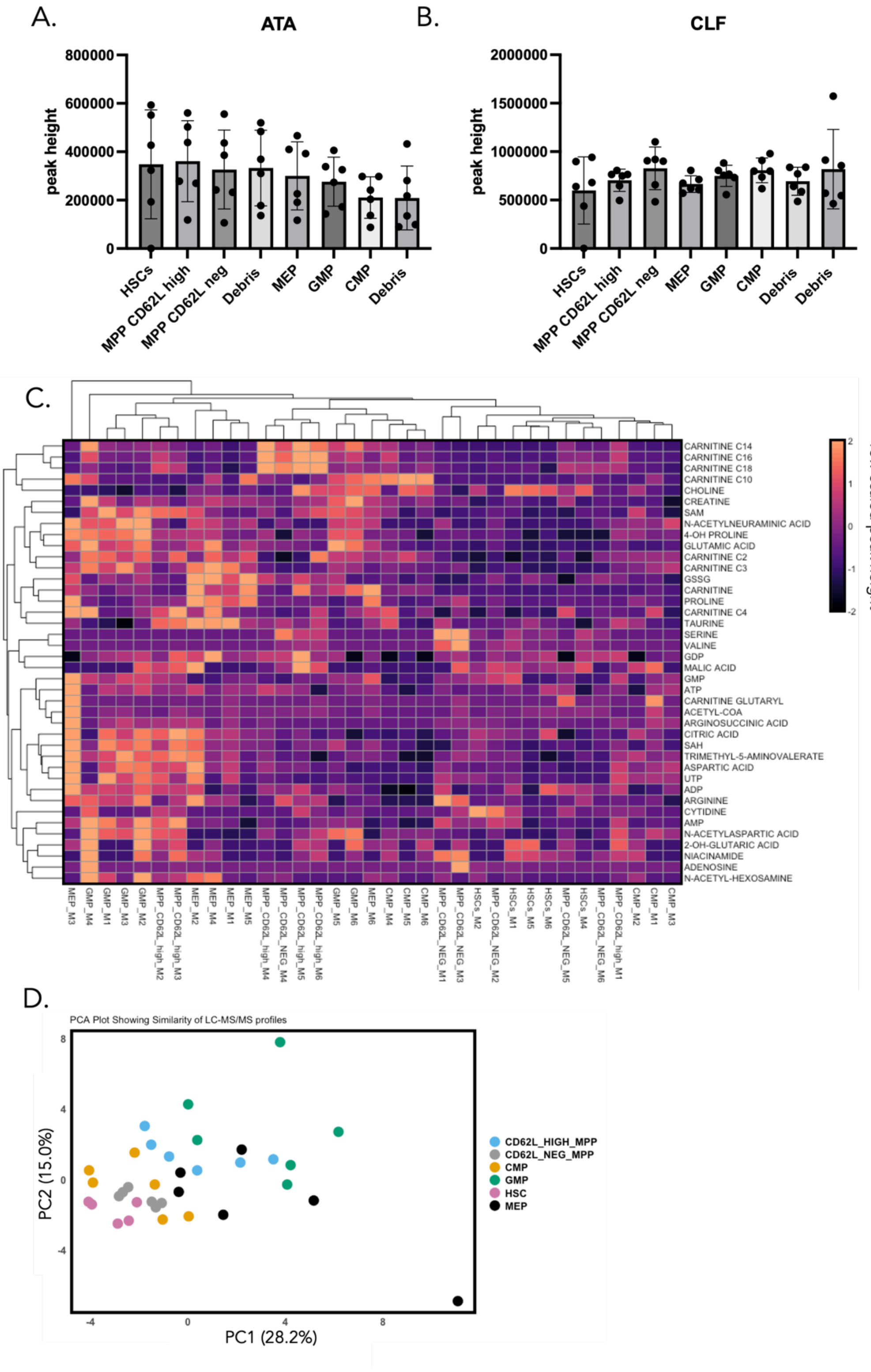

**Figure S16: Supplementary Data Relating to Metabolomics Profiling Experiments** (A) Distribution of the internal standard ATA cell-sorting controls across samples. (B) Distribution of the internal standard CLF cell-sorting controls across samples. For panels A and B no statistical differences between samples were observed, statistical analysis was performed using a Friedmans test to account for the non-normality of the data, and the pairing of cell types within mice. (C) Heatmap showing the median peak height value across all mice, normalised per metabolite for 40 metabolites that were detected across both independent experiments. Columns represent each independent sample that was processed for mass spectrometry profiling. Complete linkage hierarchical clustering based on the Euclidean distances between individual samples. (D) PCA plot computed on all metabolites in the dataset, each point represents a FACS sorted sample (n = 6 mice). ATA: aminoterephthalic acid ; CLF: 4-Chloro-DL-phenylalanine.

A.

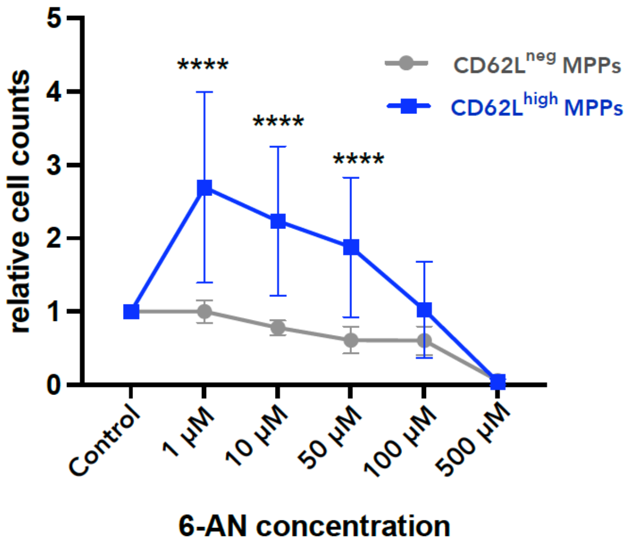

B.

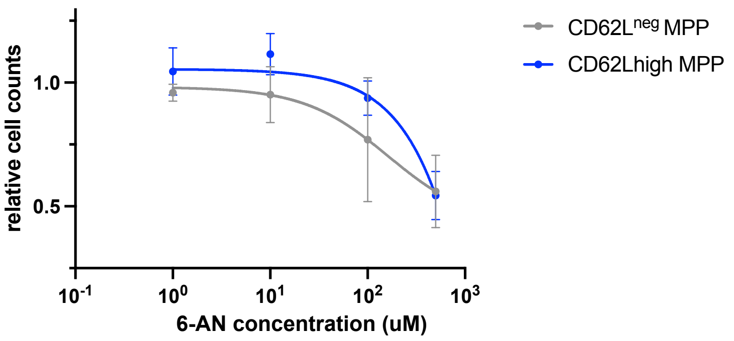

**Figure S17: 6-AN concentration** (A) Differential sensitivity of MPPs to the PPP inhibitor 6-AN computed as relative cell count to the control without 6-AN. In this experiment ckit enriched bone marrow cells were cultured in Stemspan + SCF + TPO for 16 hours in the presence of either 6-AN or control (DMSO) at the concentrations highlighted. Square points represent the mean value, and error bars represent the standard deviation. Statistical significance was tested using a 2-way ANOVA (**** = p value < 0.001). Data were pooled from 2 independent experiments. N = 8 mice. (B) same but starting from purified MPP. Data were pooled from 2 independent experiments. N = 8 mice.

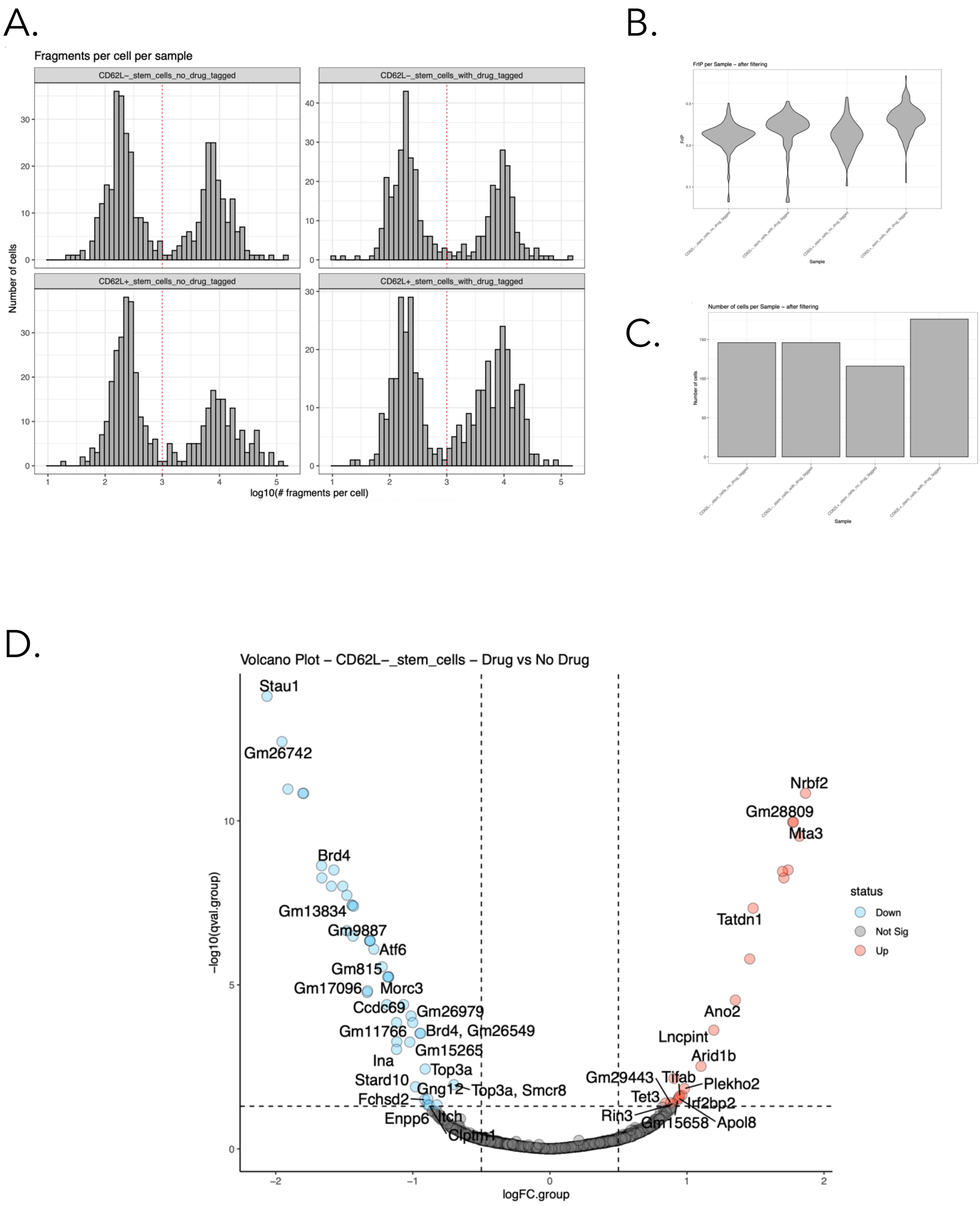

**Figure S18. Data QC and supplementary analyses for scCHICseq data on MPPs treated with 6-AN inhibitor.** (A) Distribution of sequencing depth across cells for each sample, shown as histograms of the log10 number of fragments per cell. The red dotted line indicates the lower threshold applied for cell filtering (1,000 fragments per cell). (B) Distribution of Fraction of Reads in Peaks (FRiP) per cell across samples after quality filtering, displayed as violin plots. (C) Number of cells retained per sample following quality control steps. (D) Volcano plot showing differentially enriched regions between CD62L$^{neg}$ treated and untreated cells. Significantly enriched regions were defined by a q-value < 0.05 and an absolute log2 fold change > 0.5.

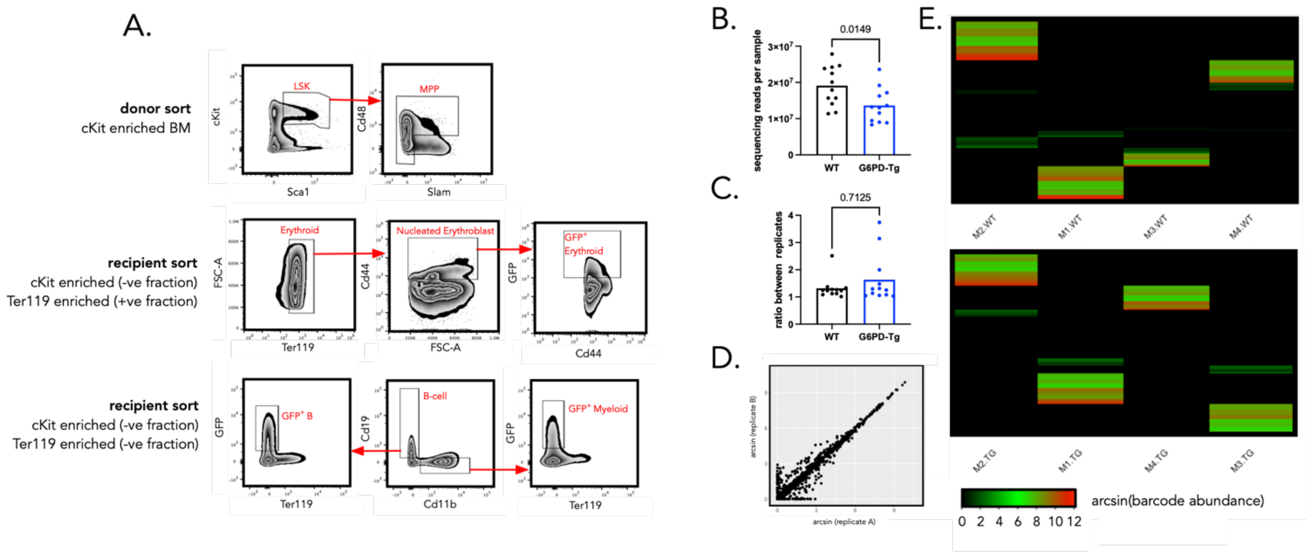

**Figure S19. Data QC for lentiviral barcoding of WT and G6PD-Tg MPPs** (A) Gating strategy to sort MPP for lentiviral barcoding experiments and to sort mature cells after transplantation. (B) The total number of sequencing reads per sample per condition. Each point represents a bulk sorted population of mature cells from one mouse. Normality was assessed using a Shapiro-Wilk test and then pairwise comparisons were made using a Students T-test (C) The ratio of barcode abundances across technical replicates in WT and G6PD-Tg conditions, each point represents a bulk sorted population of cells in one mouse. Normality was assessed using a Shapiro-Wilk test and then pairwise comparisons were made using either a Mann-Whitney test. (D) Representative scatter plot showing the consistency of read counts across technical replicates. Each point represents a unique barcode. Data are transformed using the hyperbolic arcsin function. For WT samples, the Pearson Correlation between technical replicates was of 0.89 +/-0.29 per samples and for the G6PD-tg group 0.98 +/- 0.05. (E) Heatmaps showing the distribution of barcode abundance across mice after data QC and filtering to ensure that we do not have significant numbers of barcodes shared between mice injected with the same batch, these shared barcodes are more likely to label more than 1 cell. Top heatmap represents WT samples, and bottom heatmap represents G6PD-Tg heatmaps. Data are hyperbolic arcsin transformed reads.

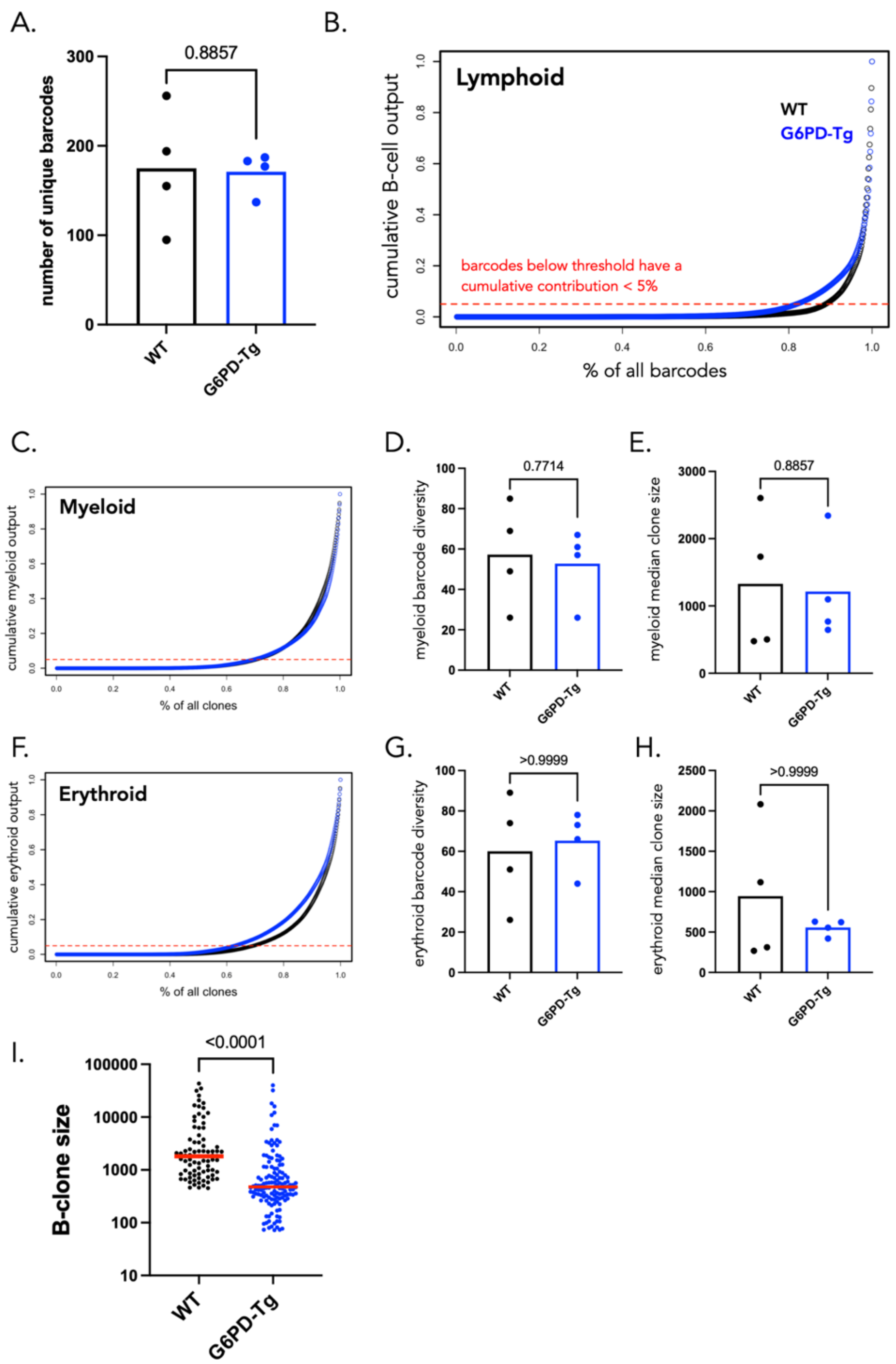

**Figure S20. Supporting analyses for lentiviral barcoding of WT and G6PD-Tg MPPs** (A) The total number of unique barcodes retrieved from WT recipient mice transplanted with WT (black) and G6PD-Tg (blue) MPPs. Each point represents 1 mouse and N = 4 mice per condition. Stats? (B) Cumulative distribution showing the abundance of each barcode in the B-cell lineage. The top *n* barcodes that contribute to 95% of all read counts for the B-lineage are classified as B-cell producing for subsequent analyses. Each point on this plot represents a distinct barcode. (C) Same as B for the Myeloid lineage. (D) The number of unique barcodes producing myeloid cells in WT recipient mice transplanted with WT (black) and G6PD-Tg (blue) MPPs. Statistical comparisons were made using a Mann-Whitney test and each point represents a unique mouse. N = 4 mice (E) The median clone ,size of all myeloid producing barcodes per mouse across WT and G6PD-Tg conditions. Statistical comparisons were made using a Mann-Whitney test and each point represents a unique mouse. N = 4 mice (F) Same as B for the erythroid lineage. (G) The number of unique barcodes producing erythroid cells in WT and G6PD-Tg conditions. Statistical comparisons were made using a Mann-Whitney test and each point represents a unique mouse.. (H) The median clone size of all erythroid producing barcodes per mouse across WT and G6PD-Tg conditions. Statistical comparisons were made using a Mann-Whitney test and each point represents a unique mouse. N = 4 mice (I) Distribution of clone sizes for B-cell producing barcodes across WT and G6PD-Tg conditions. Data pooled from 4 mice per condition. Each point represents a unique lineage barcode (81 WT barcodes, 138 G6PDtg barcodes). Pairwise comparisons were made using a Mann-Whitney test.

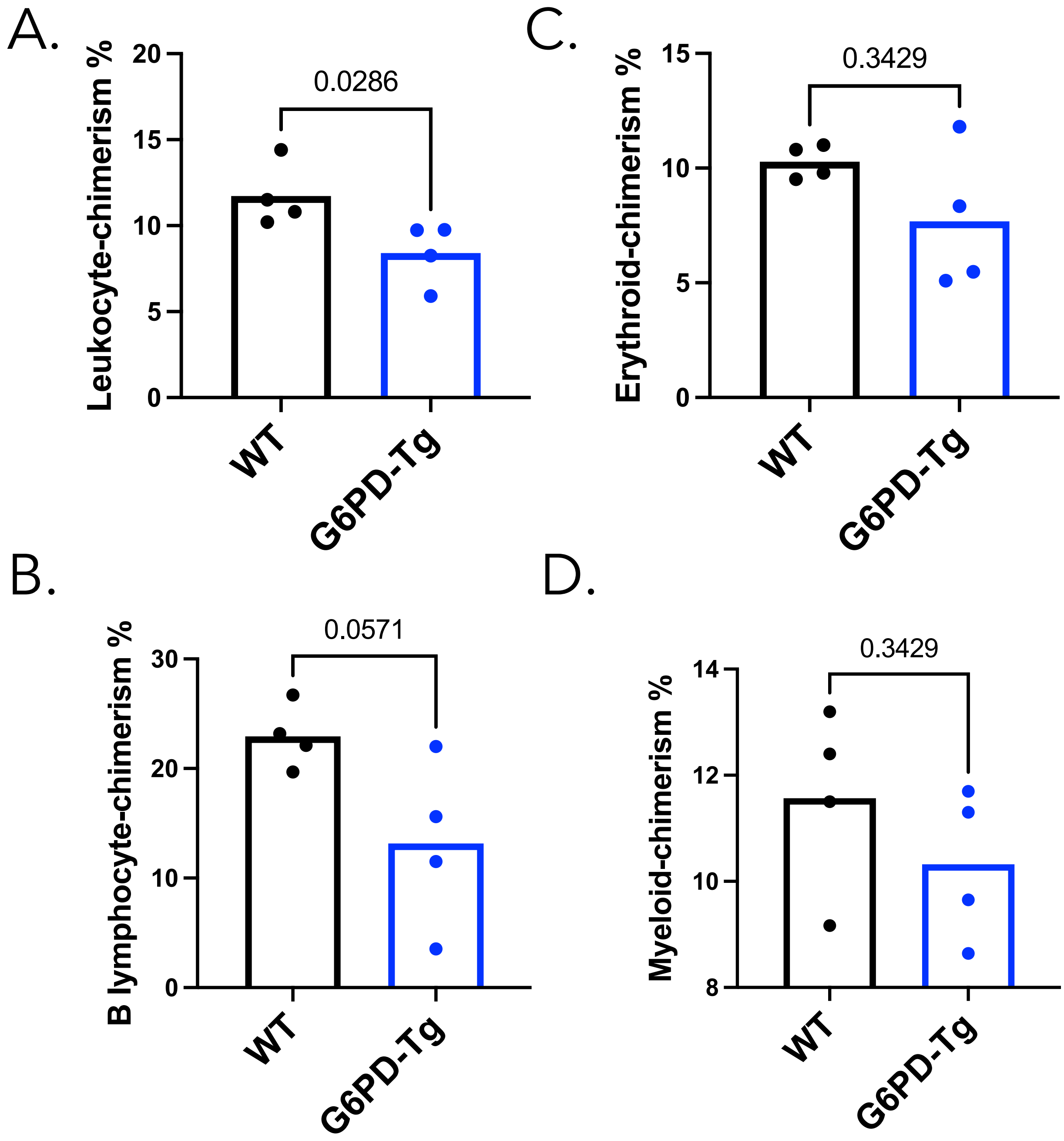

**Figure S21. Bone marrow chimerism analyses for lentiviral barcoding of WT and G6PD-Tg.** (A-D) % chimerism quantified by measuring the proportion of GFP cells relative to total live cell numbers in the respective lineage compartments by flow cytometry for the WT (black) and G6PD-Tg (blue) transplanted MPPs. Each point represents a single mouse with N=4 mice per experimental condition. Pairwise comparisons were made using a Mann-Whitney test.

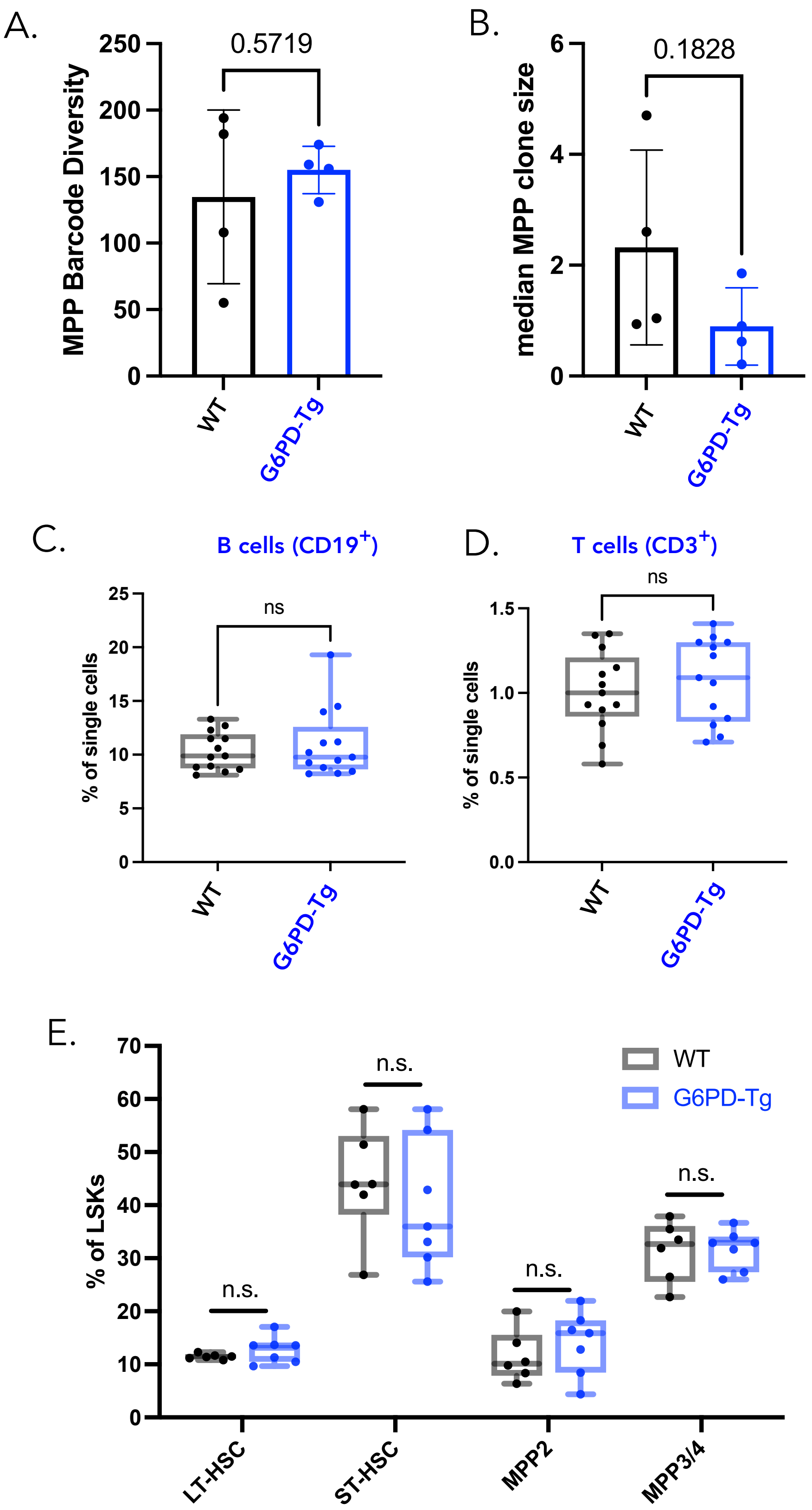

**Figure S22: G6PD Transgene expression does not influence engraftment, viability or division:** (A) Comparison of barcode diversity (number of MPPs that engraft and survive) for transplanted MPPs taken from WT (black) and G6PDTg (blue) mice. Data is reanalysed from Figure 3. Each point represents 1 mouse and n=4 per group. Statistical comparisons were made using a 2-sided unpaired T-test. (B) Comparison of median barcode clone sizes (number of MPP cells with the same barcode) for barcoded MPPs taken from WT (black) and G6PDTg (blue) mice. Data is reanalysed from Figure 3. Each point represents 1 mouse and n=4 per group. Statistical comparisons were made using a 2-sided unpaired T-test.

(C) percentage of CD19⁺ B cells in the single cell gate produced in the host mouse after transplantation of WT (black) and G6PDTg (blue) MPPs. Data from flow cytometry. Each point represents 1 mouse and n=13 per group. Statistical comparisons were made using a 2-sided unpaired T-test. (D) same as C but for CD3⁺ T cells. (E) percentage of HSC and MPP subtypes in LSKs produced in the host mouse after transplantation of WT (black) and G6PDTg (blue) MPPs. Data from flow cytometry. Each point represents 1 mouse and n=6 per group. Statistical comparisons were made using a 2-sided unpaired T-test.

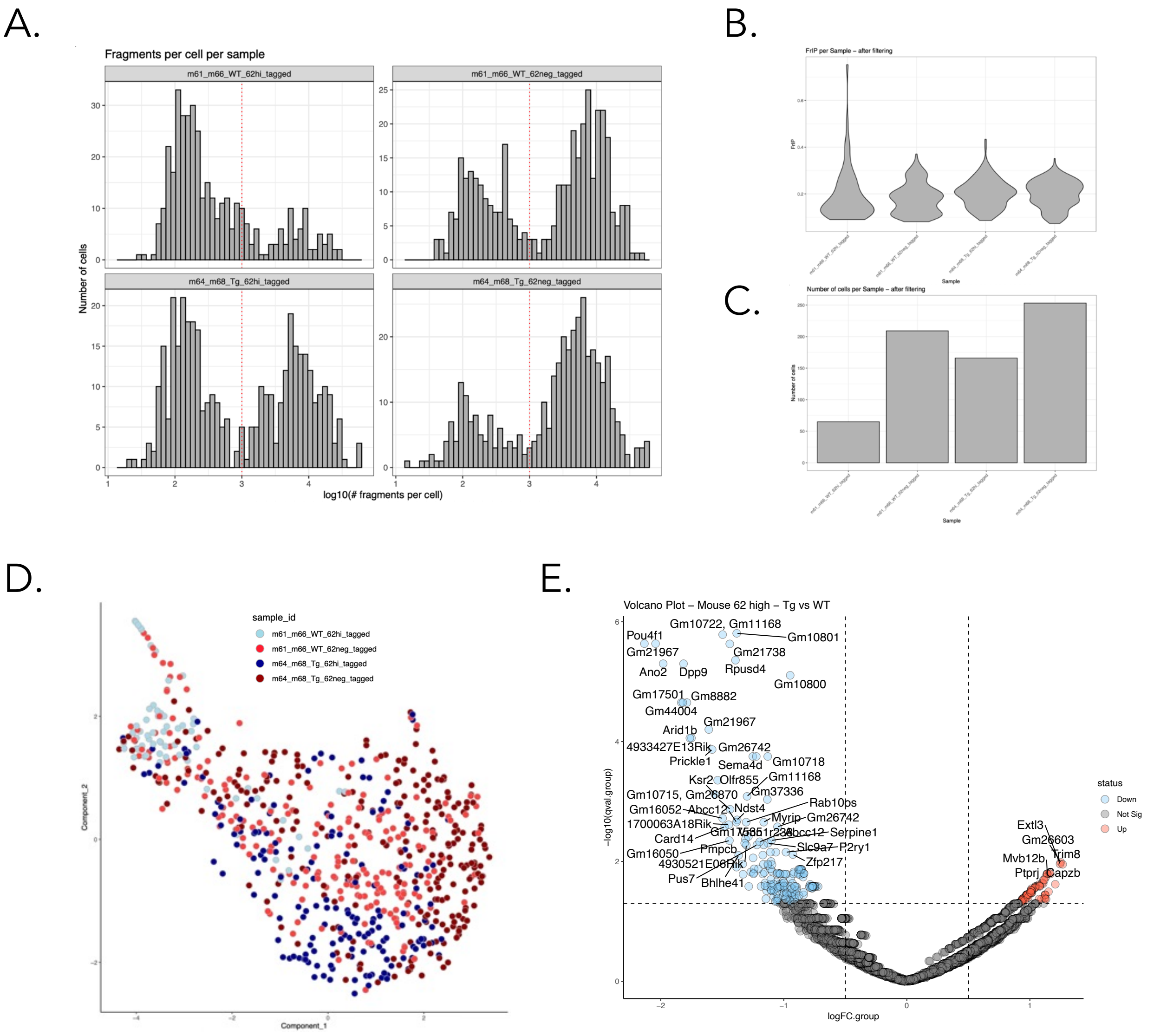

**Figure S23. Data QC and supplementary analyses for scCHICseq data on G6PD Tg and WT MPPs**

A) Distribution of sequencing depth across cells for each sample, shown as histograms of the log10 number of fragments per cell. The red dotted line indicates the lower threshold applied for cell filtering (1,000 fragments per cell). (B) Distribution of Fraction of Reads in Peaks (FRiP) per cell across samples after quality filtering, displayed as violin plots. (C) Number of cells retained per sample following quality control steps. (D) UMAP representation of all cells retained after quality control, colored by sample identity. (E) Volcano plot showing differentially enriched regions between CD62L$^{neg}$ treated and untreated cells. Significantly enriched regions were defined by a q-value < 0.05 and an absolute log2 fold change > 0.5.
